# SARS-CoV-2 ORF9b exploits mitochondrial recruitment to TOMM70 for proteasomal protection and tunes inflammatory remodeling during lung infection

**DOI:** 10.64898/2026.07.13.738231

**Authors:** Yining Zhao, Lauriane Kergoat, Guilherme Dias de Melo, Florence Larrous, Guillaume Drumont, Juan Diego Hernandez Camacho, Elodie Vimont, Elodie le Seac’h, Nell Saunders, Martin Kaechele, Etienne Kornobis, Quentin Giai Gianetto, Mariette Matondo, Isabelle Tardieux, Olivier Schwartz, Hervé Bourhy, Timothy Wai

## Abstract

Mitochondrial outer membrane proteins are exploited by diverse intracellular pathogens, to modulate cell metabolism and innate sensing pathways. Here, we demonstrate that TOMM70-dependent mitochondrial recruitment is required to protect SARS-CoV-2 ORF9b from proteasomal degradation, a dependency conserved across ORF9b homologs from related coronaviruses. ORF9b mitochondrial recruitment requires the E477 residue of TOMM70, a surface distinct from that used by the parasite *Toxoplasma gondii* to engage host mitochondria. We further show that TOMM70 is not a passive scaffold: its depletion activates interferon-stimulated gene expression independently of infection and remodels host immunity distinctly from ORF9b, establishing the receptor and viral protein as mechanistically separable. Using ORF9b-deficient SARS-CoV-2, we demonstrate that ORF9b is dispensable for viral replication and pathological responses in human respiratory epithelial cells and lungs of infected golden Syrian hamsters. Omics profiling of SARS-CoV-2 infected lungs revealed an induction of pathways related to COVID-19 in the absence of ORF9b. Notably, ORF9b-deficient virus-infected lungs show elevated expression of C15ORF48, a nuclear-encoded mitochondrial protein that substitutes for Complex IV subunit NDUFA4 to attenuate inflammation. Collectively, we propose that ORF9b is a receptor-gated viral protein whose principal measurable consequence during authentic infection is a restraint on inflammatory respiratory-chain remodeling.

## Introduction

Mitochondria are multi-functional organelles that execute essential biosynthetic, bioenergetic, and signaling functions that govern the life and death of the cell^1^. Beyond their canonical metabolic roles, mitochondria have emerged as central hubs in host-pathogen interactions, with the outer mitochondrial membrane (OMM) serving as a critical interface that both relays innate immune signals and provides a physical platform that is co-opted by diverse intracellular pathogens^2^. The convergent evolution of viral and parasitic factors that target OMM proteins underscores the strategic importance of this organelle in shaping infection outcomes: obligate intracellular parasites such as *Toxoplasma gondii* exploit the TOMM70 import receptor to establish nutrient-permissive host-parasite contact sites at the parasitophorous vacuole membrane^3–5^, while a growing roster of viral proteins have been reported to engage OMM-resident factors to modulate host cell physiology and subvert antiviral defenses^6^. Among these, SARS-CoV-2 ORF9b has attracted particular attention as a candidate innate immune antagonist and potential mediator of the mitochondrial dysfunction associated with severe COVID-19^7–10^. Whether structurally distinct surfaces on TOMM70 underlie its exploitation by microbes of different kingdoms, and how genuine ORF9b-mediated effects can be discriminated from broader infection-associated remodeling, remain unresolved.

SARS-CoV-2, the beta coronavirus responsible for the COVID-19 pandemic, encodes a repertoire of accessory proteins including ORF3a, ORF6, ORF7a, ORF7b, ORF8, and ORF9b that have been proposed to modulate host cell functions including immune evasion, autophagy, and organelle biogenesis^11^. ORF9b is a 97-amino acid protein encoded within an alternative reading frame overlapping the nucleocapsid (N) gene and expressed as a subgenomic RNA during infection. ORF9b is present in ancestral bat coronaviruses as well as in SARS-CoV-1, previously known as severe acute respiratory syndrome (SARS) coronavirus, responsible for an outbreak^12^ in 2002–2004.

Across SARS-CoV-2 variants, changes in the N/ORF9b region have been associated with altered ORF9b expression, with several variants showing increased subgenomic RNA and protein abundance relative to ancestral strains. This evolution has fueled the notion that ORF9b may act as a molecular rheostat whose tuning has influenced the balance between viral immune evasion and pathogenicity across the pandemic^10^. However, each variant of concern harbors mutations distributed mainly in the spike, but also in other structural and non-structural proteins, it is thus impossible to formally attribute the observed differences in host responses between variants to ORF9b using comparative virology approaches alone.

Global interactome studies identified TOMM70, a TPR domain-containing receptor subunit of the translocase of the outer mitochondrial membrane (TOM) complex that facilitates the chaperone-mediated delivery of nuclear-encoded mitochondrial precursor proteins prior to their translocation through the TOM40 pore, as a prominent and direct physical interactor of ORF9b in human cells^11,13^. Subsequent structural studies characterized the binding interface between ORF9b and the C-terminal helical groove of TOMM70 at atomic resolution^14^, leading to a model in which ORF9b is occupying the hydrophobic client-binding pocket of TOMM70 through its conserved alpha helix, competitively displacing chaperone-delivered precursor proteins and allosterically impairing Hsp90-mediated client delivery, thereby remodeling the mitochondrial proteome^15,16^. Consistent with this model, ORF9b expression reduces interferon production in transfection-based systems and partially phenocopies the proteomic consequences of TOMM70 depletion in HEK293 cells. More recently, a proteomics-based interactome study has proposed that sequence divergence between human and bat sarbecovirus ORF9b proteins underlies differential engagement of distinct OMM receptors, TOMM70 for human coronaviruses and MARC2 for bat-derived sarbecoviruses—which has been interpreted as a molecular signature of host receptor adaptation during zoonotic spillover^17^.

Despite substantial interest in ORF9b as an innate immune antagonist, several questions remain unresolved. First, while structural and biochemical data have implicated the conserved L52 and S53 residues of ORF9b in the TOMM70 binding interface^13,14^, whether disrupting this interaction impacts ORF9b protein stability in living cells has not been established. Second, the amino acid determinants on TOMM70 required for ORF9b recruitment have not been identified *in cellulo*, nor has it been determined whether these same residues mediate engagement of TOMM70 by other intracellular pathogens such as *Toxoplasma gondii*^3,5^, leaving open the question of whether structurally distinct surfaces on a single OMM protein underlie its exploitation by microbes of different kingdoms^2^. Third, the impact of ORF9b recruitment on mitochondrial physiology has not been assessed under conditions that discriminate genuine ORF9b-mediated effects from proteasomal degradation of non-localized viral protein, or transcriptional off-target effects^18^. Finally, and most critically, the contribution of ORF9b to innate immune remodeling and pathogenesis during authentic SARS-CoV-2 infection has not been evaluated in relevant *in vivo* models,

In this study, we set out to resolve these questions using a combination of isogenic, single-copy inducible expression systems, CRISPR-based genetic tools, quantitative omics, and authentic SARS-CoV-2 infection models with WT or ORF9b-deleted viruses in lung epithelial cells and golden Syrian hamsters.

## Results

### ORF9b recruitment to mitochondria safeguards against proteasomal degradation

Throughout the evolution of SARS-CoV-2, Orf9b protein levels have consistently fluctuated, in some cases uncoupled from subgenomic RNA levels^10^, prompting us to study the kinetics of the mitochondrial recruitment and stability of ORF9b in human cells. To do so, we engineered HeLa cells to express wild type ORF9b (Wuhan Strain) tagged with EGFP at the amino terminus (EGFP-ORF9b^WT^) from a single-copy, tetracycline-includible locus using the HeLa Flp-In™ T-REx™ system^19^. This system enables us to finely control the temporal dynamics of ORF9b and to compare variants that could impact protein stability and function. Expression of wild type ORF9b was activated with anhydrotetracycline (AHT)^19^, which can be used at far lower doses than doxycycline or tetracycline, thereby mitigating potential off-target effects caused by mitochondrial toxicity^18^. AHT-induced EGFP-ORF9b^WT^ expression was followed by high-content spinning disc microscopy in HeLa FITR cells in which mitochondria were labeled with Mitotracker DeepRed (MTDR), revealing a time-dependent increase in expression of EGFP-ORF9b signal that overlapped with mitochondria (Figure 1a, b), consistent with previous reports^9,13^. Using this approach, we observed that ORF9b levels increased rapidly in the first few hours after AHT induction and plateaued by 15-24 hours (Figure 1b). These levels did not increase even after an additional week of induction (Figure S1a, b) pointing to post-transcriptional regulation of ORF9b.

**Figure 1:**
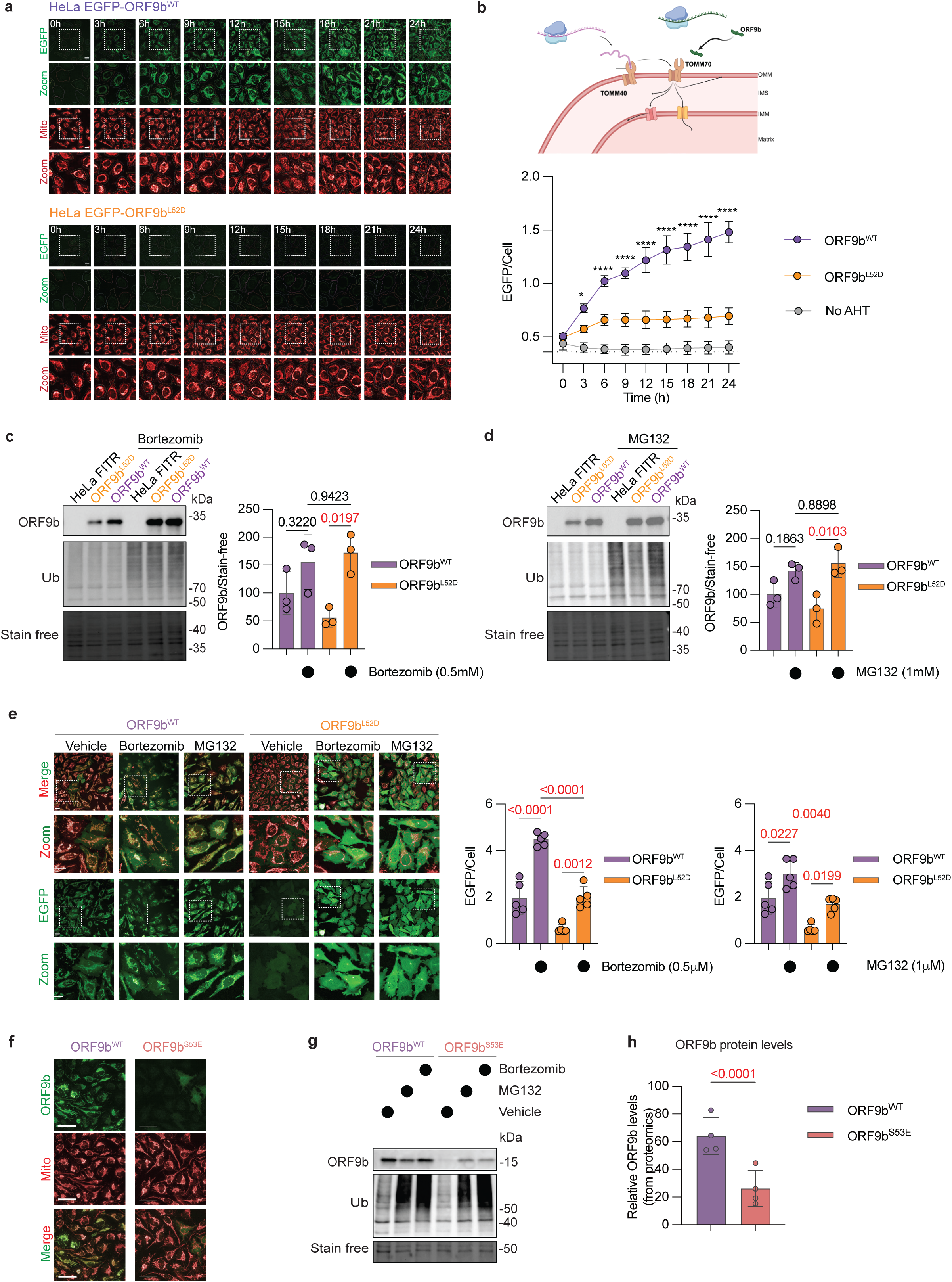
ORF9b recruitment to mitochondria limits its proteasomal degradation. a) Representative images of live imaging of HeLa Flp-In™ T-REx™ (HeLa FITR) cells harboring a stable, single copy of either EGFP-ORF9bWT (top) or mutant EGFP-ORF9bL52D (bottom) whose expressions were induced by anhydrotetracycline (AHT, 10ng/mL). Mitochondria were labeled with Mitotracker DeepRed (MTDR). Dotted white line indicates zoomed area. Scale bar=20 µm. b) (Top) Depiction of cytosolically-translated ORF9b (green) recruitment to TOMM70 on the outer mitochondrial membrane (OMM). Thin arrows depict protein import routes to the OMM, intermembrane space (IMS), inner mitochondrial membrane (IMM), and matrix. (Bottom) Quantification of mean EGFP intensity normalized to nuclei number (EGFP/cell number) using supervised machine learning workflow in Harmony 5.1. HeLa FITR inducibly expressing EGFP-ORF9bWT (ORF9bWT + AHT, purple, n=4), EGFP-ORF9bL52D (ORF9bL52D + AHT, orange, n=4), or uninduced HeLa FITR cells carrying EGFP-ORF9bWT (No AHT, grey, n=5). Dotted line represents basal EGFP signal. Data are means ± SEM of 69-534 cells measured per replicate. 2way ANOVA Šídák’s multiple comparisons test. P-values comparing ORF9bL52D vs ORF9bWT are shown. *P=0.0221, ****P<0.0001. c) Representative western blots of lysates from HeLa FITR induced with AHT to express EGFP-ORF9bWT (ORF9bWT) or mutant EGFP-ORF9bL52D (ORF9bL52D) for 16 hours in presence or absence of proteasomal inhibitor Bortezomib (0.5mM) and probed using the indicated antibodies. HeLa FITR parental cell line not carrying ORF9b (HeLa FITR) was used as a control. Densitometric quantification of EGFP-ORF9bWT (ORF9bWT, purple, n=3) and EGFP-ORF9bL52D (orange, n=3) is relative to stain-free. Data are means ± SEM, one-way ANOVA Tukey’s multiple comparisons test. P-values are indicated and statistically significance is highlighted in red. d) Representative western blots of lysates from HeLa FITR induced with AHT to express EGFP-ORF9bWT (ORF9bWT) or mutant EGFP-ORF9bL52D (ORF9bL52D) for 16 hours in presence or absence of proteasomal inhibitor MG132 (1mM) and probed using the indicated antibodies. HeLa FITR parental cell line not carrying ORF9b (HeLa FITR) was used as a control. Densitometric quantification of EGFP-ORF9bWT (ORF9bWT, purple, n=3) and EGFP-ORF9bL52D (orange, n=3) is relative to stain-free. Data are means ± SEM, one-way ANOVA Tukey’s multiple comparisons test. P-values are indicated and statistically significance is highlighted in red. e) Representative images of live imaging of HeLa FITR induced with AHT to express EGFP-ORF9bWT or mutant EGFP-ORF9bL52D for 12 hours in presence or absence of proteasomal inhibitors Bortezomib (0.5mM) or MG132 (1mM) or vehicle control. Mitochondria were labeled with Mitotracker DeepRed (MTDR). Dotted white line indicates zoomed area. Scale bar=20mm. Single-cell quantification of EGFP intensity (EGFP/Cell) determined with Harmony 5.1. Data are means ± SEM, of 238-1529 cells measured per replicate analyzed by one-way ANOVA Tukey’s multiple comparisons test. P-values are indicated and statistically significance highlighted in red. f) Representative images of HeLa FITR cells induced to express ORF9bWT-FLAG (top) or mutant ORF9bS53E-FLAG (bottom) with anhydrotetracycline (AHT, 10ng/mL) for 24 hours. Indirect immunocytochemistry was performed with anti-TOMM20 to label mitochondria and anti-ORF9b to label ectopic ORF9b. Scale bar=50 µm. g) Western blot of lysates from HeLa FITR induced with AHT to express ORF9bWT-FLAG (ORF9bWT) or mutant ORF9bS53E-FLAG (ORF9bS53E) for 16 hours in presence or absence of proteasomal inhibitors Bortezomib (0.5mM) or MG132 (1mM) or vehicle control using the indicated antibodies. Stain free used as a loading control. h) ORF9b-FLAG levels measured by data independent acquisition (DIA) proteomics of whole-cell lysates from HeLa FITR cells induced to express either ORFb9WT-FLAG (ORF9bWT, purple, n=5) or mutant ORFb9S53E-FLAG (ORF9bS53E, cyan, n=5) for 5 days (Supplemental Dataset 1). Data are means of normalized protein levels (2^data_norm_imp) ± SEM. LIMMA t-test was used and the corresponding p-values were adjusted using an adaptive Benjamini-Hochberg correction and highlighted in red if statistically significant.

Time-lapse imaging revealed that after 9 hours of induction, all the MTDR mitochondrial signal overlapped with EGFP-ORF9b^WT^ yet in some cells, EGFP signal was additionally present in the cytosol (Figure 1a), suggesting a saturation of mitochondrial ORF9b recruitment. To quantify this observation, we devised a supervised machine-learning (SML) based method (see Methods) to classify the various subcellular distributions of ORF9b we observed in four classes: 1) *colocalized*, in which EGFP-ORF9b overlapped exclusively with MTDR, 2) *saturated*, in which EGFP-ORF9b^WT^ overlapped with MTDR and was found diffusely in the cytosol, 3) *weak*, in which EGFP-ORF9b^WT^ signal was low yet still overlapped exclusively with MTDR and 4) *cytosolic*, in which EGFP-ORF9b^WT^ did not overlap specifically with MTDR and was found diffusely in the cytosol (Figure S1c). We quantified time-lapse imaging of EGFP-ORF9b^WT^ cells and confirmed the time-dependent accumulation of ORF9b on mitochondria followed by a delayed increase in the cytosol. We validated our SML workflow by silencing *TOMM70* (si*TOMM70*), which reduced EGFP-ORF9b^WT^ recruitment to mitochondria (Figure S1c, d), consistent with its role as an ORF9b receptor^9^.

To control for potential off-target effects of ORF9b induction in HeLa FITR cells, we generated HeLa FITR cells capable of inducing the expression of mutant L52D (ORF9b^L52D^). This substitution which lies in a region of SARS-CoV-2 ORF9b that is conserved across all SARS-CoV-2 variants of concern as well as ancestral bat coronaviruses and SARS-CoV-1 (Figure S1e)^9,10^, disrupts mitochondrial localization of SARS-CoV-2 ORF9b upon transient transfection^20^ making it an informative negative control. AHT-induced expression of EGFP-ORF9b^L52D^ in HeLa FITR cells was barely detectable by live-cell imaging (Figure 1a,b) and reduced by western blot analyses of whole-cell lysates (Figure 1c) despite mRNA expression at levels that were indistinguishable from EGFP-ORF9b^WT^ (Figure S1f), pointing to post-transcriptional regulation of this mutant. We thus tested whether proteasomal degradation was responsible for the decreased levels of mutant ORF9b^L52D^ and therefore induced EGFP-ORF9b^L52D^ expression in the presence of proteasomal inhibitors Bortezomib or MG132, both of which have previously been used to successfully stabilize mutant, mitochondrially-targeted proteins in cellular models of mitochondrial disease^21^. Proteasomal inhibition increased levels of mutant ORF9b but did not rescue defective mitochondrial recruitment (Figure 1c-e). Notably, we observed similar destabilizing effects of ORF9b with the ORF9b^S53E^ mutation, which alters a phosphorylation site reported to facilitate the interaction of ORF9b and TOMM70 by co-immunoprecipitation^10^. While transient transfection of wild-type and mutant ORF9b^S53E^ constructs were reported to yield comparable steady-state protein levels of ORF9b^9^, analysis of HeLa FITR cells inducibly expressing ORF9b^S53E^-FLAG revealed reduced levels of ORF9b by immunoblot and proteomic analyses (Figure 1g, h) while ORF9b^S53E^ mRNA levels were similar to that of ORF9b^WT^-FLAG control (Figure S1g). Similarly to ORF9b^L52D^, we observed that ORF9b^S53E^ was not recruited to mitochondria (Figure 1f) and was also subject to MG132- and Bortezomib-dependent degradation (Figure 1g). Taken together, our data suggest that mutations in conserved residues that disrupt the ability of ORF9b to be recruited to the OMM via TOMM70 promote proteasomal degradation of the viral protein.

### Equivalent mitochondrial recruitment kinetics of ORF9b from human and bat coronaviruses

To determine whether the mitochondrial recruitment and stability of SARS-CoV-2 ORF9b are conserved across related human and bat sarbecoviruses, we generated HeLa FITR cell lines expressing EGFP-tagged ORF9b variants from SARS-CoV-1, BANAL-236 (Ban Na Lao no. 236) and RaTG13 (*Rhinolophus affinis* bat coronavirus sample TG collected in 2013) from the same single-copy, AHT-inducible locus (Figure 2a). SARS-CoV-1 is the etiologic agent of severe acute respiratory syndrome (SARS) in humans, a highly transmissible pneumonia-like disease that caused a global outbreak in 2002–2003^12^. BANAL-236 and RaTG13 are horseshoe bat–derived sarbecoviruses that were discovered in Northern Laos and Southwestern China, respectively. They are closely related at the genome level to SARS-CoV-2 but have only been detected in bats to date, where they circulate as a natural coronavirus reservoir. BANAL-236, is most closely related virus to SARS-CoV-2 ^22^, while RaTG13, discovered in 2013 is more distant^23^. Sequence alignments revealed the greatest sequence divergence between ORF9b from RaTG13 and BANAL-236 showed 93% and 98% sequence conservation, respectively (Figure 2a).

**Figure 2:**
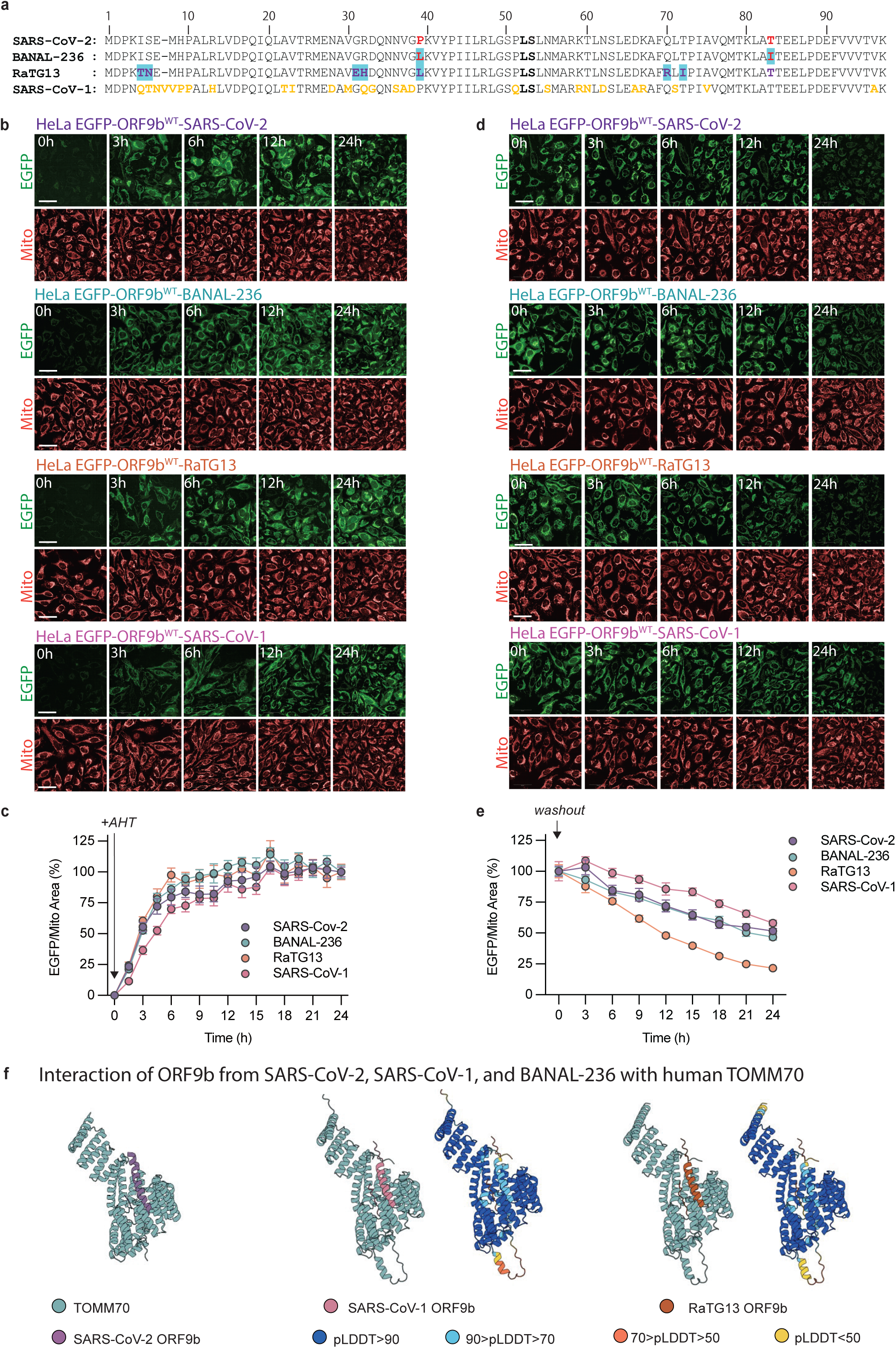
TOMM70-dependent recruitment of ORF9b from divergent sarbecoviruses. a) Amino-acid sequences of the ORF9b accessory protein from SARS-CoV-2, bat coronavirus RaTG13,bat sarbecovirus BANAL-236, and SARS-CoV-1, were aligned to highlight conserved and divergent (colored) residues. L52 and S53 residues are in bold. ORF9b sequences correspond to the translated coding region of each virus’s subgenomic ORF9b. SARS-CoV-2 and SARS-CoV-1 ORF9b protein sequences were obtained from UniProtKB (SARS-CoV-2 ORF9b, UniProt accession P0DTD2; SARS-CoV-1 ORF9b, UniProt accession P59636). RaTG13 ORF9b sequence was derived by translating the ORF9b coding region from the complete RaTG13 genome sequence (GenBank accession MN996532.2) obtained from NCBI GenBank. BANAL-236 ORF9b sequence was derived by translating the ORF9b coding region from the complete genome of bat sarbecovirus BANAL-20-236 (GenBank accession MZ937003; corresponding GISAID accession EPI_ISL_4302647)^22^. Sequence numbering refers to the ORF9b protein N-terminus. b) Representative images of live imaging of HeLa Flp-In™ T-REx™ (HeLa FITR) cells expressing EGFP-ORF9b variants from SARS-CoV-2 (purple), BANAL-236 (pink), RaTG13 (orange), and SARS-CoV-1 (teal) with anhydrotetracycline (AHT, 10ng/mL) for indicated time points. Mitochondria were labeled with Mitotracker DeepRed (MTDR). Scale bar=20 µm. c) Quantification EGFP-ORF9b variants from SARS-CoV-2 (purple, n=7), BANAL-236 (teal, n=7), RaTG13 (orange, n=7), and SARS-CoV-1 (pink, n=7) recruitment to mitochondria (MTDR) using Harmony 5.1 as integrated EGFP/MTDR every 1.5h for 24 hours induction with anhydrotetracycline (AHT, 10ng/mL). For each cell line, values were normalized to its own time course, with 0 h after AHT addition set as baseline and 24 h set as maximum (100%). Data are presented as mean ± SEM of 7 replicates with 426–1629 cells measured per replicate. d) Representative images of live imaging of HeLa Flp-In™ T-REx™ (HeLa FITR) cells expressing EGFP-ORF9b variants from SARS-CoV-2 (purple, n=7), BANAL-236 (pink, n=7), RaTG13 (orange, n=7), and SARS-CoV-1 (teal, n=7) after washout of anhydrotetracycline (AHT, 10ng/mL) in Figure 2c. Time indicates hours of washout following 24 h of AHT incubation. Mitochondria were labeled with Mitotracker DeepRed (MTDR). Scale bar=20 µm. e) Quantification EGFP-ORF9b expression after AHT washout. Cells induced with anhydrotetracycline (AHT, 10ng/mL) for 24 h in Figure 2b were washed out of AHT (washout, t = 0 h). EGFP was quantified using Harmony 5.1 and integrated EGFP intensity normalized to mitochondrial area. For each cell line, EGFP values were normalized to the signal at the time of AHT washout, which was set to 100%. Data are shown as the percentage of EGFP remaining over time (means ± SEM of 8 replicates with 376-1481 cells measured per replicate). f) AlphaFold 3.0–predicted structural models of human TOMM70 in complex with ORF9b from SARS-CoV-2, SARS-CoV-1, and BANAL-236 are shown. Human TOMM70 is shown in cyan, while ORF9b proteins are colored as follows: SARS-CoV-2 ORF9b (purple), SARS-CoV-1 ORF9b (pink), and BANAL-236 ORF9b (orange). Per-residue predicted confidence (pIDDT) is represented by the indicated color scale. All structures were visualized using PyMOL.

Expression of each of the four EGFP-ORF9b variants was monitored in HeLa FITR cells (Figure 2b). Quantitative comparison of mitochondrial localization over time revealed similar recruitment kinetics to mitochondria over 24 hours (Figure 2c, Supplemental Table 1). Following AHT washout, mitochondrially associated ORF9b from SARS-CoV-2, SARS-CoV-1 and BANAL-236 decayed with similar turnover kinetics, with no detectable differences in the rate of loss of the mitochondrial pool (Figure 2d, e, Supplemental Table 2). We observed an accelerated disappearance of EGFP signal in HeLa FITR cells expressing ORF9b from RaTG13 relative to all other ORF9b variants from 18h onwards, suggesting that this variant may be slightly more unstable. Thus, the rapid mitochondrial recruitment of ORF9b is a conserved property across these human and bat sarbecoviruses.

Because a recent proteomics-based interactome study by the Krogan lab proposed that sequence divergence between bat and human ORF9b dictates preferential engagement of distinct mitochondrial receptors—TOMM70 for human ORF9b and MARC2 for bat ORF9b^17^—we tested the contribution these two unrelated OMM proteins might have to recruitment kinetics. The only two amino acid residue differences between SARS-CoV-2 ORF9b and BANAL-236 ORF9b are at position 39 (Phenylalanine in SARS-CoV-2 and Leucine in BANAL-236) and at position 72 (Threonine in SARS-CoV-2 and Isoleucine in BANAL-236), the latter of which has been proposed as an explanation for the differential, species-specific binding affinity of ORF9b variants to TOMM70^17^. We depleted *TOMM70* or *MARC2* by siRNA in HeLa FITR cells 48h prior to inducing the expression of EGFP-ORF9b variants and then quantified mitochondrial recruitment of EGFP-OR9b. Consistent with our initial validation (Figure S1c, d), *TOMM70* knockdown markedly diminished mitochondrial recruitment for all four ORF9b homologs, reducing the mitochondrially-localized ORF9b (Figure S2a, b) to similar levels as observed previously (Figure S1c). In contrast, *MARC2* silencing had no measurable effect on the mitochondrial localization or stability for any ORF9b variant despite comparable knockdown efficiencies (Figure S2c), indicating that it is dispensable for ORF9b recruitment in human cells. We used AlphaFold 3.0 to model complexes between human TOMM70 and ORF9b from either BANAL-236 or SARS-CoV-2, in order to assess whether the amino-acid differences between these variants might alter the TOMM70–ORF9b interface and thus the propensity for binding. Consistent with our *in cellulo* data, the structures predicted same human TOMM70 binding sites for the other ORF9b variants (Figure 2f). Taken together, these data indicate that amino acid differences that distinguish SARS-CoV-2 ORF9b from SARS-CoV-1 and its closest known bat coronavirus relative are dispensable for TOMM70-dependent recruitment to human mitochondria and argue against a switch from TOMM70 to MARC2 as the primary mitochondrial receptor during adaptation for zoonotic potential.

### SARS-CoV-2 ORF9b mitochondrial recruitment requires the E477 residue on TOMM70

Having excluded that human-adapted sequence variants in sarbecovirus ORF9b variants impact the recruitment of ORF9b to mitochondria, we turned our attention back to the study of ORF9b mutants that disrupt the mitochondrial recruitment to better understand the underlying molecular mechanism (Figure 1a, f). The L52D and S53E mutations of SARS-CoV-2 ORF9b are situated in an alpha helix of ORF9b that interfaces with TOMM70 (Figure 3a) but these mutations have also been hypothesized to modulate ORF9b structure^9,24^, which raises the possibility that L52D and S53E mutations may promote rapid degradation by intrinsically destabilizing ORF9b before being recruited to the OMM. Alternatively, it is equally plausible that the conserved region harboring L52 and S53 is required to mediate the binding of ORF9b to TOMM70, without which the viral protein is rapidly degraded by the proteasome (Figure 1c-g). To distinguish between these two models, we deleted *TOMM70* in HeLa FITR cells able to inducibly express wild type ORF9b (either ORF9b^WT^-FLAG or EGFP-ORF9b^WT^) by CRISPR-based genome editing. We identified clones by western blot and characterized the loss-of-function alleles by deep next generation sequencing (NGS) of PCR amplicons covering the targeted region (Figure S3a-c), which revealed loss of function mutations in *TOMM70* CRISPR clones (*TOMM70^C^*^11^, *TOMM70^C^*^14^, and *TOMM70^C^*^15^) that were absent in the parental, wild type HeLa FITR cells.

**Figure 3:**
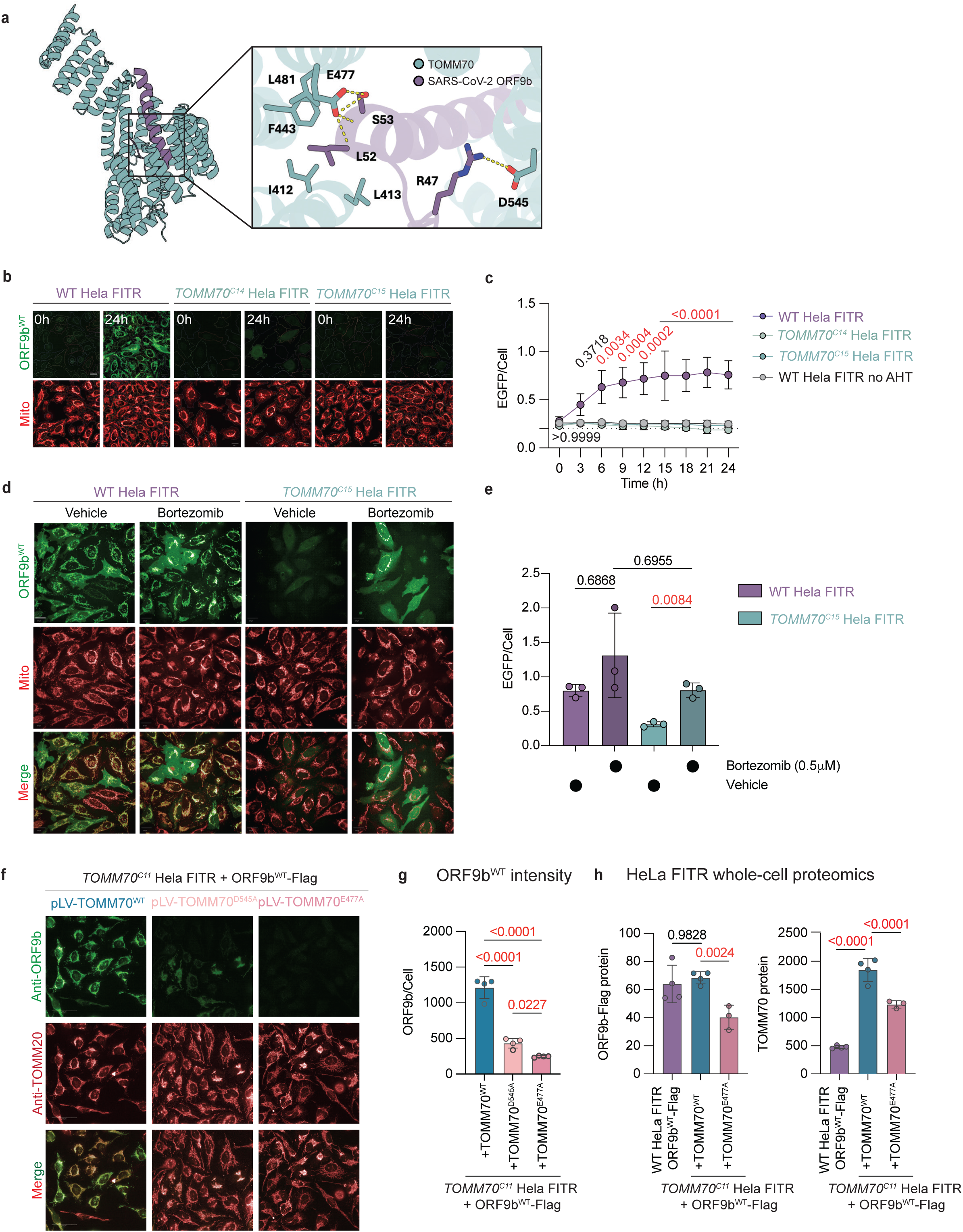
ORF9b mitochondrial recruitment requires the E477 residue on TOMM70. a) Structural representation of SARS-CoV-2 ORF9b (purple) and TOMM70 (green) binding with Alphafold 3.0 and visualized using PyMOL. Interactions interfaces with TOMM70 including the L52 and S53 residues of ORF9b are indicated in the inset. b) Live-cell imaging of wild type HeLa FITR cells and *TOMM70^C^*^14^ and *TOMM70^C^*^15^ HeLa FITR cells induced to express EGFP-ORFb9^WT^ with anhydrotetracycline (AHT, 10ng/mL) for 24 hours. Representative images at 0 and 24 hours following AHT addition. Mitochondria (Mito) were labeled with Mitotracker DeepRed. Cell segmentation with Harmony 5.1 Scale bar=20 µm. c) Quantification of mean EGFP intensity normalized to nuclei number (EGFP/cell number) using Harmony 5.1. Wild type (WT) HeLa FITR cells (WT HeLa FITR, n=3, purple) and TOMM70-deficient HeLa FITR clones 14 (*TOMM70^C^*^14^, n=3, green) and 14 (*TOMM70^C^*^15^, n=3, green) inducibly expressing EGFP-ORF9b^WT^. Uninduced WT HeLa FITR cells carrying the EGFP-ORF9b^WT^ transgene (No AHT, n=3, grey). Dotted line represents background EGFP signal. Data are means ± SEM of 3 replicates with 388-1238 cells measured per replicate. 2way ANOVA Šídák’s multiple comparisons test. P-values comparing ORF9b^L52D^ vs ORF9b^WT^ are shown and highlighted in red if statistically significant. d) Representative images of wild type (WT) HeLa FITR cells and *TOMM70^C^*^15^ HeLa FITR cells induced to express EGFP-ORFb9^WT^ (green) with anhydrotetracycline (AHT, 10ng/mL) for 12 hours in the presence Bortezomib (0.5mM) or vehicle control. Mitochondria were labeled with Mitotracker DeepRed (MTDR, red). Scale bar=20 µm. e) Quantification of mean EGFP intensity normalized to nuclei number in Figure 3d). Data are means ± SEM of 3 replicates with 235-1159 cells measured per replicate. 2way ANOVA Šídák’s multiple comparisons test. P-values comparing ORF9b^L52D^ vs ORF9b^WT^ are shown and highlighted in red if statistically significant. f) Representative images of *TOMM70^C^*^15^ HeLa FITR cells and *TOMM70^C^*^15^ HeLa FITR cells stably expressing wild type (pLV-TOMM70^WT^) or mutant TOMM70 (pLV-TOMM70^E477A^ or pLV-TOMM70^D545A^). ORF9b^WT^-FLAG expression was induced by anhydrotetracycline (AHT, 10ng/mL) immunocytochemistry was performed with against TOMM20 (mitochondria) and ORF9b. Scale bar=20 µm. g) Quantification of mean anti-ORF9b intensity normalized to nuclei number (ORF9b/Cell) in *TOMM70^C^*^15^ HeLa FITR cells stably expressing wild type (pLV-TOMM70^WT^, blue, n=4) or mutant TOMM70 (pLV-TOMM70^E477A^, light pink, n=4 and pLV-TOMM70^D545A^, dark pink, n=4) in Figure 3f. Data are means ± SEM, one-way ANOVA of 4 replicates with 428-1209 cells were imaged for each replicate. P-values are shown and highlighted in red if statistically significant. h) ORF9b-FLAG and TOMM70 levels measured by data independent acquisition (DIA) proteomics of whole-cell lysates from WT HeLa FITR cells (purple, n=4), *TOMM70^C^*^15^ HeLa FITR cells stably expressing wild type (pLV-TOMM70^WT^, blue, n=4) or mutant TOMM70 (pLV-TOMM70^E477A^, light pink, n=4). Data are means of normalized protein levels (2^data_norm_imp) ± SEM. LIMMA t-test was used and the corresponding p-values were adjusted using an adaptive Benjamini-Hochberg correction and highlighted in red if statistically significant.

In line with *TOMM70* knockdown studies (Figure S1c, d), AHT-induced expression of EGFP-ORF9b^WT^ in two independent TOMM70-deficient clones (HeLa FITR *TOMM70^C^*^14^ and HeLa FITR *TOMM70^C^*^15^) suppressed the levels of EGFP detectable by live imaging (Figure 3b, c). Similar effects of TOMM70-dependent ORF9b destabilization were observed in ORF9b^WT^-FLAG-expressing HeLa FITR *TOMM70^C^*^11^ cells (Figure S3d). As these changes were not a consequence of ORF9b transcript levels (Figure S3e), we sought to test whether proteasomal inhibition prevents ORF9b degradation in TOMM70-deficient cells. We induced EGFP-ORF9b^WT^ expression with AHT in bortezomib-treated HeLa FITR *TOMM70^C^*^15^ cells and performed live-cell imaging, which revealed that bortezomib rescued EGFP-ORF9b^WT^ levels but not mitochondrial recruitment (Figure 3d, e), phenocopying the proteasomal-dependent ORF9b destabilization we observed with TOMM70-replete HeLa FITR cells expressing mutant ORF9b^L52D^ (Figure 1d, e). Taken together, our data argue that mitochondrial recruitment of ORF9b by TOMM70 safeguards against proteasomal degradation of both the wild type and mutant viral protein.

Next, we sought to identify the amino acid residues of TOMM70 required to mediate this interaction. In silico modeling based on the published structure of ORF9b and truncated TOMM70 highlighted the importance of the E477 and D545 residues, which are both conserved amino acid residues that lie within the C3 terminal TPR motifs of the protein (Figure 3a). Combining these observations with previous *in vitro* assays measuring the thermal stability of recombinant, truncated TOMM70 and mutant ORF9b^15^, one would predict that exchanging an alanine at 477 for the glutamic acid residue (E477A) in TOMM70, which interfaces with the L52 and S53 region of ORF9b would more profoundly disrupt ORF9b stability and recruitment than the substitution of alanine for aspartate at a more distantly positioned 545 residue (D545A) of TOMM70. We directly tested this prediction by stably expressing full-length TOMM70 mutants E477A (pLV-TOMM70^E477A^) or D545A (pLV-TOMM70^D545A^) as well as wild type TOMM70 (pLV-TOMM70^WT^) in *TOMM70^C^*^11^ HeLa FITR cells capable of induced expression of ORF9b^WT^-FLAG, assessing ORF9b recruitment and expression by indirect immunofluorescence (Figure 3f). In line, we observed that ORF9b^WT^-FLAG was virtually absent in pLV-TOMM70^E477A^-expressing cells while pLV-TOMM70^D545A^-expressing *TOMM70^C^*^11^ HeLa FITR cells yielded a modest level of ORF9b expression and mitochondrial recruitment, albeit well below levels obtained through functional complementation of *TOMM70^C^*^11^ HeLa FITR cells with wild type pLV-TOMM70 or levels observed in WT HeLa FITR cells (Figure 3g, h). Data independent acquisition (DIA) proteomics revealed a significant reduction in ORF9b^WT^-FLAG levels in whole-cell lysates of HeLa FITR *TOMM70^C^*^11^ cells functionally complemented with pLV-TOMM70^E477A^ relative to pLV-TOMM70^WT^. The observed reduction in ORF9b mirrored that seen in TOMM70-replete cells expressing the mutant ORF9b^S53E^-FLAG (Figure 3i, Supplemental Dataset 1). Western blot analyses revealed TOMM70^E477A^ and TOMM70^D545A^ mutant proteins were expressed in *TOMM70^C^*^11^ HeLa FITR cells at similar levels to ectopically, stably-expressed wild type TOMM70^WT^ (Figure S3f). Quantification of ORF9b via proteomics showed that TOMM70^E477A^ re-expression was lower than of TOMM70^WT^, although still substantially higher than WT HeLa FITR parental cells (Figure 3H), ruling out the possibility that diminished mitochondrial recruitment of ORF9b by mutant TOMM70 protein results from insufficient expression. Taken together, we conclude that the E477A mutation in TOMM70 compromises the binding capacity for ORF9b recruitment, rendering it ineffective as a viral protein receptor.

### The E477 residue on TOMM70 is dispensable for parasite recruitment to mitochondria

Having established that SARS-CoV-2 ORF9b stability and recruitment to mitochondria is dependent on the glutamic acid residue at position 477 of TOMM70, we sought to determine whether it was also required for other obligate intracellular microbes known to physically and functionally interact with the TOMM70 protein. One such microbe is the protozoan parasite *Toxoplasma gondii,* which actively invades about any nucleated cells of warm-blooded hosts and subsequently resides within a growth permissive parasitophorous vacuole (PV)^25^. *T. gondii* co-opts host mitochondria via a direct association of products localized to the parasitophorous vacuole membrane (PVM) and TOMM70 at the OMM, thereby establishing intimate host-parasite contact sites that are thought important for nutrient exchange and immune evasion^5^. These parasite-derived products have been identified as members of the mitochondrial association factor one (TgMAF1) paralog repertoire, and display strain-specific restriction^3,26^.

We first confirmed that TOMM70 deficiency impairs mitochondrial recruitment in *T. gondii*–infected HeLa FITR *TOMM70^C^*^11^ mutant cells using live-cell imaging. Intracellular replicating parasites which expressed cytosolic GFP were visualized together with host mitochondria labeled with MTDR. Consistent with previous reports^5^, loss of TOMM70 markedly reduced mitochondrial association with the PV when compared to endogenous TOMM70 expression (Figure 4a). Using the same experimental setting, we next examined whether mitochondrial recruitment could be functionally rescued in HeLa FITR *TOMM70^C^*^11^ cells stably expressing either wild-type TOMM70 (pLV-TOMM70^WT^) or the E477A mutant (pLV-TOMM70^E477A^). In both complemented cell lines, elongated mitochondria were observed to typically align with the PV surface (Figure 4a). Quantitative image analysis indicates no significant difference in the extent of mitochondrial association between WT cells or cells expressing pLV-TOMM70^WT^ and pLV-TOMM70^E477A^ (Figure 4b). These data highlight that in contrast to the SARS-CoV-2 virus, the E477 residue is dispensable for TOMM70-dependent mitochondrial targeting by *T. gondii*.

**Figure 4:**
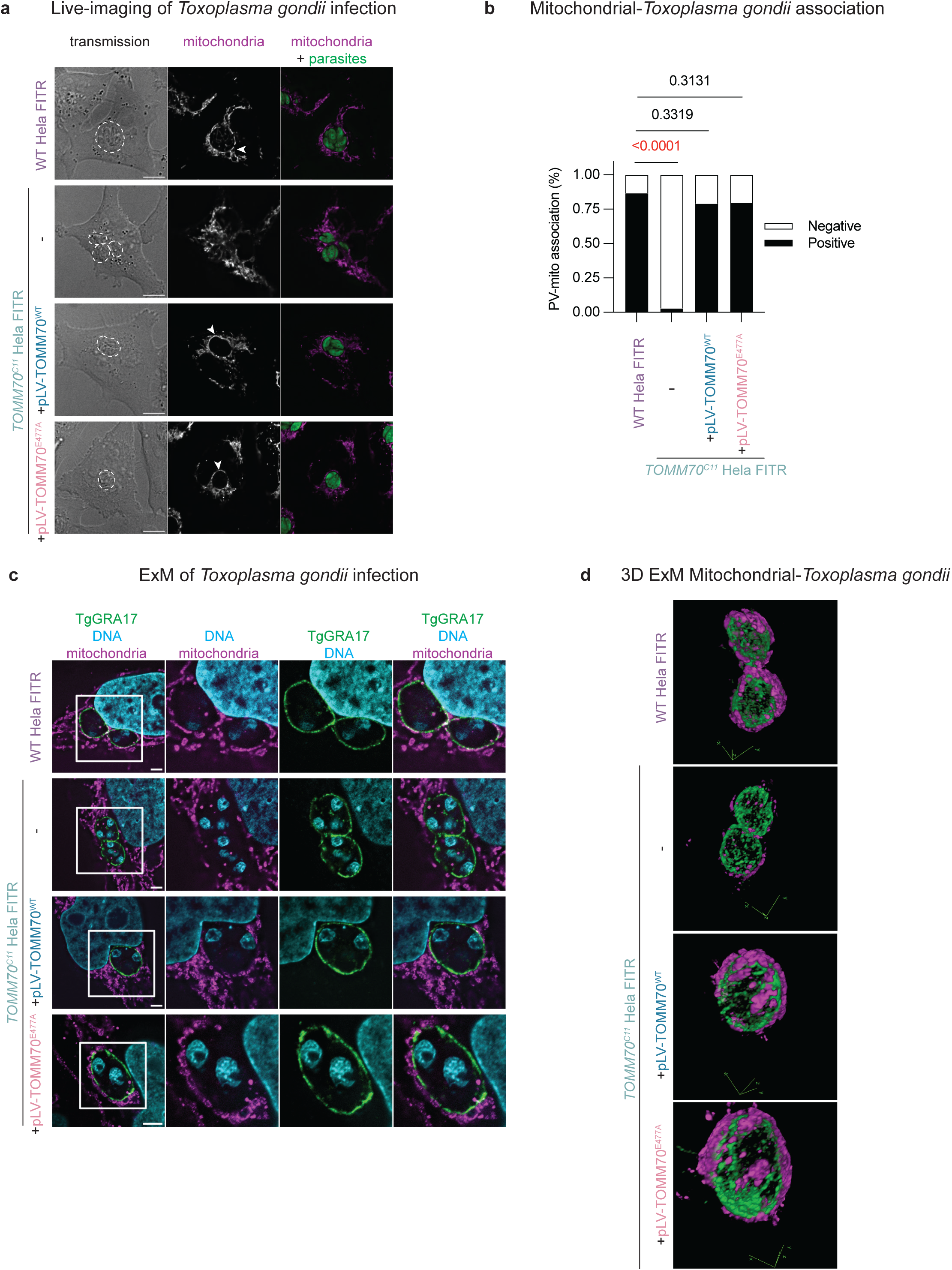
Mitochondrial-parasite interactions in *Toxoplasma*-infected HeLa cells. a) Represented images from live imaging of mitochondria labeled with MitoTracker DeepRed (MTDR, magenta) in Wild type (WT) HeLa FITR cells or TOMM70C11 HeLa FITR cells in which wild type (+pLV-TOMM70WT) or E477A mutant (+pLV-TOMM70E477A) was stably expressed. Images were acquired 18 to 20 h post infection with GFP-expressing T. gondii tachyzoites (green). All scale bars indicated 10 mM. White arrowheads point to the mitochondria juxtaposed to the parasite-containing vacuoles. b) Histogram showing the proportion of vacuoles with and without mitochondrial juxtaposition in Figure 4a. Statistical analysis was performed using a chi-squared (χ²) test; p-values are indicated and the number of PV analyzed is indicated in parentheses below the x axis. c) Expansion microscopy (ExM) images of immunolabeled samples for mitochondria (TOMM20, magenta) and the tachyzoite dense granule GRA17 protein (TgGRA17, green) prior to gelation and expansion. Host cell and parasite nuclei (DNA, blue) are stained post-expansion with Propidium iodide. In contrast to TOMM70C11 HeLa FITR cells, mitochondria are tightly aligned with the TgGRA17 delimited parasitophorous vacuole membrane (PVM) in wildtype Wild type (WT) HeLa FITR cells and TOMM70C11 HeLa FITR cells in which wild type (+pLV-TOMM70WT) or E477A mutant (+pLV-TOMM70E477A) was stably expressed. All scale bars indicated 4 mM (expansion factor x4). d) Representative plane from Figure 4c derived from 3D reconstitution (see Supplemental Movie 1) demonstrating tight, spatial alignment of mitochondria (TOMM20, magenta) TgGRA17 delimited PVM (green) in wildtype Wild type (WT) HeLa FITR cells and TOMM70C11 HeLa FITR cells in which wild type (+pLV-TOMM70WT) or E477A mutant (+pLV-TOMM70E477A) was stably expressed.

To further corroborate these observations with higher spatial resolution, we visualized host mitochondrial proximity relative to the *T. gondii* vacuole using expansion microscopy (ExM). For this purpose, we utilized transgenic *T. gondii* expressing a HA epitope tagged version of TgGRA17, a cytoplasmic dense granule protein (GRA), that is secreted upon exocytosis and targets the PVM where it contributes to pore formation^27^. Host cell mitochondria were immunolabelled with antibodies against the OMM protein TOMM20 marker. Consistent with the live-cell imaging data, ExM studies showed that mitochondria failed to juxtapose with the PVM in HeLa FITR *TOMM70^C^*^11^ cells (Figure 4c). In contrast, complementation with pLV-TOMM70^WT^ and pLV-TOMM70^E477A^ restored a typical tight alignment with the GRA17-positive PVs (Figure 4d, Supplemental Movie 1). Together, these results reveal for the first time that distinct TOMM70 interaction determinants are required for viral (ORF9b) versus *T. gondii* engagement of host mitochondria.

### Disruption of ORF9b recruitment by *TOMM70^E^*^477^*^A^* preserves canonical functions of TOMM70

Human genetic studies have revealed that pathogenic variants in *TOMM70*, including in the conserved C3 terminal tail region in which E477 and D545 are located, can lead to protein destabilization and loss-of-function^28,29^. While polymorphisms have been reported at D545, polymorphisms corresponding to an alternative E477 allele in Exon 9 have been reported with a minor allele frequency (MAF) that is well below 0.01 in the general population (Figure S4a), raising the possibility that homozygosity of alternate amino acid substitutions residue may be deleterious or even incompatible with life. To explore the functional relevance of the E477A mutation beyond its ability to abolish ORF9b recruitment, we performed mitochondrial phenotyping of *TOMM70^C^*^11^ HeLa FITR cells stably expressing mutant pLV-TOMM70^E477A^ and wild type pLV-TOMM70^WT^. First, we assessed macromolecular complex assembly by Blue Native polyacrylamide gel electrophoresis (BN-PAGE) analyses of isolated mitochondria. As expected, BN-PAGE Immunoblot analyses of mitochondria isolated from TOMM70-deficient HeLa FITR cells revealed an absence of anti-TOMM70 immunoreactivity relative to wild type HeLa FITR cells (Figure 5a). Expression of either pLV-TOMM70^WT^ or pLV-TOMM70^E477A^ in *TOMM70^C^*^11^ HeLa FITR cells elevated the levels of TOMM70 complex to equal levels by BN-PAGE immunoblot (Figure 5a), which is consistent with our observations of steady-state TOMM70 protein levels by SDS-PAGE (Figure S3f) and proteomics (Figure 3i). Notably, the TOMM40 complex did not appear to be disrupted in TOMM70-deficient cells, which is consistent with yeast studies that demonstrate that the assembly and function of TOM70 and TOM40 complexes are distinct and can be genetically uncoupled^30–33^. These findings were corroborated with indirect immunofluorescence studies of TOMM40, which were also not reduced in TOMM70-deficient cells (Figure S4b), indicating that that ablation of TOMM70 in *TOMM70^C^*^11^ HeLa FITR cells does not disrupt TOMM40 nor its macromolecular assembly (Figure 5a). In line, the siRNA-mediated downregulation of *TOMM40* did not phenocopy that of si*TOMM70* in the recruitment of ORF9b^WT^ (Figure S4c). Taken together, these data demonstrate that the *TOMM70^E^*^477^*^A^* mutation that prevents ORF9b binding and recruitment does not interfere with the assembly and/or stability of TOMM70-containing complexes, thereby demonstrating that the role of TOMM70 in maintaining mitochondrial homeostasis can be uncoupled from its function as a receptor for ORF9b.

**Figure 5:**
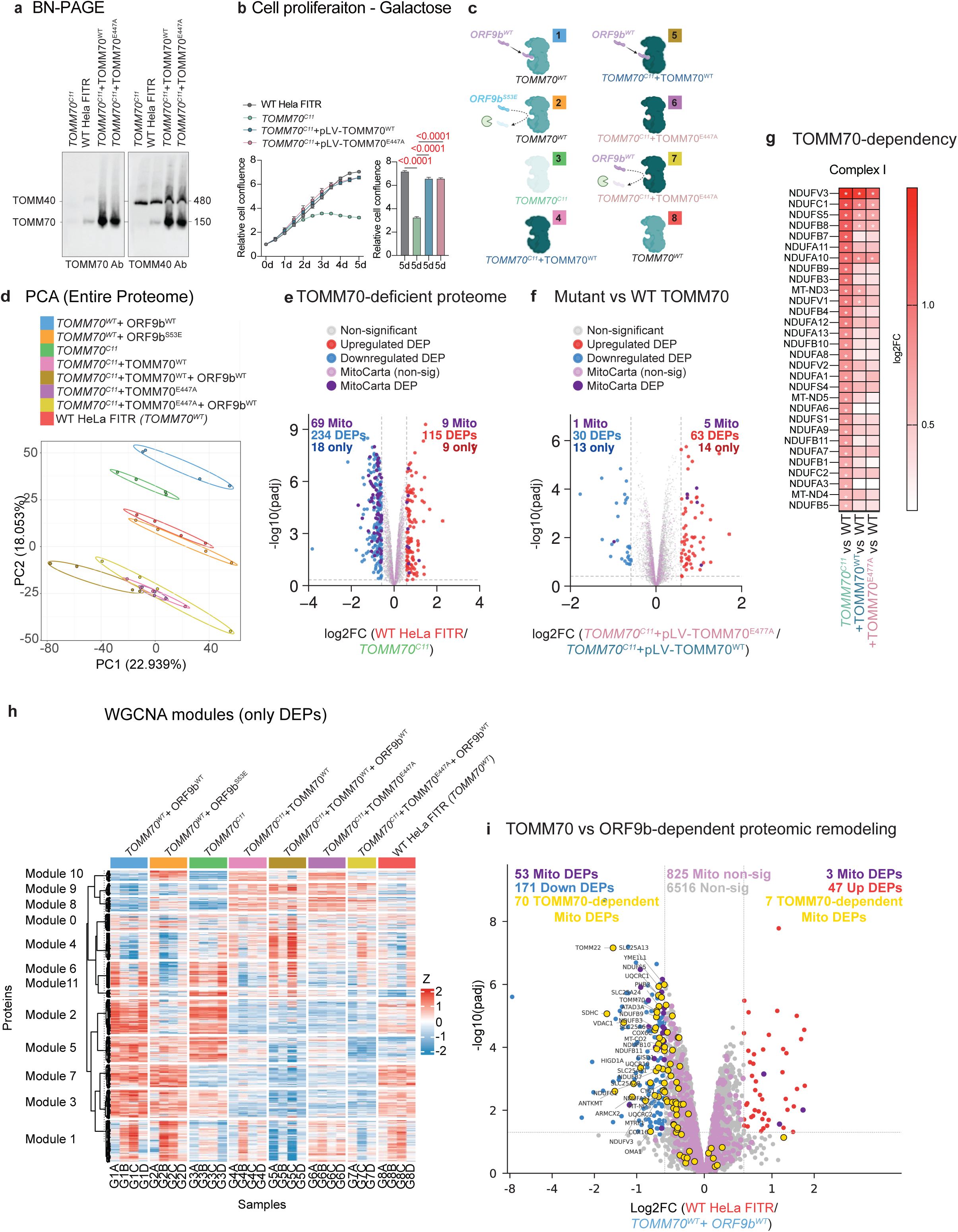
Mutant *TOMM70^E477A^* preserves canonical functions of TOMM70. a) BN-PAGE immunoblots from mitochondria isolated from wild type (WT) HeLa FITR cells, *TOMM70^C^*^11^ HeLa FITR (*TOMM70^C^*^11^) and *TOMM70^C^*^11^ HeLa FITR in which wild type (pLV-TOMM70^WT^) or mutant (pLV-TOMM70^E477A^) were stably expressed. Anti-TOMM70 was probed followed by anti-TOMM40. b) (Left) Cell proliferation in galactose-containing media of wild type (WT) HeLa FITR cells (grey, n=5), *TOMM70^C^*^11^ HeLa FITR cells (*TOMM70^C^*^11^, green, n=5) and *TOMM70^C^*^11^ cells in which wild type (pLV-TOMM70^WT^, blue, n=5) or mutant (pLV-TOMM70^E477A^, pink, n=5) were stably expressed. (Right) Bar graph represents cell confluence after 5 days of proliferation normalized to initial confluence. Data are means ± SEM, one-way ANOVA Tukey’s multiple comparisons test. P-values are indicated and statistically significance is highlighted in red. c) Graphical representation of ORF9b and TOMM70 in 8 groups of HeLa FITR cells explored by whole-cell proteomics. Group 1, wild type (WT) HeLa FITR expressing ORF9b^WT^-FLAG, Group 2, WT HeLa FITR expressing ORF9b^S53E^-FLAG, Group 3, TOMM70-deficient clone 11 (*TOMM70^C^*^11^) HeLa FITR, (grey, n=5), *TOMM70^C^*^11^ HeLa FITR expressing pLV-TOMM70^WT^ (TOMM70^WT^), Group 5, *TOMM70^C^*^11^ HeLa FITR expressing TOMM70^WT^ and ORF9b^WT^-FLAG, Group 6, *TOMM70^C^*^11^ HeLa FITR expressing pLV-TOMM70^E477A^ (TOMM70^E477A^), Group 7, *TOMM70^C^*^11^ HeLa FITR expressing TOMM70^E477A^ and ORF9b^WT^-FLAG, and Group 8 control WT HeLa FITR (*TOMM70^WT^*). See Supplemental Dataset 1. d) Principal component analysis (PCA) of whole-cell proteomes from the 8 groups indicated in Figure 5c. Normalized protein intensities were log2-transformed were used for the PCA. The proportion of variance explained by each principal component is depicted. Sample scores were plotted for the first two principal components (PC1 vs PC2) that explained respectively around 22.94% and 18.05% of variance. See Supplemental Dataset 1. e) Volcano plot of whole-cell proteome of Wild type HeLa FITR (WT, n=4) vs *TOMM70^C^*^11^ HeLa FITR (*TOMM70^C^*^11^, n=4) analyzed by mass spectrometry. (Purple) Mitochondrial proteins (MitoCarta 3.0), (light blue) Differentially expressed proteins (DEPs) more abundant in *TOMM70^C^*^11^ HeLa FITR, (dark blue), proteins exclusively measured in *TOMM70^C^*^11^ HeLa FITR, (light red) DEPs more abundant in WT HeLa FITR, (dark red), proteins exclusively measured in WT HeLa FITR, (grey) non-significant proteins, (light purple) Non-significant MitoCarta 3.0 proteins. Differential analysis using LIMMA t-test is and corresponding p-values are adjusted (padj) using an adaptive Benjamini-Hochberg correction. Differentially expressed proteins (DEPs) are defined by an adjusted p-value is below 0.01 and a log2 fold-change greater than 1, delimited by dotted grey lines. f) Volcano plot of whole-cell proteome of *TOMM70^C^*^11^ HeLa FITR expressing pLV-TOMM70^E477A^ (TOMM70^E477A^, n=4) vs *TOMM70^C^*^11^ HeLa FITR expressing pLV-TOMM70^WT^ (TOMM70^WT^, n=4) analyzed by mass spectrometry. (Purple) Mitochondrial proteins (MitoCarta 3.0), (light blue) Differentially expressed proteins (DEPs) more abundant in *TOMM70^C^*^11^ HeLa FITR expressing TOMM70^WT^, (dark blue), proteins exclusively measured in *TOMM70^C^*^11^ HeLa FITR expressing TOMM70^WT^, (light red) DEPs more abundant in *TOMM70^C^*^11^ HeLa FITR expressing TOMM70^E477A^, (dark red), proteins exclusively measured in *TOMM70^C^*^11^ HeLa FITR expressing TOMM70^E477A^, (grey) non-significant proteins, (light purple) Non-significant MitoCarta 3.0 proteins. Differential analysis using LIMMA t-test is and corresponding p-values are adjusted using an adaptive Benjamini-Hochberg correction. Differentially expressed proteins (DEPs) are defined by an adjusted p-value is below 0.01 and a log2 fold-change greater than 1, delimited by dotted grey lines.. See Supplemental Dataset 1 g) Heatmap of upregulated Complex I proteins in *TOMM70^C^*^11^ HeLa FITR, *TOMM70^C^*^11^ HeLa FITR expressing TOMM70^WT^ (+TOMM70^WT^), and *TOMM70^C^*^11^ HeLa FITR expressing TOMM70^E477A^ (+TOMM70^E477A^) relative to WT HeLa FITR. Differentially expressed proteins (DEPs) are defined by an adjusted p-value is below 0.01 and a log2 fold-change greater than 1, delimited by dotted grey lines..* denotes statistically differences. See Supplemental Dataset 1 h) Proteomics heatmap of module-clustered differentially expressed proteins across eight groups in Figure 5c. Z-score-standardized abundances (centered at zero, diverging color scale) of 2,634 proteins (FDR<0.01, Δ>0.5) from 7,668 imputed/normalized protein groups, filtered for replicate observations and variance. Proteins clustered by WGCNA modules (11 colored modules, n=2,469 proteins; ordered by eigengene similarity; hierarchical clustering within modules using Pearson distance and complete linkage); samples clustered by group. (Complex Heatmap; limma, WGCNA analyses). i) Venn diagram of upregulated mitochondrial DEPs in *TOMM70^C^*^11^ HeLa FITR cells (relative to WT HeLa FITR) and upregulated mitochondrial DEPs in WT HeLa FITR cells expressing ORF9b^WT^-FLAG. j) Volcano plot of whole-cell proteome of HeLa FITR expressing ORF9b^WT^-FLAG (*TOMM70^WT^*+ ORF9b^WT^, n=4) vs WT HeLa FITR cells (WT, n=4) analyzed by mass spectrometry. (Purple) Mitochondrial proteins (MitoCarta 3.0), (light blue) differentially expressed proteins (DEPs) more abundant in *TOMM70^WT^*+ ORF9b^WT^, (light red) DEPs more abundant in WT HeLa FITR, (grey) non-significant proteins, (light purple) Non-significant MitoCarta 3.0 proteins, (yellow) TOMM70-dependent mitochondrial DEPs that are upregulated (70) or downregulated (7) in *TOMM70^C^*^11^ HeLa FITR cells (relative to WT HeLa FITR). Differential analysis using LIMMA t-test is and corresponding p-values are adjusted using an adaptive Benjamini-Hochberg correction. Differentially expressed proteins (DEPs) are defined by an adjusted p-value is below 0.01 and a log2 fold-change greater than 1, delimited by dotted grey lines..

Next, we sought to determine whether TOMM70^E477A^ could functionally complement other mitochondrial defects observed in *TOMM70^C^*^11^ HeLa FITR cells beyond *T. gondii* recruitment to mitochondria. We first measured mitochondrial respiration rates in cells grown in galactose-containing media for at least 3 days in order to exert metabolic demands that have previously been shown to reveal TOMM70-dependent effects in cellular fitness ^16,34^. Surprisingly, *TOMM70^C^*^11^ HeLa FITR cells showed normal basal and maximal oxygen consumption rates measured by Seahorse FluxAnalyzer (Figure S4d). In line, live-cell imaging of mitochondrial membrane potential and content using Tetramethylrhodamine, Ethyl Ester, Perchlorate (TMRE) and MitoTracker DeepRed (MTDR) were both normal (Figure S4e), excluding the use of these mitochondrial readouts to test our hypothesis in this setting. However, *TOMM70^C^*^11^ HeLa FITR cells showed reduced cell proliferation rates in galactose-containing media (Figure 5b) with a limited negative impact on cell (Figure S4f), suggesting that TOMM70 ablation limits cell proliferation under such metabolically-demanding contexts. Using this assay, we compared the proliferation of cells in expressing the mutant version of TOMM70^E477A^ (*TOMM70^C^*^11^ HeLa FITR : pLV-TOMM70^E477A^) to that of cells replete with wild type TOMM70 (both the parental HeLa FITR and *TOMM70^C^*^11^ HeLa FITR: pLV-TOMM70^WT^). Reduced galactose-dependent cell proliferation in *TOMM70^C^*^11^ HeLa FITR cells was rescued by both wild type and E477A mutant TOMM70 expression (Figure 5b), supporting the notion that TOMM70^E477A^ retains mitochondrial function independently of its role in ORF9b recruitment.

We next evaluated the functional impact of the ORF9b-inhibiting E477A mutation in TOMM70 by analyzing the proteomes of wild type and *TOMM70^C^*^11^ HeLa FITR cells, as well as *TOMM70^C^*^11^ HeLa FITR cells expressing wild type (*TOMM70^C^*^11^ HeLa FITR: pLV-TOMM70^WT^) and mutant TOMM70^E477A^ (*TOMM70^C^*^11^ HeLa FITR: pLV-TOMM70^E477A^) (Figure 5c, Supplemental Dataset 1). PCA of whole-cell proteomes revealed clustering of Wild type HeLa FITR cells in a region of the plot that was distinct from that of *TOMM70^C^*^11^ HeLa FITR cells, demonstrating that the composition of these cells was altered (Figure 5d). Comparing the differentially expressed proteins (DEPs) annotated in MitoCarta 3.0 revealed a decrease in several mitochondrial proteins such as CCDC127, MTIF3, and PAICS between WT and *TOMM70^C^*^11^ HeLa FITR cells (Figure 5e). CCDC127 is a poorly-characterized coiled-coil protein implicated in cardiolipin maintenance and mitochondrial ultrastructure^35^ and whose steady-state abundance depend on TOMM70^16^. MTFI3 is a mitochondrial translation initiation factor that is functionally dispensable in HeLa cells^36^ and PAICS is a matrix-localized bifunctional phosphoribosylaminoimidazole carboxylase/succinocarboxamide synthetase enzyme involved in de novo purine biosynthesis. *TOMM70^C^*^11^ HeLa FITR cells displayed an increased abundance of 69 mitochondrial proteins involved in various mitochondrial pathways including OXPHOS assembly, mitochondrial gene expression, protein import and quality control, and nucleotide metabolism (Figure 5e). While the underlying mechanism responsible for this cellular accumulation is unclear, we sought to use them to test the dependency on TOMM70 in *TOMM70^C^*^11^ HeLa FITR cells. 53 out of the 69 mitochondrial DEPs that accumulated in *TOMM70^C^*^11^ HeLa FITR cells were no longer significantly upregulated upon functional complementation with pLV-TOMM70^WT^. Complementation of *TOMM70^C^*^11^ HeLa FITR cells with pLV-TOMM70^WT^ also rescued the depletion of CCDC127, PAICS, and MTIF3 (Figure S4g, Supplemental Dataset 1). These proteins were not differentially expressed between *TOMM70^C^*^11^ HeLa FITR cells expressing pLV-TOMM70^WT^ versus pLV-TOMM70^E477A^ (Figure 5f), further supporting the notion that the E477A mutation in TOMM70 is functionally equivalent to wild type TOMM70 in this regard. In line, we observed that the majority of upregulated DEPs in *TOMM70^C^*^11^ HeLa FITR associated with Complex I and protein homeostasis was resolved by expressing either pLV-TOMM70^WT^ or pLV-TOMM70^E477A^ (Figure 5g). Pairwise comparisons of MitoCarta 3.0-ascribed proteins in pLV-TOMM70^WT^- and pLV-TOMM70^E477A^-expressing *TOMM70^C^*^11^ HeLa FITR cells revealed only ∼0.7% (5 out of 879; MRPL52, DMA1, DNAJC15, DHRS4, and SLC25A21) quantified mitochondrial proteins were differentially expressed beyond 1.5-fold change (Figure 5h, Supplemental Dataset 1) with only MRPL52 differentially expressed beyond a 2-fold change, indicating that these cell lines are virtually identical from a mitochondrial proteome perspective. Beyond the mitochondrial proteome, WGCNA analyses of differentially co-expressed proteins revealed striking similarities between *TOMM70^C^*^11^ HeLa FITR cells expressing pLV-TOMM70^WT^ and pLV-TOMM70^E477A^, which were distinctly separated from *TOMM70^C^*^11^ HeLa FITR cells (Figure 5h, Supplemental Dataset 1). Taken together, these data argue that TOMM70^E477A^ inhibits the recruitment ofORF9b to mitochondria yet retains a similar capacity to that of wild type TOMM70 for non-viral roles.

### TOMM70 ablation and ORF9b recruitment differentially remodel the proteome

ORF9b has been proposed to disrupt mitochondrial protein import via TOMM70 by occupying the hydrophobic pocket of TOMM70 that accepts nuclear-encoded mitochondrial precursor proteins prior to their import via allosteric inhibition^14–16^. ORF9b binds in the c terminal domain (CTD) of TOMM70, clamping it and thereby weaken the engagement of Hsp90, thereby limiting chaperone-mediated delivery and import of TOMM70-dependent clients into mitochondria^16^. ORF9b binding also impairs TOMM70’s N-terminal Hsp90-“EEVD” clamp interaction (and a second Hsp90 contact near helix α7), which is needed to hand off Hsp70/Hsp90-carried preproteins to TOMM70, which would be predicted to reduce the delivery of TOMM70 client proteins to the TOM40 import channel^14,15^. Previous studies in HEK293 and yeast cells have reported an overlap between the proteomic remodeling induced by the depletion of TOMM70 and the expression of ORF9b^16^. We therefore expected to find similar proteomic changes when comparing ORF9b^WT^-FLAG-expressing to *TOMM70^C^*^11^ HeLa FITR cells. In contrast, only half (34/69) of the upregulated MitoCarta 3.0 proteins in *TOMM70^C^*^11^ HeLa FITR cells were also upregulated ORF9b^WT^-FLAG-expressing HeLa FITR cells (Supplemental Dataset 1). Next, we examined our proteomics dataset using Weighted Gene Co-expression Network Analysis (WGCNA), which is a clustering method that groups proteins by how similarly their abundances vary across samples of 8 groups, rather than just asking which individual proteins go up or down^37^. Proteins that are functionally related or co-regulated tend to fluctuate together, and WGCNA captures this by building a correlation network and clustering tightly co-varying proteins into modules (Figure 5h). This revealed distinct module-level expression patterns that separated *TOMM70^C^*^11^ HeLa FITR (Group 3) and ORF9b^WT^-FLAG expressing cells replete with TOMM70 (Group 1) from the parental WT HeLa FITR line (Group 8). Notably, Groups 1 and 3 shared similar profiles for Modules 6 and 2, consistent with a partial proteome-remodeling overlap between TOMM70 ablation and ORF9b expression. However, these groups diverged from Group 2 (ORF9b^S53E^-FLAG-expressing cells) and Group 8 (uninduced WT HeLa FITR), while PCA revealed an overlap between uninduced WT HeLa FITR cells and ORF9b^S53E^-FLAG-expressing cells. This last finding is important because the phosphomimetic S53E mutation prevents mitochondrial recruitment and thus the overlap demonstrates that the proteomic remodeling driven by ORF9b^WT^-FLAG reflects genuine mitochondrial recruitment rather than an off-target transcriptional effect of AHT induction (Figure 5h, Supplemental Dataset 1). These differences were further evident in volcano plots of pairwise comparisons, where mitochondrial DEPs significantly altered in *TOMM70^C^*^11^ HeLa FITR cells were largely absent among DEPs in ORF9b^WT^-FLAG-expressing cells, and vice versa (Figure 5i). Taken together, these data demonstrate that ectopic ORF9b expression remodels the mitochondrial proteome in a manner that is partially distinct from TOMM70 ablation, a divergence that may underlie ORF9b’s circumscribed role in mitochondrial homeostasis.

### Characterization of the effect of ORF9b recruitment on signaling, quality control and bioenergetics

The remodeling of the mitochondrial proteome induced by ORF9b prompted us to explore its impact on mitochondrial functions. Proteomics studies in ORF9b^WT^-expressing HeLa FITR cells uncovered a differential expression of proteins associated with mitochondrial dynamics such as ARMCX2, PHB2, YME1L, and ATAD3A and yeast cells heterologously-expressing ORF9b exhibit altered mitochondrial morphology^16^, so we first examined mitochondrial network morphology by immunofluorescence (Figure 6a). HeLa FITR cells expressing either EGFP- or FLAG-tagged ORF9b^WT^ were stained with MTDR to label the mitochondrial network and TMRE to report on mitochondrial membrane potential. Quantitative analyses did not reveal differences in membrane potential relative to binding-dead ORF9b mutants (Figure 6a). Next, mitochondria were segmented using a custom Fiji macro based on the MTDR channel, followed by binarization to generate total mitochondrial masks. We extracted multiple morphological descriptors, including average mitochondrial area, perimeter, circularity, Feret and MinFeret diameters and solidity, capturing changes in mitochondrial size, length, thickness, and branching complexity. (Figure S5a). Comparing wild type and mutant ORF9b-expressing cells revealed no significant differences in any of these parameters, suggesting that SARS-CoV-2-mediated mitochondrial morphology alterations may be the consequence of viral infection rather than ORF9b expression per se^38^. In line, Seahorse FluxAnalyzer studies that revealed no defects in basal and maximal oxygen consumption rates in HeLa FITR cells induced to express ORF9b^WT^-FLAG or ORF9b^S53E^-FLAG either in glucose-containing media (Figure S5b) nor for in galactose-containing media (Figure 6b). Cell proliferation in galactose-containing media was unaffected in HeLa FITR cells expressing ORF9b^WT^-FLAG or ORF9b^S53E^-FLAG (Figure S5c), further contrasting heterologous ORF9b expression from TOMM70 ablation.

**Figure 6:**
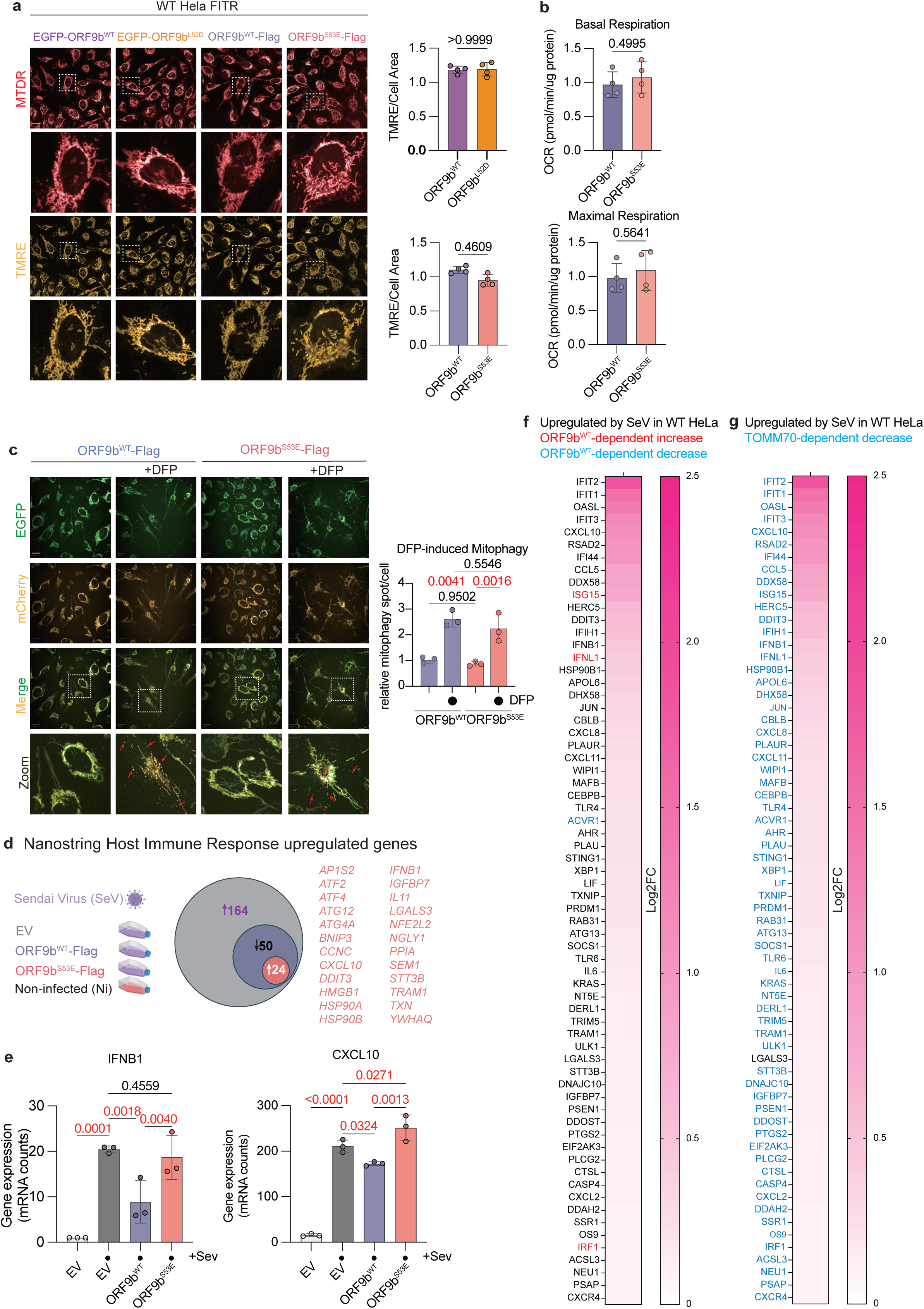
Influence of ORF9b recruitment on signaling, quality control and bioenergetics. a) Representative images of wild type (WT) HeLa FITR induced with AHT to express EGFP-ORFb9WT, EGFP-ORFb9L52D, ORF9bWT-FLAG, or ORF9bS53E-FLAG with anhydrotetracycline (AHT, 10ng/mL) for 15 hours. Mitochondria were labeled with Mitotracker DeepRed (MTDR, red) and Tetramethylrhodamine, Ethyl Ester (TMRE, orange). Scale bar=20 µm. Quantification of single-cell fluorescence signal intensity of mitochondrial membrane potential (TMRE/Cell) of cells in Figure 6a. Data are means ± SEM, 2-tailed unpaired Welch’s t test of 6 replicates with 254-658 cells were imaged for each replicate. Exact P-value indicated. b) Seahorse FluxAnalyzer analysis of mitochondrial respiration in HeLa Flp-In T-REx cells grown in galactose-containing medium for 5 days. Basal and maximal respiration rates were measured in cells treated with anhydrotetracycline (AHT) to induce expression of ORF9bWT (grayish blue) or ORF9bS53E (pale pink) for 5 days. Each condition was measured using multiple technical replicate wells per experiment. Data represent the mean ± SEM of four independent experiments. Statistical significance was assessed using a two-tailed unpaired Student’s t-test. Exact P-value indicated c) Representative images of live imaging of mitophagy in HeLa FITR stably expressing iMLS-EGFP-mCherry mitophagy reporter. Induction of mitophagy with DFP (1mM) and AHT to express ORFb9WT-FLAG or ORFb9S53E-FLAG for 24 hours. Mitochondria engulfed in lysosomes are represented as mCherry+EGFP-mitophagy punctae (red arrows) and quantified using Harmony 5.1. Data are means ± SEM, 2-tailed unpaired Student’s t test. 510-2273 cells were imaged for each replicate. P-values are indicated and statistically significance is highlighted in red. d) Host immune cell response measured by Nanostring mRNA analyses of HeLa FITR cells challenged by Sendai virus (SeV) infection. Wild type (WT) HeLa FITR cells and WT HeLa FITR expressing ORF9bWT-FLAG or ORF9bS53E-FLAG for 24 hours anhydrotetracycline (AHT, 10ng/mL) were subjected to SeV for 2 hours. Fresh media without SeV and with AHT was replenished for an additional 24 hours before Nanostring analysis. Venn Diagram represents differentially express genes (DEGs) induced by SeV in WT cells (164) and in an ORF9bWT-dependent manner (24). e) Expression of IFNB1 and CXCL10 mRNA from Figure 1d. Data are means ± SEM, 2-tailed unpaired Student’s t test. **P<0.01, ***P<0.001. f) Heatmap of SeV-induced upregulated genes in WT HeLa FITR cells that are significantly upregulated (red text) or downregulated (blue text) in SeV-infected ORF9bWT-FLAG HeLa FITR cells. Genes in black text and not significantly different between SeV-infected WT HeLa FITR and SeV-infected ORF9bWT-FLAG HeLa FITR cells. g) Heatmap of SeV-induced upregulated genes in WT HeLa FITR cells that are significantly downregulated (blue text) in SeV-infected TOMM70C11 HeLa FITR cells. Genes in black text and not significantly different between SeV-infected WT HeLa FITR and SeV-infected ORF9bWT-FLAG HeLa FITR cells. Data are generated from Host immune cell response measured by Nanostring mRNA analyses of wild type and TOMM70C11 HeLa FITR cells challenged by Sendai virus (SeV) infection as in Figure 6d.

Viral infection has been associated with an induction of mitophagy, which is predominantly regulated by the molecular remodeling of the outer mitochondrial membrane (OMM)^39^. Since proteins involved in the regulation of cellular and mitochondrial quality control physically interact with ORF9b^9^, we explored whether ORF9b-recruitment to mitochondria could alter mitophagy susceptibility. We stably expressed a fluorescent mitophagy reporter in cells that inducibly express either ORF9b^WT^-FLAG or ORF9b^S53E^-FLAG and performed live imaging under basal and stress conditions. To follow mitophagy in living cells, we used iMLS-QC^40^, which directs a bicistronic EGFP-mCherry protein to the IMM using the first 53 amino acids of the NIPSNAP1 protein, rather than the more commonly used MitoQC^41^ reporter (which is anchored to the OMM by a chimeric FIS1 domain) as we were concerned about possible interference with TOMM70 or ORF9b at the OMM. Under basal conditions, we observed a yellow mitochondrial network that reflected the bicistronic expression of mCherry and EGFP, the latter of which can be quenched by low pH when mitochondria are targeted to the acidic lysosome during mitophagy^40^ (Figure 6c). When we triggered mitophagy with DFP, an iron chelator and most potent inducer of PINK1/Parkin-independent mitophagy^41^, we observed an equivalent increase in mCherry^+^EGFP^−^ punctae in both ORF9b^WT^-FLAG and ORF9b^S53E^-FLAG-expressing cells (Figure 6c), indicating that ORF9b recruitment to mitochondria does not impact mitophagy in HeLa cells.

Next, we explored the impact of ORF9b expression on cell death, reasoning that ORF9b recruitment might perturb the OMM remodeling triggered by established intrinsic triggers that can lead to cell death independently of metabolism. We induced cell death using ABT-737, which induces the oligomerization of BAX and BAK at the OMM, along with Actinomycin D, or in the presence of Staurosporine (STS) and followed cell survival by high-content live imaging of ORF9b^WT^-FLAG and ORF9b^S53E^-FLAG-expressing cells. We observed a time-dependent increase in cell death that was not affected by either ORF9b^WT^-FLAG or ORF9b^S53E^-FLAG (Figure S5d). We obtained similar results with other cell death triggers such as Etoposide or Staurosporine (Figure S5d), indicating that ORF9b recruitment to mitochondria does not alter cell death susceptibility in HeLa cells.

We then explored the impact of ORF9b recruitment on innate immunity. ORF9b recruitment to mitochondria may blunt MAVS-dependent innate immune signaling in response to single-stranded RNA virus sensing^9,10^. How this occurs is unclear, although cellular studies have suggested that phosphorylation of IRF3 at the OMM promotes its translocation to the nucleus^42^. However, live imaging of HeLa FITR cells stably expressing IRF3-EGFP did not reveal a mitochondrial recruitment in response to infection with Sendai virus (SeV), a negative-sense single-stranded RNA paramyxovirus commonly used as a model pathogen to study innate antiviral immunity in mammalian cells (Figure S5e). Nevertheless, we sought to confirm the role of ORF9b in innate immune signaling in HeLa cells and therefore quantified the host immune response by Nanostring 24 hours after SeV infection. Comparing non-infected (Ni) and SeV-infected HeLa cells revealed an upregulation of 168 genes out of the host immune response panel of 773 genes, which include 24 canonical interferon stimulated genes (ISGs) and 106 interferon response–associated genes, as well as factors regulating viral RNA sensing, interferon production and signaling, chemokine responses, apoptosis, and autophagy, consistent with the RIG-I/MDA5-dependent type I interferon (IFN-α/β) response expected upon SeV infection. ORF9b^WT^-FLAG recruitment to mitochondria impacted immune signaling, reducing 50 out of the 164 upregulated genes, which included *CXCL10* and *IFN1b* (Figure 6d). This response was dependent on ORF9b recruitment to mitochondria as ORF9b^S53E^-FLAG expression did not reduce *CXCL10* and *IFN1b* expression (Figure 6f). Only three (ISG15, IFNL1, IRF1) out of the 66 most upregulated host immune response panel genes induced by SeV infection in wild type HeLa FITR cells were further upregulated in HeLa FITR cell expressing ORF9b^WT^-FLAG, circumscribing the limits of ORF9b’s impact on SeV-mediated immune signaling.

We repeated SeV infection experiments in *TOMM70^C^*^11^ HeLa FITR cells, which elicited a more modest host immune response by Nanostring analyses than in WT HeLa FITR cells, indicating that TOMM70-deficient and ORF9b-expressing TOMM70-proficient cells mount different viral immune responses in response to infection (Figure 6f, g). All but one of the 66 most upregulated host immune response panel genes induced by SeV infection in wild type HeLa FITR cells was still significantly induced after SeV infection in *TOMM70^C^*^11^ HeLa FITR cells. We also observed an upregulation of ISGs in *TOMM70^C^*^11^ HeLa FITR cells and wild type HeLa FITR cells silenced for *TOMM70* by siRNA in the absence of infection (Figure S5f, Supplemental Dataset 2), which was not observed for ORF9b-expressing TOMM70-replete cells. These observations indicate that TOMM70 ablation alone can activate innate immune signaling. Altogether, our results indicate that TOMM70 ablation remodels the host immune response differently from the TOMM70-dependent recruitment of ORF9b to mitochondria.

### Characterization ORF9b-deficient SARS-CoV-2 during infection of lung epithelial cells

In light of the limited impact of SARS-CoV-2 ORF9b recruitment on host immune signaling induced by SeV in HeLa cells (Figure 6d-f), we wondered whether intrinsic differences of immune signaling between epithelial cells lines or antiviral host signaling might differ between SeV versus SARS-CoV-2 infection. To resolve these questions, we set out to perform SARS-CoV-2 infection experiments in A549 lung epithelial cells stably expressing human ACE2 and TMPRSS2 (A549^+hACE2+hTMPRSS2^). A549 cells are widely used for the study of SARS-CoV-2 and the ectopic expression of hACE2 and hTMRPSS2 enables maximal viral entry, which we confirmed by immunocytochemistry (Figure S6a). We reverse genetically engineered viral variants lacking ORF9b (SARS-CoV-2^ΔORF9b^) or mutated at L52D (SARS-CoV-2^ORF9bL52D^), which were amplified in Vero-E6 cells and sequence verified by deep sequencing. Briefly, SARS-CoV-2^ΔORF9b^ was generated by introducing T>C transitions at positions 28285 and 28306 to mutate methionine codons M1 and M8, respectively (Figure 7a). The changes do not alter the amino acid sequence of the nucleocapsid (N) protein encoded in an overlapping reading frame. SARS-CoV-2^ORF9bL52D^ was generated using a similar strategy yet led to the obligate introduction of a Leucine to Isoleucine mutation at position 55 in the N protein (A55G) that is conserved across SARS-CoV-2 variants. Western blot analyses confirmed the disruption of ORF9b protein levels in cells infected with SARS-CoV-2^ΔORF9b^ or SARS-CoV-2^ORF9bL52D^, both in A549^+hACE2+hTMPRSS2^ and Vero-E6 cells (Figure 7b, S7b). Immunofluorescence studies using an anti-ORF9b antibody revealed the absence of a signal in SARS-CoV-2^ΔORF9b^ infected cells while the ORF9b signal was diffusely dispersed in SARS-CoV-2^ORF9bL52D^ infected cells (Figure S6c), consistent with the disruption of mitochondrial recruitment of ORF9b we observed using EGFP-ORF9b^L52D^ in HeLa cells (Figure 1a, b)

**Figure 7:**
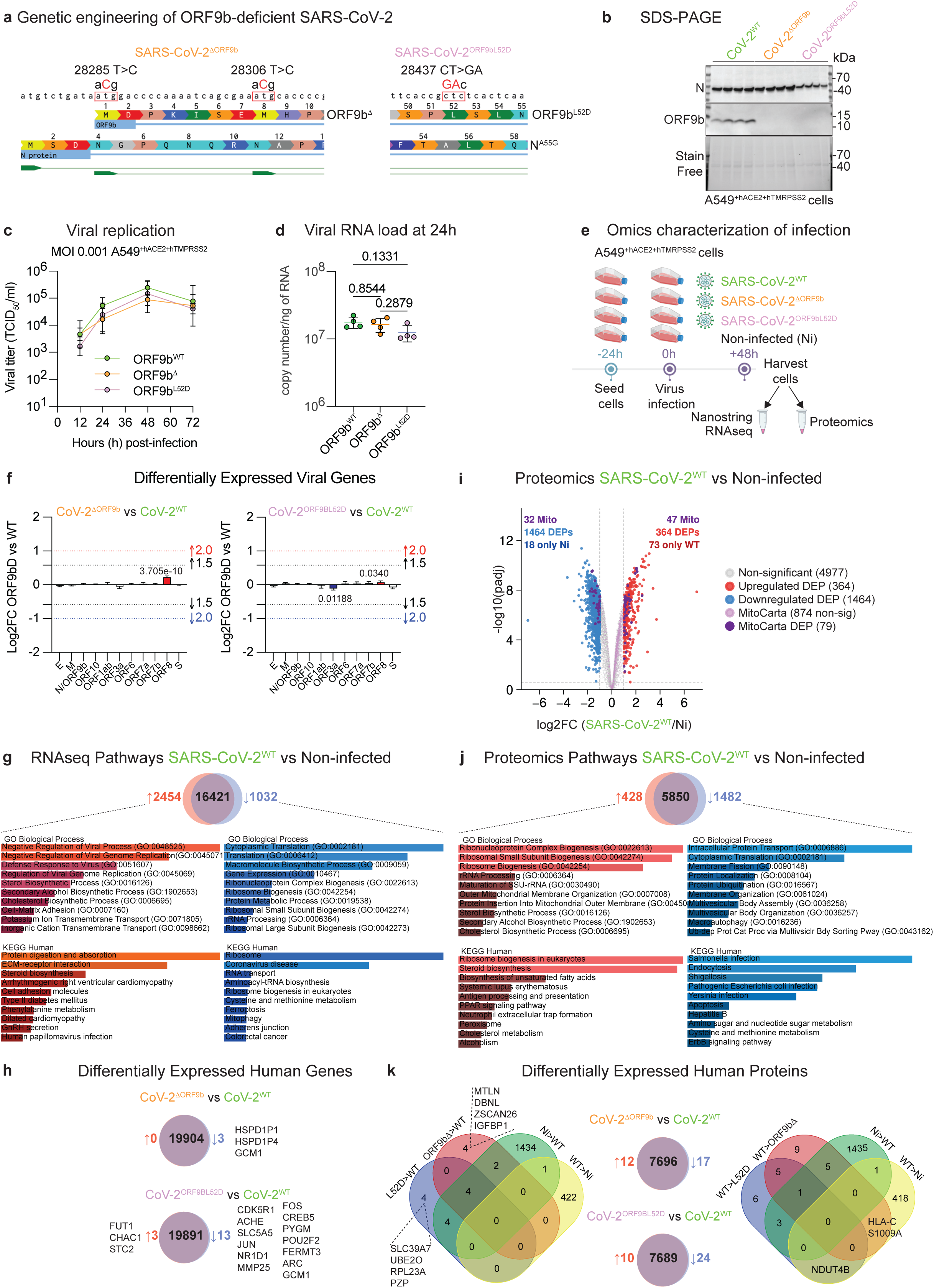
Characterization of ORF9-deficient SARS-CoV-2 virus in A549 epithelial cells. a) ORF9b-deficient SARS-CoV-2 was achieved by reverse genetic engineering. (Left) schematic representation of the two missense mutations in the overlapping reading frames of the Nucleocapsid (N) and ORF9b genes of SARS-CoV-2 (28285C>T and 28306T>C) that were introduced to disrupt the first two Methionine codons in ORF9b to yield SARS-CoV-2ΔORF9b. These mutations did not alter the coding sequence of the N gene (Right) A single missense mutation (dinucleotide substitution 28437CT>GS) was engineered to create the ORF9bL52D variant (SARS-CoV-2Orf9bL52D) and also caused an obligatory A55G missense variant in the N gene (NA55G). b) Immunoblot analysis of whole-cell lysates from A549 lung epithelial cells stably expressing human ACE2 and human TMPRSS2 (A549+hACE2+hTMPRSS2) 24 hours post-infection with wild-type Wuhan SARS-CoV-2 (CoV-2WT), SARS-CoV-2ΔORF9b (CoV-2ΔORF9b), or SARS-CoV-2ORF9bL52D (CoV-2ORF9bL52D) at MOI 0.5 and probed using the indicated antibodies. c) Viral replication of Wuhan SARS-CoV-2 (SARS-CoV-2WT, n=4 green), SARS-CoV-2ΔORF9b (n=4, orange), SARS-CoV-2Orf9bL52D (n=4, pink) viruses at an MOI of 0.001 in A549 cells stably expressing human ACE2 and human TMPRSS2 (A549+hACE2+hTMPRSS2). Viral load was measured by TCID50 assay, Data are means ± SD. d) Viral RNA load measured by qRT-PCR in A549 cells stably expressing human ACE2 and human TMPRSS2 (A549+hACE2+hTMPRSS2) 24 h post-infection with Wuhan SARS-CoV-2 (SARS-CoV-2WT, n=4 green), SARS-CoV-2ΔORF9b (n=4, orange), SARS-CoV-2Orf9bL52D (n=4, pink), viruses. Data are means ± SD, one-way ANOVA Tukey’s multiple comparisons test. P values are indicated and statistically significance is highlighted in red. e) Workflow for the characterization of A549+hACE2+hTMPRSS2 infected with Wuhan SARS-CoV-2 (SARS-CoV-2WT), SARS-CoV-2ΔORF9b, and SARS-CoV-2Orf9bL52D viruses (MOI 0.05) and Non-infected (Ni) cells using Nanostring (Supplemental Dataset 3), bulk RNAseq (Supplemental Dataset 4) and whole-cell proteomics (Supplemental Dataset 6). f) Differential expression of SARS-CoV-2 viral genes determined by bulk RNAseq. Pairwise comparisons between (left) Wuhan SARS-CoV-2 (CoV-2WT) versus SARS-CoV-2ΔORF9b (CoV-2ΔORF9b) and (right) CoV-2WT versus SARS-CoV-2Orf9bL52D (CoV-2Orf9bL52D)-infected A549+hACE2+hTMPRSS2 cells. Pvalues are indicated where significantly upregulated (red) and downregulated (blue) genes were identified. Fold-change thresholds of 1.5 (black) and 2.0 (red or blue) are indicated by dotted lines. Data are available in Supplemental Dataset 4. g) RNAseq identification of differentially expressed genes (DEGs) in A549+hACE2+hTMPRSS2 cells infected with Wuhan SARS-CoV-2 (SARS-CoV-2WT, n=4) versus non-infected A549+hACE2+hTMPRSS2 cells (n=4) 48 post-infection. Venn diagram shows 16421 overlapping genes (purple), 2454 upregulated DEGs (red) and 1032 downregulated DEGs (blue) caused by infection with SARS-CoV-2WT. Pathway analysis using Gene Ontology (GO) and KEGG databases revealed enrichment of pathways related to viral infection and replication, ECM remodeling, cholesterol metabolism, and cytosolic protein translation (see Supplemental Dataset 5). h) Venn diagram of DEGs in (top) SARS-CoV-2ΔORF9b (CoV-2ΔORF9b) versus Wuhan SARS-CoV-2 (CoV-2WT) and (bottom) SARS-CoV-2Orf9bL52D (CoV-2Orf9bL52D) versus CoV-2WT versus-infected A549+hACE2+hTMPRSS2 cells. Zero upregulated (red) and 3 downregulated (blue) DEGs were observed between CoV-2ΔORF9b and CoV-2WT. 3 upregulated (red) and 13 downregulated (blue) DEGs were observed between CoV-2Orf9bL52D and CoV-2WT. i) Volcano plot of whole-cell proteome of A549+hACE2+hTMPRSS2 cells infected with Wuhan SARS-CoV-2 (SARS-CoV-2WT, n=4) versus non-infected A549+hACE2+hTMPRSS2 cells (n=4) 48 post-infection analyzed by mass spectrometry. (Dark Purple) differentially expressed mitochondrial proteins (MitoCarta 3.0), (light purple) non-significant mitochondrial proteins, (blue) downregulated differentially expressed proteins (DEPs), (red) upregulated DEPs. Differential analysis using LIMMA t-test is and corresponding p-values are adjusted using an adaptive Benjamini-Hochberg correction. DEPs are defined by an adjusted p-value is below 0.01 and a log2 fold-change greater than 1, delimited by dotted grey lines., (see Supplemental Dataset 6). j) Identification of differentially expressed proteins (DEPs) in A549+hACE2+hTMPRSS2 cells infected with Wuhan SARS-CoV-2 (SARS-CoV-2WT, n=4) versus non-infected A549+hACE2+hTMPRSS2 cells (n=4) 48 post-infection. Venn diagram shows 5850 overlapping proteins (purple), 428 upregulated DEPs (red) and 1482 downregulated DEPs (blue) caused by infection with SARS-CoV-2WT. Pathway analysis using Gene Ontology (GO) and KEGG databases revealed enrichment of pathways related to ribosomal biogenesis, rRNA processing, cytosolic protein translation, and pathogen infection (see Supplemental Dataset 7). Source data are available for this figure (Supplemental Datasets 4, 5). k) Venn diagram of DEPs in (center, top) SARS-CoV-2ΔORF9b (CoV-2ΔORF9b) versus Wuhan SARS-CoV-2 (CoV-2WT) and (center, bottom) SARS-CoV-2Orf9bL52D (CoV-2Orf9bL52D) versus CoV-2WT versus-infected A549+hACE2+hTMPRSS2 cells. 12 upregulated (red) and 17 downregulated (blue) DEGs were observed between CoV-2ΔORF9b and CoV-2WT. 10 upregulated (red) and 24 downregulated (blue) DEPs were observed between CoV-2Orf9bL52D and CoV-2WT. Commonly upregulated DEPs (left) or downregulated DEPs (right) between SARS-CoV-2ΔORF9b (ΔΔOrf9b) and SARS-CoV-2Orf9bL52D (L52D) relative to Wuhan SARS-CoV-2 (WT) or non-infected (Ni) cells are indicated.

To assess the impact of ORF9b on viral fitness, we next compared replication kinetics of SARS-CoV-2^WT^, SARS-CoV-2^ΔORF9b^ and SARS-CoV-2^ORF9b^^L52D^ by TCID_50_ titration in A549^+hACE2+hTMPRSS2^ cells and found no significant differences in infectious virus production among the three variants (Figure 7c), which we corroborated in Vero-E6 cells (Figure S6d). Viral genome quantification by qPCR revealed indistinguishable levels of SARS-CoV-2 24 hours post infection in A549^+hACE2+hTMPRSS2^ cells (Figure 7d), indicating that ORF9b is dispensable for viral entry and replication. NGS of viral genomes did not reveal mutational hotspots or sequence variants that differed significantly between SARS-CoV-2^WT^, SARS-CoV-2^ΔORF9b^ and SARS-CoV-2^ORF9bL52D^ (Supplemental Table 4), indicating that ORF9b does not regulate viral genome integrity. Next, we measured the host immune response of A549^+hACE2+hTMPRSS2^ cells after 48h of infection using the Nanostring nCounter Host Response Panel, which revealed a significant upregulation of 152 of the 791 genes compared to non-infected (Ni) control cells, including ISGs such as *IFIT1*, *IFITM1*, *IFI6*, *MX1*, *IFI44*, *OAS1*, *OAS2*, *OAS3*, and *OASL* (Figure S6e, Supplemental Dataset 3). Of the 152 genes that were significantly upregulated by SARS-CoV-2^WT^, only one gene, IL31, was differentially expressed in both SARS-CoV-2^ΔORF9b^ and SARS-CoV-2^ORF9bL52D^ infected cells. IL31 is a cytokine typically associated with type 2 immune responses and predominantly produced by activated T helper cells. In contrast, a limited number of genes, including PRKCSH, LAMP2, and ITGAX, were modestly upregulated under the same conditions. However, these changes did not converge on a coherent functional pathway and were not associated with alterations in type I interferon signaling or canonical antiviral gene programs. (Supplemental Dataset 3). Altogether, these data strongly argue that ORF9b has a limited effect on SARS-CoV-2-dependent innate immune signaling in lung epithelial cell lines.

To gain a more comprehensive view of ORF9b-dependent molecular remodeling induced by SARS-CoV-2, we performed comparative bulk RNAseq and proteomic studies on Ni and SARS-CoV-2^WT^, SARS-CoV-2^ΔORF9b^ and SARS-CoV-2^ORF9bL52D^ infected A549 cells at 48h (Figure 7e). We were able to detect the expression of SARS-CoV-2 genes, including S, N/ORF9b, M, E, ORF1 (a and b), ORF3, ORF6, ORF7a, ORF7b, and ORF10, none of which were dysregulated beyond 1.5 fold in SARS-CoV-2^ΔORF9b^ and SARS-CoV-2^ORF9bL52D^ infected cells (Figure S6f), consistent with our NGS viral detection studies demonstrating no significant differences in viral RNA levels between WT and ORF9b-deficient viruses (Figure 7c, d). Beyond ORF9b, none of the other viral proteins we measured by proteomics were dysregulated in SARS-CoV-2^ΔORF9b–^infected cells, including the N protein (Figure S6f), consistent with western blot in A549^+hACE2+hTMPRSS2^ cells (Figure 7b). While N protein levels were also unchanged in SARS-CoV-2^ORF9bL52D^ infected cells, we observed a ∼2-fold depletion of ORF6 and ORF8 by proteomics (Figure S6f), which was not associated with altered viral replication (Figure 7c, d).

Comparing Ni and SARS-CoV-2^WT^ transcriptomes revealed an upregulation of 2454 genes (12.3%) and a downregulation of 1032 genes (5.2%) beyond a threshold of >2-fold change (Log2FC>1). Pathway enrichment analyses of the upregulated transcriptome reflected potent antiviral immunity with negative regulation of viral process, defense response to virus, and response to interferon-beta as top pathways (Figure 7f), as previously reported^10^. Simultaneously, sterol and cholesterol biosynthetic processes were elevated, consistent with the notion that host cell biosynthesis is hijacked to support viral membrane biogenesis. Dysregulation of ion transmembrane transport (potassium, sodium, chloride) and extracellular matrix organization were also consistent with cellular structural and metabolic remodeling during infection. On the other hand, the downregulated transcriptome showed coordinated suppression of protein synthesis machinery, including cytoplasmic translation, ribosome biogenesis, and rRNA processing—a canonical viral strategy to redirect ribosomes toward viral protein synthesis^43^. Most critically, mitochondrial translation, mitochondrial gene expression, and protein targeting to mitochondrion were markedly suppressed, consistent with the notion that ORF9b’s disruption of mitochondrial biogenesis.

Next, we used Gene Set Enrichment Analyses (GSEA) to further probe transcriptional remodeling. GSEA analyses revealed a suppression of ribosome pathways that include mitochondrial ribosomes (hsa03010, hsa03008), Oxidative Phosphorylation (hsa00190), and Coronavirus disease - COVID-19 (hsa05171) (Figure S6h, Supplemental Dataset 7) and an upregulation of pathways associated with cytokine signaling, pathogen infection, and ECM remodeling, which is consistent with previous findings from biopsies of SARS-CoV-2 infected humans^7^. However, pairwise comparisons of SARS-CoV-2^WT^ and SARS-CoV-2^ΔORF9b^ using stringent cutoffs revealed remarkably similar transcriptomes: no genes were upregulated (>Log2FC, padj<0.05) and only 3 genes (*GCM1*, *HSPD1P1*, *HSPD1P4*) were downregulated leaving 19904 genes not differentially expressed including all known factors involved in innate immunity, inflammation, and MitoCarta 3.0. From a transcriptional perspective, these data suggest that ORF9b is dispensable for remodeling of mitochondria and innate immune signaling. Of the 3 ORF9b-dependent DEGs, *HSPD1PD1* and *HSPD1PD4* are pseudogenes, and *GCM1*, which was also downregulated in SARS-CoV-2^ORF9bL52D^ infected cells (Figure 7g), encodes a transcription factor known to be upregulated in Flu-infected A549 cells. In contrast to SARS-CoV-2^ΔORF9b,^ SARS-CoV-2^ORF9b^^L52D^ infection yielded 3 upregulated genes (FUT1, CHAC1, STC2) and 13 downregulated genes (*CDK5R1, ACHE, SLC5A5, JUN, NR1D1, MMP25, FOS, CREB5, PYGM, POU2F2, FERMT3, ARC, GCM1*). CHAC1, which encodes a γ-glutamyl cyclotransferase that specifically degrades glutathione (GSH) to regulate redox balance, was also upregulated in SARS-CoV-2^ΔORF9b^ by 1.92-fold (Log2FC=0.941), suggesting that ORF9b might be involved in modulating redox-sensitive processes such as apoptosis and ferroptosis. CHAC1 expression is induced by transcription factors such as ATF4 as part of the integrated stress response (ISR) that can be mediated by various inputs including mitochondrial dysfunction (OMA1-DELE1-HRI), ER stress (PERK), viral infection (PKR), and amino acid starvation (GCN2) (Figure S6g). However, we observed little overlap between the list of the 94 ATF4-dependent ISR human genes^44^ and the DEGs identified in Ni and SARS-CoV-2^WT^: of the 2545 human genes upregulated by SARS-CoV-2^WT^, only 0.2% were shared (LAMP3, MXD1, APOL6, and NES), none of which were differentially altered in SARS-CoV-2^ΔORF9b^ or SARS-CoV-2^ORF9bL52D^ infected cells (Figure 7g), indicating that SARS-CoV-2, in contrast to other mitochondrial-targeting pathogens such as *T. gondii*^45^, does not trigger ISR activation in A549 cells. Despite a limited number of DEGs, GSEA analyses comparing SARS-CoV-2^WT^ and SARS-CoV-2^ΔORF9b^ revealed differences in pathways associated with DNA replication and repair as well as the metabolism of fatty acids and hormones, which has previously been implicated in COVID-19^46^. SARS-CoV-2^ΔORF9b^ infection was associated with an activation of ribosome (and mitochondrial ribosome) genes relative to SARS-CoV-2^WT^ infection, but without a measurable enrichment of Oxidative Phosphorylation or innate immune pathways (Figure S6i).

Consistent with the trends observed by bulk RNAseq, the substantial proteomic remodeling between Ni and SARS-CoV-2^WT^ infected cells was largely preserved in ORF9b-deficient SARS-CoV-2 variants (Figure 7k). Yet in contrast to the transcriptomic remodeling, downregulated differentially expressed proteins (DEPs) outnumbered upregulated DEPs. Comparisons of Ni and SARS-CoV-2^WT^ revealed 428 proteins were upregulated (5.6%), 1428 proteins were downregulated (18.5%), and 5850 were not differentially expressed beyond 2-fold (Figure 7h, i). Among the DEPs ascribed to MitoCarta 3.0, 50 were increased and 32 were decreased. We noticed an upregulation of the mitochondrial fission regulator DNM1L and a downregulation of the mitochondrial fusion protein OPA1 (Figure 7h) prompting us to assess mitochondrial morphology. Quantitative analyses of mitochondrial morphology in fixed, SARS-CoV-2^WT^ infected, N protein-positive cells revealed a modest yet significant reduction in mitochondrial fragmentation (Figure S6j), consistent with previous label-free imaging studies of SARS-CoV-2^WT^ infected U2OS cells^38^. However, the degree of mitochondrial fragmentation was equivalent in SARS-CoV-2^ΔORF9b^ or SARS-CoV-2^ORF9bL52D^ infected cells (Figure S6j), indicating that these alterations are independent of ORF9b. In line, none of the MitoCarta 3.0 DEPs were altered between wild type and mutant SARS-CoV-2. Of the 10 and 12 proteins that were upregulated in SARS-CoV-2^ΔORF9b^ and SARS-CoV-2^ORF9bL52D^ infected cells, respectively, compared to SARS-CoV-2^WT^ infected cells, only 4 were common to both mutant viruses: SLC39A7, UBE2O, RPL23A, PZP. SLC39A7 is a Golgi/ER zinc transporter previously identified as a specific interactor of SARS-CoV-2 nsp4, implicating it in virus–host interface remodeling and TNF receptor trafficking during infection. UBE2O is a multifunctional E2/E3 ubiquitin-conjugating enzyme that can reshape the ubiquitin–proteasome landscape, and RPL23A encodes a 60S ribosomal protein linked to the monoubiquitin–ribosomal fusion protein RPS27A, which directly interacts with SARS-CoV-2 and is implicated in viral translation control. PZP is a broad-spectrum protease inhibitor with immunomodulatory activity. On the other hand, of the 17 and 24 proteins that were downregulated in SARS-CoV-2^ΔORF9b^ and SARS-CoV-2^ORF9bL52D^ infected cells, respectively, only 2 were common to both mutant viruses: NDUT4B and S110A8. NDUT4B, which is a cytosolic enzyme that cleaves the beta-phosphate from diphosphate groups in diphosphoinositol polyphosphates to regulate their turnover, was downregulated by wild-type SARS-CoV-2 and further downregulated by ORF9b-deficient viruses. While Nudix hydrolases such as NUDT2 have been implicated in the degradation of 5′-triphosphorylated viral RNA^47^, no involvement of NUDT4B (or its paralogue NUDT4) in viral genome regulation has been reported. S100A8 is a calcium- and zinc-binding protein from the S100 family that heterodimerizes with S100A9 to regulate inflammatory processes and immune responses. We observed elevated levels of S100A8 upon infection with SARS-CoV-2^WT^, which was significantly blunted in SARS-CoV-2^ΔORF9b^ and SARS-CoV-2^ORF9bL52D^ infected cells (Figure 7i, j), however S100A9 was not among the DEPs in our study calling into question the functional relevance of S100A8 dysregulation. Taken together, these data argue for a limited impact of ORF9b on innate immunity and mitochondrial proteome remodeling and content upon acute infection of A549 cells.

### Defining the physiological relevance of ORF9b-deficient SARS-CoV-2 viruses

The pathophysiological effects of SARS-CoV-2 can be investigated *in vivo* using the golden Syrian hamster (*Mesocricetus auratus*), which is naturally permissive to infection and faithfully recapitulates many COVID-19–associated phenotypes in humans, including the characteristic lung pathology that underlies severe disease^7,48,49^. To study the *in vivo* relevance of ORF9b, male adult golden Syrian hamsters were intranasally inoculated with 6 x 10^4^ PFU of either SARS-CoV-2 of SARS-CoV-2^WT^, SARS-CoV-2^ΔORF9b^ and SARS-CoV-2^ORF9bL52D^ (Figure 8a). This inoculum size was selected based on our previous studies^49^ and 4 day endpoint was chosen because SARS-CoV-2^WT^ invariably causes symptomatic infection in golden Syrian hamster, with a high incidence of clinical severity and high viral loads in the upper and lower respiratory tracts within 4 days post-inoculation (dpi). *In vivo* experiments were performed in males, as clinical signs are more evident than in females following intranasal SARS-CoV-2 infection^49,50^. As expected, we observed a progressive decrease in body weight and an increase in clinical score in hamsters infected with SARS-CoV-2^WT^ relative to mock-infected control hamsters (Figure 8b, c, S8a). These responses were unchanged in hamsters infected with either SARS-CoV-2^ΔORF9b^ and SARS-CoV-2^ORF9bL52D^. At 4 dpi, lung weight/body weight ratios were equally elevated in SARS-CoV-2^WT^ and SARS-CoV-2^ΔORF9b^ infected animals indicating that the lung pathology (edema, congestion, inflammation) is not dependent upon ORF9b (Figure 8d). In line, lung viral titers and viral RNA loads were indistinguishable in hamsters infected with wild type or mutant viruses (Figure 8e, f), and the accumulation of viral genome mutations was similarly comparable between variants (Supplemental Table 5), indicating that ORF9b is dispensable for viral replication and SARS-CoV-2 virulence in the lungs. Lung histopathological analyses revealed congestion, oedema, desquamation of the bronchiolar epithelium and massive, predominantly mononuclear, inflammatory infiltration in the alveolar and peribronchiolar areas (Figure 8h). SARS-CoV-2 N protein staining revealed wide-spread dissemination of SARS-CoV-2 (Figure 8g) in the bronchiolar epithelium, and in the inflamed alveolar and peribronchiolar areas. These changes were accompanied by TUNEL staining indicating widespread cell death in regions of the lung that were subject to inflammatory remodeling and infection (Figure 8g). Quantitative analyses of TUNEL and H&E staining revealed no significant differences in cell death (Figure 8g) nor lung damage (Figure 8h) in hamsters infected with either SARS-CoV-2^ΔORF9b^ versus SARS-CoV-2^WT^, arguing that ORF9b does not significantly contribute to SARS-CoV-2 lung pathology *in vivo*.

**Figure 8:**
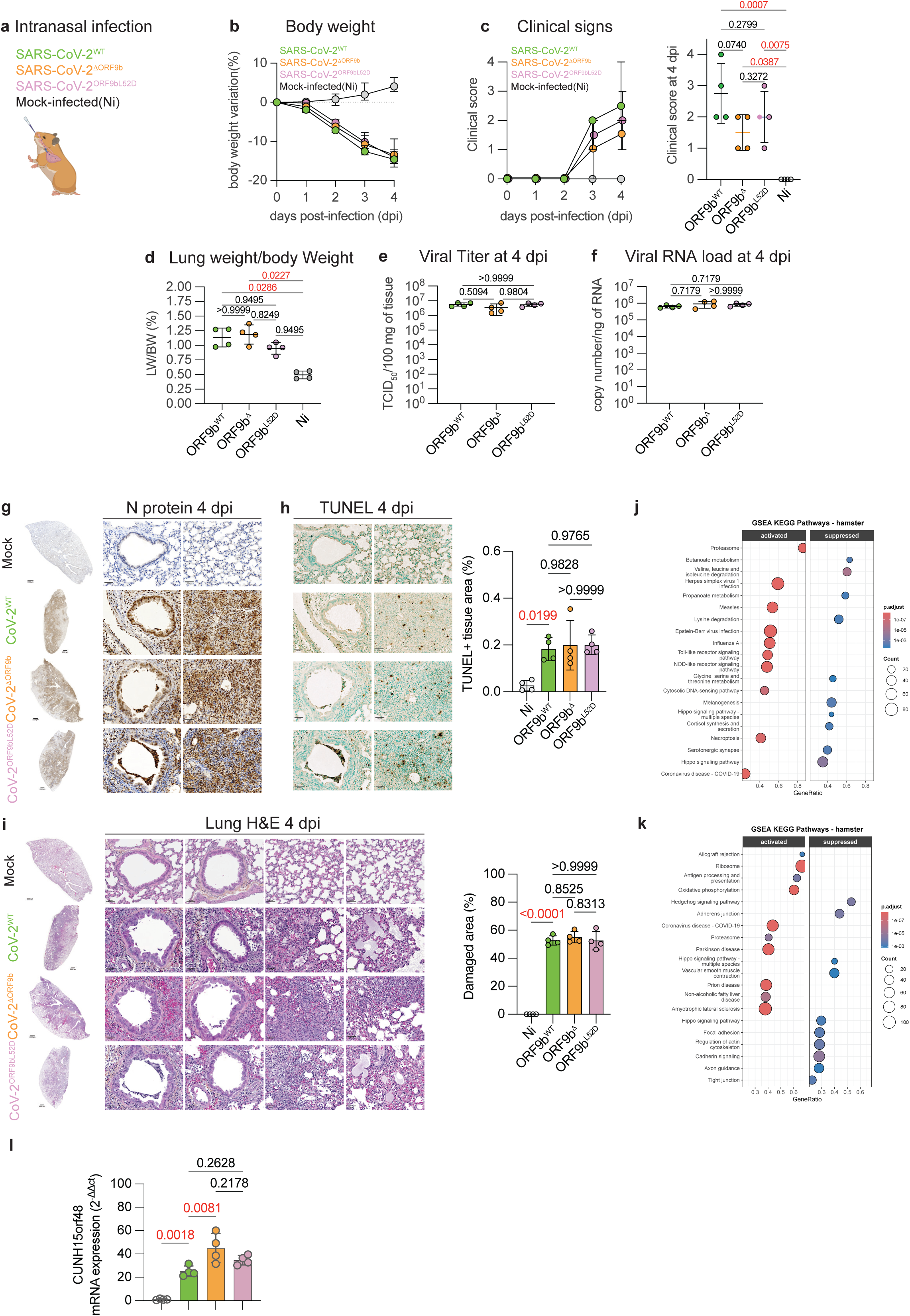
Characterization of ORF9-deficient SARS-CoV-2 infection in hamsters. a) Intranasal infection of Syrian golden hamsters with 6 x 104 PFU of either Wuhan SARS-CoV-2 (SARS-CoV-2WT), SARS-CoV-2ΔORF9b, and SARS-CoV-2Orf9bL52D viruses. b) Body weight variation over four days post-infection of hamsters in Figure 8a. c) (Left) clinical score over four days post-infection of hamsters in Figure 8a. The clinical score is based on a cumulative 0–4 scale: ruffled fur; slow movements; apathy; and absence of exploration activity. (Right) clinical score at 4 days post-inoculation (dpi). Data are means ± SD, one-way ANOVA Tukey’s multiple comparisons test. P values are indicated and statistically significance is highlighted in red. d) Lung weight to body weight (LW/BW) ratio at 4 dpi of hamsters in Figure 8a. Data are means ± SD, one-way ANOVA Tukey’s multiple comparisons test. P values are indicated and statistically significance is highlighted in red. e) Infectious viral titer measured by TCID50 assay per 100mg of lung tissue at 4 dpi. Data are means ± SD, one-way ANOVA Tukey’s multiple comparisons test. Exact p-values are indicated. f) Viral RNA load measured by RT-qPCR. Data are means ± SD, one-way ANOVA Tukey’s multiple comparisons test. Exact p-values are indicated and statistically significance is highlighted in red. g) Representative immunohistochemistry for SARS-CoV-2 Nucleocapsid (N) protein staining of 6-week old male Syrian hamsters at 4 days post-infection (dpi) with Wuhan SARS-CoV-2 (CoV-2WT, n=4), SARS-CoV-2ΔORF9b (CoV-2ΔORF9b, n=4) SARS-CoV-2Orf9bL52D (CoV-2Orf9bL52D, n=4) viruses or non-infected Mock controls (n=4). h) Representative immunohistochemistry of TUNEL staining of cell death (right) of 6-week old male Syrian hamsters at 4 days post-infection (dpi) with Wuhan SARS-CoV-2 (CoV-2WT, n=4), SARS-CoV-2ΔORF9b (CoV-2ΔORF9b, n=4) SARS-CoV-2Orf9bL52D (CoV-2Orf9bL52D, n=4) viruses or non-infected Mock controls (n=4). Quantification of % TUNEL-positive tissue area by QuPath-based image analysis of entire sections. Data are means ± SD, one-way ANOVA Tukey’s multiple comparisons test. Exact p-values are indicated and statistically significance is highlighted in red. i) Representative hematoxylin and eosin (H&E) staining of 6-week old male Syrian hamsters at 4 days post-infection (dpi) with Wuhan SARS-CoV-2 (CoV-2WT, n=4), SARS-CoV-2ΔORF9b (CoV-2ΔORF9b, n=4) SARS-CoV-2Orf9bL52D (CoV-2Orf9bL52D, n=4) viruses or non-infected Mock controls (n=4). Quantification of % damaged tissue area (dark purple) by QuPath-based image analysis of entire sections. Data are means ± SD, one-way ANOVA Tukey’s multiple comparisons test. Exact p-values are indicated and statistically significance is highlighted in red. j) Gene Set Enrichment Analyses (GSEA) representation of activated and suppressed KEGG pathways comparing bulk RNAseq transcriptomes of Wuhan SARS-CoV-2 (SARS-CoV-2WT) versus non-infected hamsters in Figure 8a. Data are available in Supplemental Dataset 7. k) Gene Set Enrichment Analyses (GSEA) representation of activated and suppressed KEGG pathways comparing bulk RNAseq transcriptomes of SARS-CoV-2ΔORF9b (CoV-2ΔORF9b) versus Wuhan SARS-CoV-2 (CoV-2WT) infected hamsters in Figure 8a. Data are available in Supplemental Dataset 7. l) Quantification of C15ORF48 mRNA levels by RT-qPCR performed on lung biopsies from animals in Figure 8g: 6-week old male Syrian hamsters at 4 days post-infection (dpi) with Wuhan SARS-CoV-2 (CoV-2WT, n=4), SARS-CoV-2ΔORF9b (CoV-2ΔORF9b, n=4) SARS-CoV-2Orf9bL52D (CoV-2Orf9bL52D, n=4) viruses or non-infected Mock controls (n=4). Data are means ± SEM, one-way ANOVA Tukey’s multiple comparisons test. P-values are indicated and statistically significance is highlighted in red.

To gain insights into molecular remodeling driven by ORF9b-deficient SARS-CoV-2, we performed bulk RNAseq on lungs of hamsters at 4 dpi. As expected, SARS-CoV-2^WT^ infection triggered the upregulation of 1668 genes enriched in immune system, cytokine signaling, interferon and interleukin-10 genes (Figure S7b) when compared to mock-infected lungs. This is consistent with the diffuse alveolar damage, inflammatory infiltration, and endothelial injury seen at peak acute disease. 1331 downregulated genes were associated with collagen biosynthesis and degradation, which is consistent with ECM remodeling and early acute injury features of basement-membrane disruption and stalled repair. We also observed a suppression of pathways related to angiogenesis and vasculogenesis, which could reflect the endothelial injury and microvascular dysfunction that accompanies transient suppression of angiogenic signaling^51,52^. Downregulated genes were also enriched in pathways associated with abnormal trachea and pulmonary morphology, which is consistent with the histopathological lesions characterized by bronchial epithelial loss, ciliary dysfunction, and interstitial and intra-alveolar edema and immune cell infiltrates. GSEA analyses revealed an enrichment of pro-inflammatory, viral infection, and immune pathways that were activated by SARS-Cov-2 infection and a suppression of branched chain amino acid metabolism and Hippo signaling pathways (Figure 8i), evocative of previously reported metabolic and mechanosensing alterations in humans^53^. Transcriptomic analyses with GSEA comparing SARS-CoV-2^WT^ versus SARS-CoV-2^ΔORF9b^ infected lungs revealed the top three pathways transcriptionally activated by ORF9b-deficient virus to be Ribosomes (cge03010), Oxidative phosphorylation (cge00190), and Coronavirus disease - COVID-19 (cge05171). Additionally, we observed the activation of pathways including Antigen presentation, Allograft rejection, Autoimmune thyroid disease, Rheumatoid arthritis and the suppression of Hippo signaling in SARS-CoV-2^ΔORF9b^ infected lungs, which is consistent with the notion that ORF9b ablation further activates inflammatory signaling and metabolic remodeling *in vivo* (Figure 8j). Using stringent, classical cutoffs (−1>LOG2FC>1) comparing DEGs between SARS-CoV-2^WT^-infected lungs and SARS-CoV-2^ORF9bL52D^-infected lungs, we detected only two DEGs (out of 16906 genes): CUNH1orf115, which was downregulated roughly two-fold (Log2FC= −1.03) and CUNH15orf48, which was up-regulated two-fold (Log2FC=1.01) (Figure S7c). Neither CUNH1orf115 (141 amino acids) nor CUNH15orf48 (83 amino acids) have been functionally characterized in Syrian hamsters, yet the orthologous proteins in humans identified by BLAST are RDD1 (require for drug-induced death 1)^54^ and COXFA4l3 (Cytochrome c oxidase associated subunit FA4L3), respectively. In humans, RDD1 loss-of-function has been associated with resistance to cell death from an array of chemotherapeutic agents^55^ by modulating ABDB1/MDR1-dependent drug efflux. C15ORF48/COXFA4L3, also known as MOCCI (for Modulator of cytochrome C oxidase during Inflammation) is a nuclear-encoded mitochondrial protein that is upregulated during viral infection and inflammation, substitutes for a NDUFA4 (COXFA4), a canonical structural subunit of Complex IV, leading to reduced enzymatic activity, mitochondrial membrane potential and ROS production^56,57^. In COVID-19 patients, the upregulation of C15ORF48 is well-documented and has been proposed to be both induced by and responsible for inflammatory remodeling in response to cytokine and viral triggers *in vitro*^56–58^. In murine macrophages, the genetic deletion or degradation of NDUFA4 is sufficient to boost innate immune signaling and confer anti-tumor immunity in vivo^59^. By RT-qPCR, we validated that CUNH15ORF48 mRNA (*C15ORF48* in humans) was upregulated by SARS-CoV-2^WT^ infection and further induced SARS-CoV-2^ΔORF9b^ infection in the lungs of infected hamsters (Figure 8k), which is consistent with enhanced innate immune signaling resulting from ORF9b deficiency. Conversely, stable over-expression of ORF9b in A549 lung epithelial cells led to a ∼45% downregulation of C15ORF48 protein under basal conditions without altering the levels of *C15ORF48* mRNA levels (Figure S7d-f, Supplemental Dataset 10). Treatment of wild type A549 cells with IL1Δ induced the expression of C15ORF48 protein nearly 20-fold in a manner that was dependent on the pre-existing expression of ORF9b. While IL1Δ induced *C15ORF48* mRNA levels > 50 fold, the presence of ORF9b did not influence the amount of *C15ORF48* mRNA accumulation (Figure S7f). In both cell lines, *C15ORF48* mRNA accumulation induced by IL1Δ was associated with a reduction in NDUFA4 protein but not mRNA, consistent with previous reports that miR147 encoded from this locus can control NDUFA4 stability^56,57,59^. NDUFA4 depletion was not influenced by ORF9b-dependent changes in C15ORF48 protein accumulation (Figure S7f), arguing that ORF9b expression influences the biogenesis of C15ORF48 protein but not mRNA under basal and inflammatory conditions. Taken together, these results demonstrate that ORF9b recruitment to mitochondria dampens subclinical innate immune signaling and OXPHOS transcriptional pathways, revealing an *in vivo* association between ORF9b and C15ORF48 that warrants future investigation.

## Discussion

We set out to study SARS-CoV-2 ORF9b because it had emerged as one of the most prominent mitochondrial interactors of any SARS-CoV-2 protein and a leading candidate antagonist of mitochondrial antiviral immunity, based largely on overexpression and interactome studies that placed it at the outer mitochondrial membrane (OMM) in complex with the import receptor TOMM70. By combining isogenic, single-copy inducible expression of ORF9b and its orthologues with reverse-genetic SARS-CoV-2 and authentic infection of human lung epithelial cells and golden Syrian hamsters, our study revises this view on several fronts. First, we show that the most robustly conserved property of ORF9b in human cells is not an effect on mitochondrial physiology but its own dependence on recruitment to TOMM70 for stability: docking at the receptor shields the viral protein from constitutive proteasomal degradation (Figs. 1–3). Second, we identify a single residue on TOMM70, E477, that is required for ORF9b recruitment yet entirely dispensable for the engagement of host mitochondria by *Toxoplasma gondii* (Fig. 4) and for the homeostatic functions of TOMM70 in complex assembly and metabolism (Fig. 5). Third, and most unexpectedly, deletion of ORF9b from the virus left replication, innate immune signaling, and lung pathology essentially intact (Figs. 7, 8), unmasking instead a discrete inflammatory respiratory-chain response characterized by an upregulation of C15ORF48 that is dependent on the presence of ORF9b. C15ORF48 is a nuclear-encoded gene whose expression is induced by infection and inflammation, leading to the production of a mitochondrial protein that displaces a NDUFA4, a structural subunit of Complex IV. *C15ORF48* also encodes miR-147b, which works synergistically to blunt NDUFA4 levels and Complex IV activity as well as mitigating antiviral Type I interferon (IFN) signaling^56–58^. Compelling evidence of this synergy was provided by the study of transgenic mouse models in which either C15ORF48 (referred to as NDUFA4L3) and/or miR147 (the mouse orthologue of miR147b) were genetically ablated in macrophages and the subsequent effects on NDUFA4 stability and innate immune signaling^59^. Here, we hypothesize that SARS-CoV-2 may induce immune evasion by regulating C15ORF48 biogenesis, NDUFA4, and consequently the composition of Complex IV. Complex IV contains 14 protein subunits and stands alone among ETC complexes in substituting its core subunits with closely related nuclear-encoded isoforms, which is a mechanism that lets it adjust activity according to extracellular signals^60–63^. Among these 14 subunits, C15ORF48 stands apart as the most significantly depleted protein by ORF9 expression in A549 cells (Supplemental Dataset 10). Collectively, these findings argue that ORF9b is a receptor-gated viral protein with a far more circumscribed role in mitochondrial and infection biology than the prevailing model has held.

The evolutionary conservation of ORF9b across sarbecoviruses need not imply functional essentiality: it is encoded in an alternative reading frame nested within the essential nucleocapsid (N) gene, its retention across sarbecoviruses may reflect purifying selection acting on N rather than selection for ORF9b function itself. Hence, it is not surprising that significant sequence divergence observed across ORF9b variants derived from bat and human sarbecoviruses (Fig. 2) had little impact on the recruitment and degradation kinetics of induced ORF9b in Hela cells. Why, then, has ORF9b come to be regarded as a dominant antagonist of innate immunity? We believe several non-exclusive factors reconcile our observations with the existing literature. First, most prior conclusions rest on comparative virology across variants whose host innate immune response differences cannot be formally attributed to ORF9b given concurrent mutations in spike and elsewhere^9,10,64^. Second, the induction of innate immune signaling and interferon stimulated genes (ISGs) is dynamic and complex and the heterogeneous expression of differential expressed genes may not converge to a signaling pathway appropriately encapsulated by a single ISG reporter construct or even a limited panel of ISGs^10,65^. Our bulk RNAseq studies of SARS-CoV-2-infected A549 cells (Fig. 7, S7) and Syrian hamster lungs (Fig. 8) highlight the complex and subtle changes in ISG and inflammatory gene expression rewiring that can be instigated by wild type and ORF9b-deficient viruses, respectively. Third, the transient overexpression of ORF9b in cell lines may yield supraphysiological levels of ORF9b that would engage protein quality control pathways that may lead us to misassign ORF9b-attributable effects. The single-copy inducible HeLa FTIR system was employed expressly to minimize these confounders, and our L52D and S53E binding-dead mutants of ORF9b are instructive in this regard as they prevent mitochondrial recruitment. A further and frequently conflated point is that TOMM70 is not an inert scaffold for ORF9b-recruitment. Beyond its defined role in mitochondrial protein biogenesis^66^, deletion or silencing of TOMM70 was itself sufficient to activate interferon-stimulated gene expression in the absence of infection and to remodel the host immune response in a manner distinct from ORF9b recruitment, and only a subset of the mitochondrial proteome changes caused by TOMM70 loss were reproduced by ORF9b expression (Figs. 5, 6). Phenotypes ascribed to an ORF9b–TOMM70 axis through perturbation of the receptor therefore cannot be assumed to report the activity of receptor-bound ORF9b and we have shown the two manipulations to be mechanistically and proteomically separable.

We detected ORF9b-dependent effects on innate signaling in a mitochondrial recruitment-dependent dampening of a subset of Sendai virus-induced host genes in HeLa cells, including *CXCL10* and *IFNB1* (Fig. 6), however these were quantitatively limited and were not recapitulated in authentic infection with SARS-CoV-2. In A549 cells and in the lungs of golden Syrian hamsters, deletion of ORF9b left viral replication, type I interferon and antiviral gene programs, mitochondrial proteome and morphology, lung viral titers, histopathology, and cell death essentially unchanged (Figs. 7, 8). The most parsimonious explanation is functional redundancy: SARS-CoV-2 encodes multiple, partially overlapping interferon antagonists and that loss of any single factor is buffered at the level of the intact virus. Cell-type context likely compounds this, as epithelial readouts may not capture the plasmacytoid dendritic cell and macrophage compartments that dominate type I interferon output *in vivo*, and a subclinical shift in signaling need not register as altered gross pathology at the peak of acute disease^65^. Consistent with a circumscribed rather than central role, we were moreover unable to recapitulate the proposed recruitment of IRF3 to the outer membrane upon Sendai virus infection (Fig. S5), suggesting that at least one mechanism advanced for ORF9b-mediated antagonism may not operate under the conditions we tested.

Perhaps the most striking outcome of deleting ORF9b *in vivo* was not the loss of an antiviral brake but the unmasking of a specific inflammatory respiratory-chain response. Transcriptomes of ORF9b-deficient virus-infected lungs were enriched for COVID-19, oxidative phosphorylation, and immune pathways, and among the handful of individually significant genes was *C15ORF48*, which was induced by wild-type SARS-CoV-2, further elevated in the absence of ORF9b (SARS-CoV-2^ΔORF9b^), and blunted by ectopic ORF9b expression (Fig. S8). C15ORF48 upregulation is well documented in COVID-19 patients^56,57^ and also elevated in cancer patients that exhibit increased survival and responsiveness to checkpoint inhibitor therapies^59^, presumably due to its strong anti-tumor immunity action. With the exception of the gut and lymphoid tissues, C15ORF48 protein and RNA expression is strikingly silent across other humans (https://www.proteinatlas.org/ENSG00000166920-C15orf48), which may reflect the role of C15ORF48 induction as a general pro-immunity response that rewires mitochondrial metabolism and immunity. We therefore propose a model in which ORF9b does not silence innate immunity wholesale but restrains the inflammation-driven remodeling of the respiratory chain, limiting induction of this NDUFA4-displacing modulator and thereby tuning the metabolic arm of the host response. This role is fully consistent with the circumscribed, mitochondrially focused phenotype we observe throughout and with the recent demonstration that chemically and genetically blunting OXPHOS function in lung epithelial cells enables a pro-viral response of SARS-CoV-2 *in vitro*^67^. We emphasize that this in relationship is presently correlative, resting on an *in vivo* association and ectopic blunting, and that its mechanistic basis remains to be defined in vivo. In A549 cells, ORF9b expression alone is sufficient to blunt the levels of C15ORF48 protein but not RNA both under basal and cytokine stimulation, arguing for a role in protein biogenesis. In vivo, however, the elevation of C15ORF48 could be a direct transcriptional consequence of ORF9b at the mitochondrial surface, a secondary effect of the dampened immune signaling we observe, or a downstream feature of the suppressed oxidative-phosphorylation program. Dissecting these possibilities and establishing whether the ORF9b–C15ORF48 axis influences disease outcome, is a priority that we are actively pursuing. Nevertheless, our data align with an emerging and unexplained view that dampening Complex IV assembly may facilitate innate immune suppression during infection through a mechanism that does not rely on viral RNA sensing^56,59^. Supporting this unexplained link between Complex IV deficiency and viral infection are studies of Complex IV-deficient mice and human patients. In mutant mice deficient for TACO1^68^, which is required to assembly Complex IV and whose deficiency in humans results in late-onset Leigh Syndrome^69^, cytomegalovirus infection exacerbates mitochondrial and physiological defects. In Leigh Syndrome patients carrying pathogenic mutations in the Complex IV assembly factor SURF1, viral infection is associated with metabolic decompensation and increased morbidity^70^. Yet whether ORF9b-mediated suppression of Complex IV contributes to the suppression of innate immunity *in vivo* remains to be defined.

Several boundaries delimit our conclusions. Our *in vivo* analysis examined a single permissive species of hamster, in males only at a single peak-disease timepoint and inoculum. Therefore, we do not address possible contributions of ORF9b at early infection time-points (e.g. several hours post-infection) nor to chronic and post-acute infection and re-infection. Second, as the SARS-CoV-2^ORF9bL52D^ virus expressing ORF9b with the binding-dead L52D point mutation also necessarily carries an obligate substitution at residue 55 of the overlapping nucleocapsid protein, which is not typically present in SARS-CoV-2 VoCs. Hence, the resulting post-infection results with the SARS-CoV-2^ORF9bL52D^ virus be interpreted with these two variants in mind, although the ORF9b-deleted virus SARS-CoV-2^ΔORF9b,^ which leaves the nucleocapsid intact, constitutes the unambiguous comparator throughout. We elected to generate these mutant viruses via reverse genetic engineering of the Wuhan (WA1) strain of SARS-CoV-2 because it expresses the highest amount of N/ORF9b RNA. However, we cannot exclude that ORF9b deletion in other VoCs or coronaviruses may have different effects, due to the differential expression and mutation of other viral factors. In addition, our structural inferences rest on AlphaFold predictions rather than experimental structures of the orthologue complexes. Finally, the small number of individually significant proteomic changes we report in infected cells should be regarded as hypothesis-generating. In summary, our study repositions SARS-CoV-2 ORF9b not as a master switch over mitochondrial antiviral immunity but as a receptor-gated viral protein whose principal measurable consequence during authentic infection is a restraint on inflammatory respiratory-chain remodeling. By resolving which effects belong to ORF9b, which to its receptor TOMM70, and which to viral infection itself, our work provides a framework for dissecting the expanding repertoire of pathogen factors that converge on the outer mitochondrial membrane. Our study establishes TOMM70, a single receptor engaged through distinct surfaces by virus and parasite alike, as a focal point at the intersection of mitochondrial biology, innate immunity, and host-directed antimicrobial strategy.

## Methods

### Molecular cloning and mutagenesis

Plasmids encoding C-terminal EGFP-tagged SARS-CoV-2 ORF9b wild type (EGFP-ORF9b^WT^) and L52D mutant (EGFP-ORF9b^L52D^) were obtained commercially (Addgene#165122,165129). These constructs were subcloned into the Gateway entry vector pDONR221 via BP recombination and cloned into a modified, gateway-compatible tetracycline-inducible destination vector pcDNA5/FRT/TO used previously^19^ by LR recombination. Human and bat EGFP-ORF9b constructs derived from SARS-CoV-1, BANAL-236, and RaTG13 were generated by gene synthesis (Thermo Fisher Scientific) directly in pDONR221 and gateway cloned into gateway-compatible pcDNA5/FRT/TO. To generate ORF9b^WT^-FLAG, pDONR223-SARS-CoV-2_ORF9b (Addgene#141280) was used as template to introduce an N-terminal FLAG epitope by site-directed mutagenesis using the Q5 site-directed mutagenesis kit (New England Biolabs) and the forward (5’gatgacgacaagTAATACCCAACTTTCTTGTACAAAG3’) and reverse (5’ gtctttgtagtcCTTCACGGTCACCACCAC3’) primers according to the manufacturer’s protocol. The S53E mutation in ORF9b^WT^-FLAG and was generated using site-directed mutagenesis with the forward (5’CAGCCCTCTGgaaCTGAACATGG3’) and reverse (5’ CCCAGTCTCAGGATGATAGG3’) primers to produce ORF9b^WT^-FLAG. Wild-type (pDONR223-ORF9b^WT^-FLAG) and mutant (pDONR223-ORF9b^S53E^-FLAG) entry clones were transferred into gateway-compatible pcDNA5/FRT/TO by LR recombination. Entry clones and destination constructs were verified by Sanger sequencing or whole-plasmid sequencing (Eurofins).

To generate human TOMM70 constructs, a codon-optimized *TOMM70* cDNA (Eurofins) was used as template. The *TOMM70* coding sequence was subcloned into the Gateway entry vector pDONR221 by BP recombination to generate the wild-type entry clone. Based on thepDONR221-TOMM70 wild-type entry clone, the E477A mutation was generated by site-directed mutagenesis using the forward (5’ GGGATATGGATATGCCCTGTATGCC3’) and reverse (5’ GCGGCACACGGCACATCGGGGAAATT 3’) primers and the D545A mutation was generated by site-directed mutagenesis using the forward (5’ TGCCTTCTTCGCCTATGAGACAATG 3’) and reverse (5’ CACTTATTCACACTTATTGTCGATTTC 3’) primers. Wild-type and mutant TOMM70 entry clones were transferred into a gateway-compatible lentiviral destination vector (pLV_Destination) under the control of a CMV promoter by LR recombination to generate lentiviral expression constructs. All entry clones and destination vectors were verified by Sanger sequencing prior to downstream applications.

### Generation of genetically modified inducible HeLa FlpN TRex cell lines

Flp-In T-REx HeLa (HeLa FITR) cells^19^ were used to generate isogenic tetracycline-inducible cell lines. pcDNA5/FRT/TO constructs encoding ORF9b variants (either N-terminal FLAG–tagged or C-terminal EGFP–tagged) were stably integrated into the single-copy locus following co-transfection with a Flp recombinase expression plasmid (pOG44, Invitrogen) using Lipofectamine 2000, according to the manufacturer’s instructions. Flp-mediated recombination enabled site-specific integration at the genomic FRT locus. Stable integrants were selected using hygromycin B (400 µg/mL) and blasticidin (15 µg/mL) until resistant populations were established. Resulting cell lines were maintained under dual antibiotic selection. ORF9b expression was induced by anhydrotetracycline (AHT; 10 ng/mL) and validated by immunoblotting and RT–qPCR prior to experimental use. Genetic disruption of *TOMM70* in HeLa FITR cells carrying either EGFP-ORF9b^WT^ or ORF9b^WT^-FLAG was achieved by CRISPR-based genome editing. A lentiviral lentiCRISPR plasmid encoding a sgRNA targeting human *TOMM70* (kind gift from Professor Lena Pernas) was used for gene editing^5^. Lentiviral particles were generated in HEK293T cells by co-transfection of the lentiCRISPR-sgTOMM70 plasmid together with packaging (pCMV-delta R8.2) and envelope (pCMV-VSV-G) plasmids using Lipofectamine 2000 (Thermo Fisher Scientific). Viral supernatants were collected 48 h post-transfection, filtered (0.45 μm), and applied to target cells. Stable integrants were selected with puromycin (2 μg/mL) for 7 days. TOMM70-deficienct clones were established by limiting dilution and validated by immunoblotting (anti-TOMM70; Proteintech, catalog 66593-1-IG) and deep sequencing of PCR products amplified with forward (5’ TCGTCGGCAGCGTCAGATGTGTATAAGAGACAGCGGGTGCCATATACCTGTG3’) and reverse (5’ GTCTCGTGGGCTCGGAGATGTGTATAAGAGACAGCAAATGCCCCACTCCCATCT3’) primers flanking the targeted genomic locus. To reconstitute TOMM70 expression in HeLa FITR *TOMM70^C^*^11^ and HeLa FITR *TOMM70^C^*^14^ mutant cells, lentiviral constructs encoding wild-type or mutant TOMM70 were transduced by using lentiviral particles. Because recipient cells were already resistant to puromycin (due to stable integration of lentiCRISPR-sgTOMM70), single-cell clones were isolated by fluorescence-activated cell sorting (FACS) and screened by immunoblotting using an anti-TOMM70 antibody to identify clones exhibiting comparable TOMM70 expression levels. To monitor mitophagy in real time, an mCherry–GFP iMLS mitophagy reporter construct (pLenti-III-PGK-Zeo-iMLS; gift from Anne Simonsen) was introduced into all ORF9b-inducible Flp-In T-REx HeLa cell lines described above by lentiviral transduction. Stable reporter-expressing populations were established by selection with Zeocin (15 µg/mL) and reported expression and correct mitochondrial localization were verified by fluorescence microscopy.

### Cell culture

HEK293T (293T) and Vero-E6 (ATCC, catalog.CRL-1586) cells were from the American Type Culture Collection and were cultured in growth medium consisting of DMEM 1× + GlutaMAX™ containing 4.5 g/L D-glucose and sodium pyruvate (Thermo Fisher Scientific, catalog. 10569010), supplemented with 5% fetal bovine serum (FBS) and 50 μg/mL penicillin–streptomycin, and maintained at 37 °C in a humidified atmosphere with 5% CO₂. HeLa Flp-In T-REx (HeLa FITR) were described previously and were cultured in DMEM 1× + GlutaMAX™ containing 4.5 g/L D-glucose and sodium pyruvate (Thermo Fisher Scientific, catalog. 10569010), supplemented with 10% fetal bovine serum (FBS) and 50 μg/mL penicillin–streptomycin, and maintained at 37 °C in a humidified atmosphere with 5% CO₂. A549 cells stably expressing human ACE2 (hACE2)^71^ and human TMPRSS2^72^ (hTMPRSS2), generated by sequential lentiviral transductions to yield A549^+hACE2+hTMPRSS2^ cells, which were cultured with 10 µg/mL blasticidin+100 µg/ml Hygromycin B, cultured in Ham’s F-12K (Kaighn’s) medium (Thermo Fisher Scientific, catalog. 21127030), supplemented with 10% fetal bovine serum (FBS) and 50 μg/mL penicillin–streptomycin and maintained at 37 °C in a humidified atmosphere with 5% CO₂. A549 cells stably expressing ORF9b (pLV-ORF9b^WT^) were generated in a similar manner. For experiments using HeLa FITR inducible cell lines, anhydrotetracycline (AHT; 10 ng/mL) was continuously maintained in the culture medium to sustain transgene expression, unless otherwise indicated. Stable expression of C15ORF48 using the previously validated pCDH hC15orf48-orf construct was stably expressed in HeLa Flp-In T-REx (HeLa FITR) carrying ORF9b^WT^-FLAG and single-cell clones were isolated by FACS. For experiments requiring galactose-based culture conditions, cells were cultured in glucose-free DMEM (Thermo Fisher Scientific, catalog 11966025) supplemented with galactose (4.5 g/L), sodium pyruvate, 10% FBS, and 50 μg/mL penicillin–streptomycin. Unless otherwise indicated, cells were allowed to adapt to the indicated metabolic conditions prior to downstream analyses. For proteasome inhibition studies, cells were treated with MG132 (1 µM; Sigma-Aldrich) or bortezomib (0.5 µM; Sigma-Aldrich), with DMSO used as vehicle control for the indicated duration. Cells were routinely screened for mycoplasma. Cells were authenticated by genotyping (Eurofins).

### Spinning disc fluorescence imaging and analyses

Cells were seeded in 96-well CellCarrier Ultra imaging plates (PerkinElmer) before imaging. Nuclei were labeled using NucBlue™ Live ReadyProbes™ Reagent (Thermo Fisher Scientific) and mitochondria were labeled with 100 nM Mito Tracker Deep Red(MTDR) and/or 100 nM tetramethyl rhodamine Ethyl Ester Perchlorate (TMRE) for 30 minutes at 37°C with 5% CO₂ before washout. Spinning disk confocal images were captured using the Operetta CLS or Opera Phenix High-Content Analysis systems (PerkinElmer) with 20× Water/1.0 NA or 63× Water/1.15 NA objectives. Automated image analysis was performed using Harmony5.1(PerkinElmer). Image processing and single-cell analysis were implemented using a predefined PhenoLOGIC™ supervised machine-learning workflow. Nuclei were identified using the “Find Nuclei” building block based on the Hoechst 33342 channel. Cell and mitochondrial segmentation were performed using the “Find Cytoplasm” building block using the MitoTracker Deep Red (MTDR) channel. The subcellular distribution of EGFP-tagged ORF9b was quantified on a per-cell basis relative to the segmented mitochondrial and cytoplasmic compartments. A supervised machine-learning classifier (PhenoLOGIC™) was trained to distinguish distinct patterns of ORF9b subcellular localization based on multiparametric image features, including signal intensity distribution and spatial relationships between EGFP-ORF9b and mitochondrial masks. Training datasets were generated by manual annotation of cells pooled from multiple time points, with approximately 50–100 cells per class, yielding a balanced training set across 4 classes. No explicit colocalization coefficients were applied; classification was based on supervised pattern recognition informed by manually curated examples. Following training, the classifier was applied to automatically classify non-training (unknown) cells at the single-cell level. Cells were categorized into four localization pattern classes—colocalized, saturated, weak, or cytosolic—as defined in the Results section. For each condition, cells from six technical replicate wells were analyzed, with a total of 491 to 1,823 cells quantified per condition. Classification output was expressed as the proportion of cells assigned to each category.

### Cell death assay

HeLa FITR cells were seeded in 96-well imaging plates (CellCarrier Ultra, PerkinElmer) and cultured for at least 24 h at 37 °C in a humidified 5% CO₂ atmosphere. On the day of the experiment, cells were incubated with NucBlue™ Live ReadyProbes™ reagent (Thermo Fisher Scientific) and propidium iodide (PI) (Sigma-Aldrich) and treated with the indicated cytotoxic stimuli. Cells were treated with 1 μM actinomycin D plus 10 μM ABT-737, 0.5 μM staurosporine, or 100 μM etoposide, with 0.3% DMSO used as a vehicle control, for the indicated durations. Live-cell imaging was performed every 1 h using an Operetta CLS High-Content Imaging System (PerkinElmer) equipped with a 20× Water/1.0 NA. NucBlue and PI were excited using 355–385 nm and 530–560 nm LED channels, respectively. Total cell numbers (NucBlue-positive nuclei) and dead cells (PI-positive) were quantified over time using Harmony 5.1 (PerkinElmer). Cell death was expressed as the ratio of PI-positive cells to total cells.

### Mitophagy assay and quantification

HeLa FITR cells harboring inducible ORF9b^WT^-FLAG or ORF9b^S53E^-FLAG and stably expressing the iMLS-QC mitophagy reporter were induced with anhydrotetracycline 24 hours prior mitophagy induction. To induce mitophagy, cells were treated with 100 μM deferiprone (DFP) for 24 hours and 0.1% DMSO was used as vehicle control. Cells were imaged using an Opera Phenix Plus High-Content Screening System (PerkinElmer) equipped with a 63× water-immersion objective (NA 1.15) under standard culture conditions (37 °C, 5% CO₂). Mitophagy was quantified by measuring the number of mCherry-only puncta per cell, image analysis and puncta quantification were performed using the default analysis algorithms implemented in the Harmony 5.1 (PerkinElmer).

### Immunofluorescence staining of fixed cells

Cells were seeded in 96-well CellCarrier Ultra imaging plates (PerkinElmer) and cultured under standard conditions. For immunofluorescence staining, cells were washed once with PBS and fixed with 4% paraformaldehyde (PFA) for 15 min at room temperature. After fixation, cells were washed three times with PBS and permeabilized with 0.1% Triton X-100 in PBS for 10 min at room temperature. Cells were washed and blocked in 10% fetal bovine serum (FBS) in PBS for at least 1 h at room temperature (or overnight at 4 °C). Primary antibodies diluted in 5% FBS in PBS were applied and incubated overnight at 4 °C. Following three washes with PBS containing 0.1% Tween-20 (PBST), cells were incubated with appropriate Alexa Fluor–conjugated secondary antibodies (Alexa Fluor 488, 568, or 647; diluted 1:1,000 in 5% FBS-PBS) for 1 h at room temperature in the dark. After washing three times with PBST, nuclei were counterstained with Hoechst 33342 (1:20,000 in PBS) for 5 min at room temperature, protected from light, followed by three final washes with PBS.

### Mitochondrial morphology image analysis

Image acquisition was performed using the Operetta CLS High-Content Analysis System (PerkinElmer) equipped with a 63× objective. Raw image measurements were exported for analysis. For mitochondrial quantification, total mitochondrial volume was calculated using a custom segmentation macro in Fiji (ImageJ), followed by image binarization. Mitochondrial morphology was assessed by quantifying the following parameters: mean area (Area), perimeter (Perim), circularity (Circ), solidity (Solid), Feret’s diameter (Feret), and minimum Feret’s diameter (MinFeret). Area refers to the surface of the selected region of interest, expressed in calibrated units (µm²). Perimeter corresponds to the total length of the outer boundary of the selection. Circularity is defined as 4π × area/perimeter², where a value of 1.0 indicates a perfect circle and values approaching 0 indicate increasingly elongated or irregular shapes. Solidity is calculated as the ratio of the selection area to its convex hull area (area/convex area), reflecting the degree of irregularity or concavity of the mitochondrial outline. Feret’s diameter corresponds to the longest distance between any two points along the selection boundary (also known as the caliper diameter), while minimum Feret’s diameter represents the shortest such distance, perpendicular to Feret’s diameter. For infected cells analysis, the overlap between the SARS-CoV-2 nucleocapsid antibody channel and the TOMM40 antibody channel was analyzed, enabling the quantification of mitochondria in cells positive for the SARS-CoV-2 nucleocapsid antibody and at the same time in cells lacking this marker. The macro is freely available on a public repository (https://github.com/jdhercam10/MitoShape) and will be updated regularly. Radar plots were generated using a custom Python script, in which all morphological descriptors were normalized to wild-type levels, such that wild-type values are represented as 1 and all other values are expressed relative to this reference.

### Live-cell confluence analysis by IncuCyte

Live-cell monitoring and confluence analysis were performed using an IncuCyte SX5 Live-Cell Analysis System. HeLa cells were seeded at a density of 2,000 cells per well in 96-well plates and allowed to adhere overnight. After complete attachment, cells were washed and cultured in either glucose-containing or galactose-containing medium before immediate transfer to the IncuCyte system. Phase-contrast images were acquired every 12 h for 5 consecutive days under standard culture conditions (37 °C, 5% CO₂). Cell confluence was quantified automatically using the default IncuCyte confluence algorithm. Each condition was analyzed using six technical replicate wells, and confluence values were used to assess cell growth dynamics under different metabolic conditions.

### Analysis of oxygen consumption rates

Oxygen consumption was measured with the XFe96 Analyzer (Seahorse Biosciences). 8000 HeLa FITR cells per well were seeded onto 96-well XFe96 cell culture plates. On the following day, cells were washed and incubated with Seahorse XF Base Medium completed on the day of the experiment with 1 mM Pyruvate, 2 mM Glutamine, and 10 mM Glucose. Cells were washed with the Seahorse XF Base Medium and incubated for 45 min in a 37°C non-CO2 incubator before starting the assay. Following basal respiration, cells were treated sequentially with: oligomycin 1 µM, CCCP 2 µM and Antimycin A 1 µM + 1 µM Rotenone (Sigma). Measurements were taken over 2-min intervals, proceeded by a 1-min mixing and a 30 s incubation. Three measurements were taken for the basal OCR, three for the non-phosphorylating OCR, three for the maximal OCR, and three for the extramitochondrial OCR. To account for variations in cell density, OCR data were normalized to cell confluence measured via the Incucyte SX5 Live-Cell Analysis System (Sartorius). Phase-contrast images were acquired for each well immediately prior to the Seahorse assay. The Integrated Confluence (percentage area covered by cells) was quantified using the Incucyte SX5 software segmentation algorithm.

### siRNA transfection

Gene silencing was performed using forward siRNA transfection with 20 nM siRNA and Lipofectamine RNAiMAX (Invitrogen). Cells were incubated for 72 h prior to analysis. Non-targeting control siRNA (Dharmacon, catalog. D-001220-01-05) and siRNAs targeting human TOMM70 (Dharmacon, catalog. L-054815-01-0005), TOMM40 (Dharmacon, catalog. M-012732-00-0005), MAVS (Dharmacon, catalog. M-024237-02-0005), and MARC2 (Dharmacon, catalog. M-018689-00-0005) were used as indicated. Downregulation efficiency was validated by RT-qPCR and/or immunoblot analyses were indicated.

### RT–qPCR

Total RNA was extracted using NucleoSpin RNA kit (MACHEREY-NAGEL). RNA concentration was measured using NanoQuant Plate™ (Infinite M200, TECAN), and 500 ng of total RNA was converted into cDNA using the iScript Reverse Transcription Supermix (Bio-Rad). RT–qPCR was performed using the CFX384 Touch Real-Time PCR Detection System (Bio-Rad) and SYBR® Green Master Mix (Bio-Rad) using the primers listed in Reagents and Tools Table. Human beta-actin and Hamster *Gapdh* were amplified as internal standards. Data were analyzed according to the 2^−ΔΔCT^ method^73^.

### Transcriptomic analyses by bulk RNAseq and NanoString

Total human RNA was isolated from snap-frozen cells by the RNeasy Kit (Qiagen, 74104). Hamster RNA extraction was performed using Direct-zol RNA MiniPrep w/ Zymo-Spin IIC Columns kit (OZYME catalog R2050) as previously described^49^. For bulk RNAseq studies, quality control was performed on an Agilent BioAnalyzer. Libraries were built using a TruSeq Stranded mRNA library Preparation Kit (Illumina, USA) following the manufacturer’s protocol. The A549 cells and hamster libraries were sequenced on two separate runs both on an Illumina NextSeq 2000 platform using paired-end 52b reads. The RNA-seq analysis was performed with Sequana 0.18.1^74^. In particular, we used the RNA-seq pipeline (version 0.20.0) (https://github.com/sequana/sequana_rnaseq) built on top of Snakemake 7.32.4^75^. Reads were trimmed from adapters using Fastp 0.23.2^76^ then mapped to the reference genome (GRCh38 from Ensembl for A549 cells and BCM_Maur_2.0 from NCBI for the hamster samples) using STAR 2.7.10a^77^. FeatureCounts 2.0.1 was used to produce the count matrix, assigning reads to features using corresponding annotations (GRCh38_113 for A549 cells and release 103 for BCM_Maur_2.0) with strand-specificity information^78^. Quality control statistics were summarized using MultiQC 1.16.0^79^. Clustering of transcriptomic profiles were assessed using a Principal Component Analysis (PCA). Due to the heterogeneity of variance observed within groups, differential expression testing was conducted using the voom method from the limma library 3.66.0^80,81^ indicating the significance (Benjamini-Hochberg adjusted p-values, false discovery rate FDR < 0.05) and the effect size (fold-change) for each comparison. Over-representation analysis (ORA) was performed to determine if genes modulated by genotype or infection are more present in specific pathways. ORA was performed on WebGestalt (https://www.webgestalt.org/). RNAseq data have been deposited at ENA with the dataset identifier E-MTAB-16841 (human) and E-MTAB-16842 (hamster). Functional enrichment analyses were performed with g:Profiler^82^, Enrichr^83^, and GSEA^84^.

To investigate SARS-CoV-2 expression in these datasets, a similar workflow was used using Bowtie 2.4.5 as mapper with the SARS-CoV-2 Wuhan reference genome from NCBI (GCF_009858895.2). Differential expression testing was performed with DESeq2 1.38.3.

NanoString gene expression analysis was performed using the nCounter XT Human Host Response CodeSet (Panel XT Hs Host Response CSO, catalog 115000449) together with the nCounter XT Master Kit (12 reactions; catalog 100052), following the manufacturer’s protocol. Briefly, equal amounts of total RNA were hybridized with reporter and capture probes at 65 °C for 16–18 h, and samples were processed and scanned on the nCounter Analysis System (NanoString Technologies). Raw RCC files were imported into nSolver Analysis Software (NanoString Technologies) for quality control and data processing. Quality control metrics, including imaging quality control, binding density, positive control linearity, and limit of detection, were evaluated using default thresholds recommended by the manufacturer. Background correction was performed using negative control probes, followed by two-step normalization based on internal positive control probes and housekeeping genes using the geometric mean method. Normalized gene expression values were used for downstream analyses.

### Isolation of mitochondria from human cells

Crude mitochondria were isolated from cultured HeLa FITR cells by differential centrifugation under cold conditions. All steps were performed at 4 °C unless otherwise indicated. Cells grown in 15-cm dishes were harvested by scraping in culture medium and pelleted by centrifugation at 600 × g for 5 min. Cell pellets were washed once with ice-cold PBS and centrifuged again at 600 × g for 5 min. Pellets were resuspended in homogenization buffer (20 mM HEPES–KOH, pH 7.4, 220 mM mannitol, 70 mM sucrose, 1 mM EGTA, 0.1% (w/v) BSA, supplemented with EDTA-free complete protease inhibitor cocktail) at a ratio of 1 mL buffer per 15-cm dish, and incubated on ice for 15 min. Cells were mechanically disrupted by repeated passage through a glass Potter homogenizer. Homogenates were centrifuged at 600 × g for 5 min to remove unbroken cells and nuclei. The supernatant was collected, and the pellet was resuspended in fresh homogenization buffer and subjected to a second round of homogenization and centrifugation. Supernatants from both rounds were pooled and centrifuged once more at 600 × g for 5 min to further clear nuclear debris. Mitochondria were pelleted by centrifugation at 8,000 × g for 10 min. The mitochondrial pellet was gently resuspended in homogenization buffer and centrifuged again at 8,000 × g for 10 min for washing. The final mitochondrial pellet was either processed immediately or stored at −80 °C. For storage, pellets were resuspended in homogenization buffer lacking BSA or in freezing buffer (300 mM trehalose, 10 mM HEPES–KOH, pH 7.4). Protein concentration was determined using the Bradford assay..

### SDS–PAGE and immunoblot analysis

Cells were lysed in ice-cold RIPA buffer (50 mM Tris–HCl, pH 7.4, 150 mM NaCl, 1% (v/v) Triton X-100, 0.1% SDS, 0.05% sodium deoxycholate, 1 mM EDTA) supplemented with a complete protease inhibitor cocktail (Roche). Lysates were incubated on ice for 30 min and clarified by centrifugation at 16,000 × g for 10 min at 4 °C. Protein concentrations were determined using the Bradford assay (Sigma-Aldrich) with BSA as standard, and absorbance was measured at 595 nm using an Infinite M2000 microplate reader (TECAN). Equal amounts of protein (20 μg) were mixed with 6× Laemmli sample buffer containing 2-mercaptoethanol, heated at 95 °C for 5 min, and resolved on 4–20% Mini-PROTEAN® TGX Stain-Free™ precast gels (Bio-Rad). Proteins were transferred onto nitrocellulose membranes using the Trans-Blot® Turbo™ Transfer System (Bio-Rad). Equal protein loading was verified by stain-free total protein detection. Membranes were blocked for 1 h at room temperature in 5% (w/v) non-fat dry milk in TBST (Tris-buffered saline, 0.1% Tween-20), then incubated overnight at 4 °C with primary antibodies diluted 1:1,000 in 2% (w/v) BSA in TBST. After washing, membranes were incubated for at least 1 h at room temperature with HRP-conjugated secondary antibodies diluted 1:10,000 in 5% milk. Chemiluminescent signals were developed using Clarity™ Western ECL substrate (Bio-Rad) and acquired with a ChemiDoc® Imaging System (Bio-Rad). Densitometric analysis was performed using Image Lab software (Bio-Rad).

### Blue native PAGE (BN-PAGE) of mitochondrial respiratory complexes

BN-PAGE was performed on crude mitochondrial fractions isolated from HeLa FITR cells. All steps were carried out at 4 °C unless otherwise indicated. Mitochondrial pellets were resuspended in Buffer 17 (30 mM HEPES, 150 mM potassium acetate, 12% (v/v) glycerol, 2 mM 6-aminohexanoic acid, 1 mM EDTA) and clarified by centrifugation (10,000 × g, 10 min). Protein concentration was determined by Bradford assay, and 25 μg mitochondrial protein was loaded per lane. Mitochondrial membranes were solubilized with ultra-pure digitonin using a digitonin-to-protein ratio of ≤6:1 (w/w) for 30 min on ice, followed by centrifugation (20,000 × g, 10 min) to remove insoluble material. The supernatant was supplemented immediately before loading with NativePAGE™ 5% G-250 Sample Additive (Thermo Fisher Scientific, catalog BN2004).Native protein complexes were resolved on NativePAGE™ 3–12% Bis-Tris gels (Thermo Fisher Scientific, catalog BN1001BOX) using an XCell SureLock Mini-Cell system with NativePAGE anode buffer and dark blue/light blue cathode buffers according to the manufacturer’s instructions. Proteins were transferred onto PVDF membranes using a Trans-Blot® Turbo system (Bio-Rad) with Bio-Rad transfer buffer. Membranes were fixed in 8% acetic acid for 5 min, destained with 100% methanol, blocked in 5% milk in TBST, and subjected to immunoblotting as described above. Antibody details are provided in Reagents and Tools Table.

### Structural modeling and visualization of the TOM70–ORF9b complex

Structural visualization of the TOM70–ORF9b complex was performed using PyMOL (Schrödinger, LLC). The experimentally determined structure of the human TOM70 in complex with SARS-CoV-2 ORF9b (Protein Data Bank PDB ID: 7DHG) was used as the reference model. Structural representations and residue-level interaction views were generated based on this structure.

To compare ORF9b variants from different viral origins, amino acid substitutions corresponding to SARS-CoV-1 ORF9b and RaTG13 ORF9b were manually introduced into the ORF9b sequence within the complex model using PyMOL mutagenesis tools. Side-chain conformations were selected based on steric compatibility and minimized locally when necessary. The resulting models were used for qualitative structural comparison. For predicted structures, AlphaFold-derived models of TOM70–ORF9b complexes were visualized and colored according to per-residue confidence scores (pLDDT), as indicated. Structural figures highlighting overall complex architecture, interaction interfaces, and residue-level contacts were rendered in PyMOL.

### Reverse genetic engineering of mutant ORF9b SARS-CoV-2

The recombinant viral genomes were constructed as previously described by de Melo et al., 2023^85^. Eleven overlapping fragments around 3 kb each and two fragments encoding for yeast-specific selection genes (His3 and Leu2) were assembled in the yeast centromere plasmid pRS416 to generate the complete genome of the recombinant virus. The T7 promotor was placed just before the 5’UTR sequence and a unique restriction site EAG1 was placed just after the poly(A) tail. Viral fragments were obtained either by RT-PCR on RNA extracted from Vero-E6 cells infected by SARS-CoV-2 using the SuperScript IV VILO Master Mix (11756050, Thermo Fisher Scientific) according to the manufacturer’s protocol or using synthetic genes (GeneArt). All PCR amplifications were performed using Phusion™ High-Fidelity DNA Polymerase (F530, Thermofisher). To produce recombinant viruses deleted or mutated in ORF9b (Figure 7), the PCR product of the fragment 10-1 which includes N and ORF9b protein sequence was cloned in Topo-TA vector, using TOPO™ TA Cloning™kit (K465001, Invitrogen) according to the manufacturer’s protocol. The deletion of ORF9b was achieved by mutating its start codon ATG to ACG in order not to alter the protein sequence of the N protein. A second start codon located 8 amino acids downstream of the initial one was also mutated in the same way to avoid the transcription of a shorter ORF9b. The leucine in position 52 of the ORF9b protein was mutated in aspartic acid (L52D) inducing a mutation in N protein sequence (A55G). Those mutations were introduced by Site-Directed Mutagenesis following Site-Directed Mutagenesis Phusion Kit manufacturer’s protocol (F541, Thermo Fisher Scientific). TOPO-TA-F10-1 plasmid was used as a template, and mutagenesis PCRs were performed with specific primers:

Del-ORF9b-For : AATCAGCGAAA**<u>C</u>**GCACCCCGCATTACG

Del-ORF9b-Rev : TTGGGGTCC**<u>G</u>**TTATCAGACATTTTAGTTTGTTCG

ORF9b-L52D-For : GTCTTGGTTCACCG**<u>GA</u>**CTCACTCAACATGG

ORF9b-L52D-Rev : TTGGGGTCCGTTATCAGACATTTTAGTTTGTTCG

Recombination of 11 overlapping viral DNA fragments and 3 yeast specific fragments (PRS416, Leucine-2 and Histidine-3) was performed into Saccharomyces cerevisiae BY4740 strand cultured for three days at 30 °C. For each recombination, subcultured clones were screened by Multiplex PCR Kit to verify the presence of the different fragments (data not shown) (206143, QIAGEN) according to the manufacturer’s protocol, using specific subsets of primer pairs as previously described [30]. To perform purification of a large amount of recombinant plasmid, the yeast colony of interest was cultured overnight in 200 mL of SD-His-Ura-Leu-medium at 30 °C under agitation (200 rpm). DNA extraction was processed using the NucleoBond Xtra Midi Plus kit (740422, Macherey–Nagel). Manufacturer’s protocol was followed with a beforehand lysing step in which harvested cells were incubated in 16 mL RES-Buffer complemented with 1.6 mg Zymolyase® 100-T (120493–1, AmsBio) and 160 μl of β-mercapthoethanol and incubated for 1 h at 37 °C before the addition of LYS-Buffer. The YAC plasmids containing viral cDNA of SARS-CoV-2^WT^, SARS-CoV-2^ΔORF9b^ or SARS-CoV-2^ORF9L52D^ constructs were digested at the unique restriction site located downstream of the 3′ end poly(A) tail using EagI-HF® enzyme (R3505, New England Biolabs) following the manufacturer’s protocol. The cDNA purified by a classical phenol–chloroform process was then transcribed *in vitro* using the T7 RiboMax™ Large Scale RNA Production System (P1300, Promega). The synthesized RNA was finally purified using a classical phenol–chloroform method, precipitated, and resuspended in Dnase/Rnase free water. Then, 12 μg of complete viral mRNA and 4 μg of pCMV plasmid encoding the viral nucleoprotein (N) gene were electroporated into 8.106 Vero-E6 cells (ATCC #CRL-1586) resuspended in 0.8 mL of Electroporation Solution (MIR50114, Mirus Bio™ Ingenio™) using the Gene Pulser Xcell Electroporation System (1652660, BioRad) with a pulse of 270 V and 950 μF. Cells were then transferred to a T75 culture flask with 12 mL of DMEM supplemented with 2% FCS (v/v) and cultured at 37 °C, 5% CO2 for several days until the cytopathic effect (CPE) was observed. The supernatant that corresponds to a P0 stock was harvested, aliquoted and frozen at −80 °C until titration. The viral sequence was controlled by NGS sequencing (Illumina).

### Viral stock titration

Sendai virus was obtained from American Type Culture Collection (ATCC, VR-105) and was not additionally sequence verified. For SARS-CoV-2, the titer of recombinant viruses from the P0 stock was determined by a lysis plaque assay. 1×10^6^ Vero-E6 cells were seeded into each well of 6-well plates and cultured overnight at 37 °C, 5% CO2. The virus was serially diluted in DMEM without FBS and 400 µl diluted viruses were transferred onto the monolayers. The viruses were incubated with the cells at 37 °C with 5% CO_2_ for 1 h. After the incubation, inoculum was removed, and an overlay medium was added to the infected cells per well. The overlay medium per well contained MEM 1X (prepared from MEM 10X (21430020, Gibco™), distilled water (15230162, Gibco™), L-glutamine (25030123, Gibco™) and gentamicin 10 mg/ml (15710064, Gibco™), 0.25% sodium phosphate (25080094, Gibco™), and 1X AVICEL (RC-581, Dupont). After a 4 to 6 days of incubation, the plates were stained with crystal violet (V5265, Merck) and lyses plaques were counted to assess virus titer. Next generation sequencing (NGS) of viral stocks was performed to confirm the absence of genetic revertants.

### Next-generation sequencing

Viral RNAs were extracted from 200 μl of viral stocks with TRIzol extraction and Rneasy mini kit (QIAGEN) elution, purified using Agencourt RNAClean XP beads (Beckman Coulter) at a ratio of 1.8, as recommended by the manufacturer, and eluted in 10 μl of nuclease-free water. Eight microliters of the purified RNA were then reverse-transcribed in cDNA using random hexamers (Invitrogen, Illkirch, France) and Superscript III reverse transcriptase (Invitrogen), according to the manufacturer’s instructions. Afterward, double-stranded DNA (dsDNA) was synthesized at 16°C for 2 hours in 80 μl of final reaction mixture, including 20 μl of cDNA, 8 μl of 10× Second Strand Reaction Buffer (NEB, Evry, France), 3 μl of deoxynucleotide triphosphate mix (10 mM; Invitrogen), 1 μl (10 U) of Escherichia coli DNA ligase (NEB), 4 μl (40 U) of E. coli DNA polymerase I (NEB), 1 μl (5 U) of E. coli ribonuclease H (NEB), and 43 μl of nuclease-free water. Last, dsDNA was purified using AMPure XP (Beckman Coulter) at a ratio of 1.8, as recommended by the manufacturer, and eluted in 20 μl of nuclease-free water.

The dsDNA libraries were constructed using the TruSeq DNA PCR-Free libraries prep kit (Illumina, San Diego, USA) and sequenced on Illumina NextSeq 2000 platform. The data obtained were analyzed using the Galaxy@Pasteur software^86^: The sequences are first assembled de novo, without any reference sequence, to obtain “contigs,” and then they are aligned on a reference sequence, making it possible to detect the nucleotide differences present in at least 2% of the cases.

### Virus infection in cultured cells

For the infection of HeLa cells (Sendai Virus, MOI =5), A549^+hACE2+hTMPRSS2^ (SARS-CoV-2, MOI = 0.001, 0.01 and 0.5), the virus was added directly to the cells and incubated for one hour at 37 °C with 5% CO_2_. Following this, the infected media was removed, and the cells were incubated in media without FBS under 37 °C with 5% CO_2_ conditions before being harvested for analysis at the indicated time points. All experiments involving the infection of SARS-CoV-2 were conducted within the biosafety level-3 (BSL-3) laboratory located at Institute Pasteur, following standard operating procedures.

### Virus growth kinetics in cultured cells

A549^+hACE2+hTMPRSS2^ cells were used to compare the replication kinetics of SARS-CoV-2 reference strain and ORF9b-deficient SARS-CoV-2^ΔORF9b^ and SARS-CoV-2^Orf9bL52D^ mutant viruses. On the day before infection, 8×10^6^ A549^+hACE2+hTMPRSS2^ cells were seeded into a T75 flask. Cells were washed once with PBS and inoculated with the different viruses in 4 mL of their respective cell culture media for 1h at MOI of 0.01 and 0.001. Inoculums were then removed, and 12 mL of their respective cell culture media were added to each flask. All cells were maintained at 37 °C in a humidified atmosphere in the presence of 5% CO2. Cell-culture supernatants were collected at the indicated time points after infection (12, 24, 48, and 72 hours post infection, hpi) and frozen at −80 °C. Viral titers were obtained by classical TCID50 method^87^ on Vero-E6 cells after 72 hpi. Experiments were performed in triplicate (n = 3).

### Human cell and parasite culture

Mycoplasma-free human foreskin fibroblasts (HFFs; ATCC CCL-171) and HeLa FITR cells (this study) were cultured at 37 °C under a humidified 5% CO₂ atmosphere in complete medium consisting of Dulbecco’s Modified Eagle Medium (DMEM) supplemented with 10% heat-inactivated fetal bovine serum (Invitrogen), HEPES buffer (pH 7.2), 2 mM GlutaMAX, 100 U/mL penicillin, and 50 µg/mL streptomycin. All culture reagents were from Gibco – Life Technologies (St Aubin, France). *Toxoplasma gondii* strains were maintained by serial passage on confluent HFF monolayers. Experiments were performed using the type I RH Δku80 strain expressing either cytosolic NanoLuc-GFP or the dense-granule protein GRA17 fused to an HA epitope. Both parasite lines were kindly provided by A. Bougdour (University of Grenoble-Alpes, France) and have been previously described^27,88^.

### Parasite infection in HeLa FITR cells

HeLa FITR cells were seeded on poly-L-lysine (Sigma-Aldrich P8920)–coated coverslips placed in 12-well plates and cultured in complete medium for about 48 h. Freshly egressed tachyzoites harvested from infected HFF monolayers were added to each well and centrifuged at low speed (300g, 3 min) to synchronize parasite entry. The infection progression was monitored by light microscopy and stopped by cold HBSS^++^ washes when an average of one to two parasites per host cell was observed (approximately 15 min post-contact). Infected cells were subsequently incubated for 16-18 h in complete medium at 37 °C under 5% CO₂.

### Live imaging

Prior to imaging, the culture medium of HeLa cells infected with RH Δku80 NanoLuc-GFP tachyzoites was removed and replaced with prewarmed HBSS^++^. After several washes, MitoTracker Deep Red was added to a final concentration of 50 nM in HBSS^++^ (from a 1 mM DMSO stock) and incubated for 20 min. Coverslips were mounted in Chamlide chambers (LCI Corp., Seoul, Korea) and imaged using an Eclipse Ti inverted confocal microscope (Nikon France Instruments, Champigny-sur-Marne, France) equipped with temperature- and CO₂-controlled stages (LCI Corp.). The microscope was fitted with a CSU-X1 spinning disk (Yokogawa), a 60x objective, and a sCMOS Prime camera (Photometrics). Image acquisition was controlled using MetaMorph software (Universal Imaging Corporation). All live imaging experiments were performed at 37 °C under 5% CO₂. Z-stack images were acquired with a step size of 0.2 µm using laser excitation at both l 488 nm and 642 nm. Laser intensity was kept to a minimum to reduce phototoxicity and prevent mitochondrial fragmentation.

### Expansion microscopy

The expansion microscopy (ExM) protocol was adapted from Pavlou *et al.* (2020)^89^. Briefly, samples were incubated overnight at 4 °C with 0.1 mg/mL Acryloyl-X in PBS in a humid chamber. Gelation was initiated at 4 °C for 10 min and completed at 37 °C for 50 min. Gels were digested in 3 mL of digestion buffer containing 0.5% Triton X-100 and 8 U Proteinase K (New England Biolabs, P8007S) for 30 min at 37 °C with gentle agitation. After digestion, gels were washed twice in PBS (15 min each) to remove residual Proteinase K. A 4-mm-diameter gel fragment was excised using a biopsy punch and expanded by immersion in three successive ddH₂O baths (5 mL each, 20 min per bath at RT). DNA staining was performed during the first bath using propidium iodide (2 µg/mL). Expanded gel pieces were immobilized on poly-L-lysine–coated coverslips. Excess liquid was removed while maintaining gel hydration using a humid Kimwipe placed on top. Imaging was performed using the same microscope setup as described for live imaging, with laser excitation at l 488 nm, 568 nm, and 642 nm using the live super-resolution (SR) module (Gataca systems).

### Image analysis and processing

Images were processed and analyzed using ImageJ. Hyperstacks were separated by channel, and each channel was deconvolved using the DeconvolutionLab2 plugin with the Richardson–Lucy algorithm and an appropriate point spread function (PSF). Ten iterations were used for live-cell imaging data and five iterations for ExM data. Channels were subsequently reassembled and resliced from 0.2 µm to 0.04 µm. Quantification was carried out manually by blinded visual inspection of live-cell imaging data. A parasitophorous vacuole membrane (PVM) was scored as positive when host-cell mitochondria exhibited a quasi-continuous alignment along the contours of the parasitophorous vacuole. Three-dimensional reconstructions were generated in ImageJ. ExM stacks containing PVM signals were duplicated, manually thresholded for optimal contrast, and converted to binary masks. Masks were expanded ten times using the dilate function to include approximately 1.6 µm surrounding the PVM. The Image Calculator function was then applied to mitochondrial stacks using the “multiply” option with the expanded mask. Resulting stacks were merged with the original PVM signal, resliced from 0.2 µm to 0.07 µm, and rendered in 3D using the 3D Viewer plugin. 360° rotation movies were generated, annotated, and a representative frame was selected for figure presentation (Supplemental Movie 1).

### Data Independent Acquisition-based Proteomics

#### Sample preparation

A549 and HeLa FITR cell pellets were lysed by sonication using a Covaris E220 instrument equipped with Adaptive Focused Acoustics technology (Covaris). Samples were l transferred to microTUBE-15 AFA Beads Screw-Cap tubes and sonicated using the following settings: 175 W peak incident power, 200 cycles per burst, 10% duty factor and 120 s processing time at 8 °C. Insoluble material was removed by centrifugation at 3,000 × g for 5 min at room temperature, and the clarified supernatants were transferred to clean low-binding microcentrifuge tubes. A549 and HeLa FITR cell lysates were then processed using a single-pot, solid-phase-enhanced sample preparation (SP3) workflow adapted from Hughes et al^90^. Briefly, protein disulfide bonds were reduced with 5 mM tris(2-carboxyethyl) phosphine (TCEP) for 30 min at room temperature. Free cysteine residues were alkylated with 20 mM iodoacetamide for 30 min at room temperature in the dark. Sera-Mag Carboxylate-Modified SpeedBeads, hydrophobic and hydrophilic particles (Cytiva; cat. nos. 45152105050250 and 65152105050250), were combined at a 1:1 ratio, washed three times with LC–MS-grade water and resuspended in water at 10 µg/µL. Beads were added to each lysate at a 10:1 bead-to-protein ratio. Protein binding to the paramagnetic beads was induced by addition of acetonitrile to a final concentration of 75% (v/v). Samples were mixed for 30 min at room temperature at 800 rpm in a ThermoMixer. Tubes were then placed on a magnetic rack until the bead–protein complexes had completely migrated to the tube wall, and the supernatant was carefully removed. Bead-bound proteins were washed twice with 80% acetonitrile in water and once with 100% acetonitrile. For each wash, beads were fully resuspended off the magnetic rack, collected again on the magnet, and the supernatant was discarded. Proteins were digested directly on the beads by addition of sequencing-grade modified trypsin at a 1:25 enzyme-to-protein ratio in 100 mM ammonium bicarbonate. Digestion was performed overnight at 37 °C with shaking at 1,000 rpm. The reaction was stopped by acidification with formic acid to a final concentration of 1% (v/v).

Following digestion, samples were briefly centrifuged at 3,000 × g for 1 min and placed on a magnetic rack to collect the beads. Peptide-containing supernatants were transferred to clean tubes. To maximize peptide recovery, beads were rinsed once with LC–MS-grade water, and the rinse was combined with the corresponding peptide fraction. Peptides were desalted using AssayMAP C18 cartridges, 5 µL bed volume, on a Bravo automated liquid-handling platform (Agilent Technologies). Cartridges were primed with 70% acetonitrile/0.1% trifluoroacetic acid and equilibrated with 0.1% trifluoroacetic acid in water. Peptide samples were loaded onto the cartridges, washed with 0.1% trifluoroacetic acid to remove residual salts and contaminants, and eluted with 70% acetonitrile/0.1% trifluoroacetic acid. Eluates were dried by vacuum centrifugation and resuspended in 2% acetonitrile/0.1% formic acid before LC–MS/MS analysis.

#### Liquid chromatography - Mass spectrometry LC-MS/MS analysis

Peptide mixtures were analyzed on a NanoElute 2 UHPLC system (Bruker Daltonics) coupled to a timsTOF Ultra 2 mass spectrometer (Bruker Daltonics) operated in DIA-PASEF mode. Peptides were separated on a PepSep Ultra C18 analytical column (25 cm × 75 μm, 1.5 μm; Bruker Daltonics) maintained at 50 °C. Chromatographic separation was performed at a flow rate of 0.25 µL/min using buffer A (0.1% FA in water) and buffer B (0.1% FA in ACN). The 30-min gradient consisted of: 2% to 4% B in 1 min, 4% to 20% B from 1 to 20 min, 20% to 25% B from 20 to 25 min, followed by an increase to 95% B in 1 min and a hold at 95% B for 4 min. The timsTOF Ultra 2 was operated in DIA-PASEF acquisition mode using 24 isolation windows of 25 m/z each, covering a precursor mass range of 400–1000 m/z. Instrument parameters were set as follows: capillary voltage, 1600 V; dry gas flow, 3 L/min; dry temperature, 200 °C. TIMS settings included an ion mobility range of 0.64– 1.45 V·s/cm² (1/k₀), with a ramp time of 100 ms and 100% duty cycle.

#### Protein identification and quantification

Raw MS data were processed in Spectronaut (v.20.2.250922.92449; Biognosys) using the directDIA mode with BGS Factory Settings. Searches were performed against the *Homo sapiens* UniProt protein database (*83,526 entries*) using the UniProt FASTA parsing rule. The directDIA+ (Deep) workflow with Pulsar Search was used for peptide identification. Carbamidomethylation of cysteine was set as a fixed modification, while protein N-terminal acetylation and methionine oxidation were specified as variable modifications (maximum of five per peptide). Enzyme specificity was set to Trypsin/P and LysC/P, allowing up to two missed cleavages, and peptide length was restricted to 7–52 amino acids. PSM, peptide, and protein group identifications were controlled at a 1% FDR. Precursor Q-value threshold was 0.01, and protein group Q-value cutoffs were set to 0.01 at the experiment level and 0.05 at the run level. Protein inference was performed using the IDPicker algorithm. Quantification was carried out at the MS2 level using peak area and the automatic LFQ strategy, selecting the top 1–3 peptides per protein group and 1–3 precursors per peptide. Cross-run normalization was applied using the automatic strategy with local (non-linear) regression for precision iRT calibration. Interference correction was enabled, requiring at least 2 MS1 and 3 MS2 data points. Dynamic extraction windows were used for both the m/z and ion mobility dimensions with a correction factor of 1.

#### Statistical analysis of mass spectrometry data

To identify proteins with differential abundance between conditions, quantified protein intensities exported from Spectronaut were analyzed. Only protein groups containing at least one peptide unique within the FASTA database (“unique peptide”) were retained. In addition, proteins were required to be quantified in at least two biological replicates of at least one of the two conditions being compared.

Proteins lacking intensity values in one condition (i.e., quantified exclusively in the other) were considered present in one condition and absent in the other. These proteins were set aside as differentially abundant by definition and displayed separately, ranked by their iBAQ (intensity-based absolute quantification) values^91^. For proteins quantified in both conditions, intensity values were log₂-transformed and normalized by median centering within each condition (section 3.5 in^92^. Missing values were imputed using the impute.slsa function of the R package imp4p^93^.

Differential abundance was assessed using a moderated *t*-test implemented in the limma R package^81^. Resulting *p*-values were corrected for multiple testing using an adaptive Benjamini–Hochberg procedure via the *adjust.p* function of the cp4p package^94^, employing the robust estimation method described in^95^ to infer the proportion of true null hypotheses. Proteins identified as statistically significant by this analysis, together with proteins present in only one condition, were reported as differentially abundant and visualized in volcano plots.

### Proteomics heatmap and module analysis

Protein-group quantifications of 7668 input protein groups were analyzed from the imputed and normalized dataset. They were filtered to ensure sufficient data for downstream inference and clustering. First, proteins were required to have observed replicates within each of the eight groups. Proteins with zero variance across observed values were excluded. After these basic filters, 7524 proteins remained from protein groups.

To identify proteins varying across the eight groups, we used *limma* moderated linear models with a group-only design matrix and an overall omnibus F-test for any group effect^81^. Empirical Bayes moderation with trend adjustment was applied. P-values were adjusted for multiple testing using the Benjamini–Hochberg false discovery rate (FDR) with a threshold of FDR < 0.01. To ensure biological relevance in addition to statistical significance, an effect-size filter was applied based on the range of per-group means:

Delta_i = max_g(m_{ig}) - min_g(m_{ig}),

where m_{ig} is the mean intensity of protein i in group g. Proteins were retained if Delta_i> 0.5, indicating a minimal effect-size along the groups. This effect-size criterion reduced the dataset to 2,634 proteins, which were used for module detection and heatmap visualization.

Modules of co-varying proteins were identified using Weighted Gene Co-expression Network Analysis (WGCNA)^37^. The abundance matrix was analyzed using a signed network based on robust biweight midcorrelation (bicor) computed with pairwise-complete observations. The soft-thresholding power was selected using the scale-free topology criterion (pickSoftThreshold; searched powers 1–12), the resulting power was 6. Networks were constructed and modules were detected using blockwiseModules with TOMType = “signed”, minimum module size of 30, deepSplit = 2, and module merging threshold mergeCutHeight = 0.25. In total, 12 modules were detected including the grey/unassigned set (module 0), 11 non-grey modules contained 2,469 proteins and the grey set contained 165 proteins. Module eigengenes (first principal component of each module) were computed excluding the grey module and used to summarize module-level patterns.

For visualization and clustering, protein abundances were standardized to protein-wise z-scores across samples. Let x_{ij} denote the intensity of protein i in sample j. For each protein i, the mean mu_i and standard deviation sigma_i were computed across samples with observed values, and z-scores were calculated as

z_{ij} = (x_{ij} - mu_i)/sigma_i

Heatmaps were generated with the ComplexHeatmap^96^ R package. Proteins were split by WGCNA module assignment, and hierarchical clustering was performed within each module slice using Pearson correlation distance (1 – r, computed using pairwise-complete observations) and complete linkage, with the corresponding dendrogram displayed for each module. Modules were ordered by similarity of their module-average z-score profiles across samples to facilitate interpretation of related module behaviors. Heatmap values were shown with a diverging color scale centered at zero.

### Principal component analysis (PCA)

Protein intensities obtained with the Spectronaut software (“PG.Quantity”) were log2-transformed. To correct for systematic intensity shifts between replicates within each condition, a within-condition median normalization was applied. For each condition, the sample median was computed using only proteins with no missing values in that condition; each sample profile was then shifted to the mean of the condition-specific medians, thereby aligning medians across replicates within the same condition while preserving between-protein differences (see section 3.5)^92^. The resulting normalized matrix was used for the PCA. The matrix was restricted to proteins that have at least one quantified value in each of the conditions. Next, missing values were imputed using maximum-likelihood estimation as implemented in the « imp4p » R package (« impute.mle » function) to finally get a matrix without missing values^93^. PCA was performed from the resulting matrix with the « prcomp » function in R using centering and unit-variance scaling, ensuring that each protein contributed equally to the decomposition irrespective of its absolute variance. The proportion of variance explained by each principal component was visualized with a scree plot for the first ten components. Sample scores were plotted for the first two principal components (PC1 vs PC2) that explained respectively around 22.94% and 18.05% of variance.

### Animal Housing and Viral Inoculation

Animal care was conducted in accordance with European animal welfare laws (Directive 2010/63/EU). The French Ministry of Research and local Animal Ethics Committees (CETEA 89) reviewed and approved all experiments under the authorized protocol (dap250050; APAFIS #56633-2025082618074677 v2. Male golden Syrian hamsters (*Mesocricetus auratus*) aged 5–6 weeks (body weight 60–80 g) were maintained under specific pathogen-free conditions in a Biosafety Level 3 facility. Animals were housed in individual isolators with *ad libitum* access to food and water and allowed a 1-week acclimation period prior to experimental manipulation. Animals were anesthetized with an intraperitoneal injection of 100 mg/kg ketamine (Imalgène 1000, Merial) and 5 mg/kg xylazine (Rompun, Bayer) and inoculated intranasally with 100 µL of physiological saline containing 6 x 10⁴ PFU of SARS-CoV-2 (50 µL per nostril). Mock-infected controls received physiological saline alone via the same route.

### Clinical Monitoring

Infected and mock-infected animals were housed in separate isolators to prevent cross-contamination. Daily clinical assessments were performed for 4 days post-infection, recording body weight and clinical score on a cumulative 0–4 scale reflecting observable signs of disease (ruffled fur, reduced locomotor activity, behavioral lethargy, and decreased exploratory behavior) as previously described^49^.

### Tissue Collection and Processing

At 4 days post-infection, animals were euthanized via intraperitoneal overdose of anesthetics (ketamine and xylazine) followed by exsanguination. Lung tissues were rapidly excised and immediately frozen at −80°C for downstream RNAseq and viral analyses. Briefly, frozen samples were weighed and transferred to Lysing Matrix M 2 mL tubes (116923050-CF, MP Biomedicals) containing 1 mL of ice-cold DMEM (Dulbecco’s Modified Eagle Medium, Gibco) supplemented with 1% penicillin/streptomycin (15140148, Thermo Fisher). Samples were homogenized using the FastPrep-24™ system (MP Biomedicals) with the following scheme: homogenization at 4.0 m/s for 20 s, incubation at 4 °C for 2 min, and a further homogenization at 4.0 m/s for 20 s. Tubes were centrifuged at 10,000 × g for 2 min at 4 °C, and the supernatants were collected and stored at −80 °C until further analysis. These supernatants were used for both RNA extraction and virus titration

#### RNA Extraction

Total RNA from lungs was extracted using the Direct-zol RNA MiniPrep kit (R2052, Zymo Research). Briefly, 125 µL of tissue homogenate was incubated with 375 µL of TRIzol LS (10296028, Invitrogen), and extraction was performed according to the manufacturer’s instructions, including DNase treatment.

#### Titration of SARS-CoV-2

Infectious viral titers were determined by TCID50 assay on the lung homogenate supernatants obtained after grinding (see Grinding of Lungs, above), following the same protocol used for the kinetic titration. Supernatants were serially diluted 1:10 in DMEM (Gibco) supplemented with 1% penicillin/streptomycin (15140148, Thermo Fisher) and 1 µg/mL trypsin-TPCK (4370285, Sigma-Aldrich), and 100 µL of each dilution was added to six wells of a 96-well plate (starting at 1:10). Next, 100 µL of a suspension containing 8 × 10⁵ Vero-E6 cells per mL in DMEM supplemented with 1% penicillin/streptomycin and 1 µg/mL trypsin-TPCK was added to each well. Plates were incubated at 37 °C and 5% CO2 for 3 days. Wells containing lysed cells and/or cytopathic effect (fused multinucleated cells, formation of syncytia) were scored using the 4× objective of an EVOS M5000 imaging system (Thermo Fisher). Viral titers were calculated by the classical TCID50 method using the TCID50 calculator, with a limit of quantification of 102 TCID50/mL.

#### Histology

The entire lung lobes were fixed in 10% neutral buffered formalin (VWR chemicals) for 24 hours and then transferred to 70% ethanol. The tissues were then embedded in paraffin and sectioned into 4 μm thick slices using a microtome. These sections were deparaffinized in xylene, rehydrated, and subsequently stained with hematoxylin and eosin (H&E) or TUNEL according to standard protocols. Immunohistochemical (IHC) staining to detect the virus was performed using an anti-SARS Nucleocapsid Protein antibody (NB100-56576, Novus Biologicals) and a biotinylated goat anti-rabbit Ig secondary antibody (E0432, Dako, Agilent). The images were captured using a slide scanner AxioScan.Z1 (ZEISS). Histopathological evaluation was performed on digitized whole-slide images using QuPath software^97^. All analysis settings and annotations were stored within the same project. The annotation tool was used to contour the entire lung lobe. For TUNEL quantification, positive-stained area was quantified within the total area, and the results were expressed as the percentage of positive-stained area. For damaged-area quantification in H&E-stained sections, the annotation tool was used to segment the damaged area within the contour of the entire lung lobe, and results were expressed as the percentage of the total area. The ratio of the damaged area to the total tissue area was calculated by three independent, blinded users and then represented.

### Statistical Analyses

Experiments were conducted at least three times, with quantitative analyses performed in a blinded manner. For high-throughput assessments (e.g., proteomics, immunoblots, qRT-PCR, imaging, RNAseq), all groups were measured in parallel to minimize experimental bias. Statistical analyses were conducted using GraphPad Prism v10 software, and data are presented as mean ± SD or SEM, as specified. Statistical tests and replicate numbers are detailed in the figure legends. Comparisons between two groups were made using an unpaired two-tailed T-test, while one-way or two-way ANOVA was used to compare more than two groups or groups across multiple time points. Exact P values are indicated.

### Graphics

Figures 1b, S1c, 5c, 7e, and 8c were created with Biorender.com. Figure S4a and S7a were created with Benchling.com. All other graphics were created with Adobe Illustrator.

## Data availability

All source data for the experiments are available with this manuscript. Proteomics datasets generated in this study have been deposited in the Proteomics Identification Database (PRIDE): HeLa cell proteomics (PRIDE identifier: PXD074755), A549 cell proteomics (PRIDE identifier: PXD074770). Transcriptomics datasets generated in this study for A549 cells (ENA E-MTAB-16841) and hamster lungs (E-MTAB-16842) have been deposited at EMBL-EBI: (https://www.ebi.ac.uk/biostudies/arrayexpress/studies). The processed and analyzed data for proteomics and RNAseq are included as Supplementary Datasets. Complete datasets for confocal imaging by Opera Phenix Plus HCS imaging are not available due to inoperability of exported images. However, these datasets are available upon request. Source data are included with this paper.

## Acknowledgments

We acknowledge Nassim Mahtal for imaging assistance, David Hardy for histology service, Élodie Turc and Georges Haustant from Biomics sequencing Platform, Victoria Pakulska for proteomics service (supported by the Agence Nationale de la Recherche ANR-21-CE35-0007), Zhengrui Zhang for AlphaFold-predicted structures and Protein interaction surface and Marie Lemesle for administrative assistance. We are grateful to all the members of the Wai, Schwartz, and Bouhry labs for helpful discussions. We thank Lena Ho for insights into C15ORF48 biology and for plasmid constructs. We thank Anne Simonsen for mitophagy constructs. TW is supported by the Agence Nationale de la Recherche (ANR-21-CE14 0052-02, ANR-23-CE13-0043-01). Part of this work was supported by the ANRS Emerging infectious diseases under the France 2030 program (ANRS-24-PEPRMIE-0006). The UTechS Photonic BioImaging, C2RT/DT, Institut Pasteur, is supported by the French National Research Agency (France BioImaging, ANR-24-INBS-0005 FBI (BIOGEN); Investments for the Future) and acknowledges Institut Pasteur and the Région Île-de-France (DIM1Health program) funding for the use of the Opera Phenix Plus and SiMA systems and the Tims TOF Ultra 2 (SLIMe Project). Biomics sequencing Platform, C2RT, Institut Pasteur, Paris, France, supported by France Génomique (ANR-10-INBS-09) and IBISA.

## References

1. Picard, M. & Shirihai, O. S. Mitochondrial signal transduction. Cell Metab. 34, 1620–1653 (2022).

2. Delgado, J. M. & Pernas, L. Mitochondria as sensors of intracellular pathogens. Trends Endocrinol. Metab. 0, (2024).

3. Pernas, L. et al. Toxoplasma effector MAF1 mediates recruitment of host mitochondria and impacts the host response. PLoS Biol. 12, e1001845 (2014).

4. Pernas, L., Bean, C., Boothroyd, J. C. & Scorrano, L. Mitochondria Restrict Growth of the Intracellular Parasite Toxoplasma gondii by Limiting Its Uptake of Fatty Acids. Cell Metab. 27, 886–897.e4 (2018).

5. Li, X. et al. Mitochondria shed their outer membrane in response to infection-induced stress. Science 375, eabi4343 (2022).

6. Defresne, T., Suspène, R. & Vartanian, J.-P. Cellular titanomachy: Viral forces clash with mitochondrial power. Annu. Rev. Virol. 12, 157–178 (2025).

7. Guarnieri, J. W. et al. Core mitochondrial genes are down-regulated during SARS-CoV-2 infection of rodent and human hosts. Sci. Transl. Med. 15, eabq1533 (2023).

8. Guarnieri, J. W. et al. Mitochondrial antioxidants abate SARS-COV-2 pathology in mice. Proc. Natl. Acad. Sci. U. S. A. 121, e2321972121 (2024).

9. Gordon, D. E. et al. Comparative host-coronavirus protein interaction networks reveal pan-viral disease mechanisms. Science 370, (2020).

10. Bouhaddou, M. et al. SARS-CoV-2 variants evolve convergent strategies to remodel the host response. Cell 0, (2023).

11. Gordon, D. E. et al. A SARS-CoV-2 protein interaction map reveals targets for drug repurposing. Nature 583, 459–468 (2020).

12. Ksiazek, T. G. et al. A novel coronavirus associated with severe acute respiratory syndrome. N. Engl. J. Med. 348, 1953–1966 (2003).

13. Jiang, H.-W. et al. SARS-CoV-2 Orf9b suppresses type I interferon responses by targeting TOM70. Cell. Mol. Immunol. 17, 998–1000 (2020).

14. Gao, X. et al. Crystal structure of SARS-CoV-2 Orf9b in complex with human TOM70 suggests unusual virus-host interactions. Nat. Commun. 12, 2843 (2021).

15. Ayinde, K. S., Pinheiro, G. M. S. & Ramos, C. H. I. Binding of SARS-CoV-2 protein ORF9b to mitochondrial translocase TOM70 prevents its interaction with chaperone HSP90. Biochimie 200, 99–106 (2022).

16. Lenhard, S. et al. The Orf9b protein of SARS-CoV-2 modulates mitochondrial protein biogenesis. J. Cell Biol. 222, (2023).

17. Batra, J., et al. Coronavirus protein interaction mapping in bat and human cells identifies molecular and genetic switches for immune evasion and replication. bioRxivorg (2025) doi:10.1101/2025.07.26.666918.

18. Moullan, N. et al. Tetracyclines Disturb Mitochondrial Function across Eukaryotic Models: A Call for Caution in Biomedical Research. Cell Rep. 10, 1681–1691 (2015).

19. Pujol, C. et al. MPC2 variants disrupt mitochondrial pyruvate metabolism and cause an early-onset mitochondriopathy. Brain 146, 858–864 (2022).

20. Miserey-Lenkei, S. et al. A comprehensive library of fluorescent constructs of SARS-CoV-2 proteins and their initial characterisation in different cell types. Biol. Cell 113, 311–328 (2021).

21. Mohanraj, K. et al. Inhibition of proteasome rescues a pathogenic variant of respiratory chain assembly factor COA7. EMBO Mol. Med. 11, (2019).

22. Temmam, S. et al. Bat coronaviruses related to SARS-CoV-2 and infectious for human cells. Nature 604, 330–336 (2022).

23. Zhou, P. et al. A pneumonia outbreak associated with a new coronavirus of probable bat origin. Nature 579, 270–273 (2020).

24. Jin, X. et al. Structural characterization of SARS-CoV-2 dimeric ORF9b reveals potential fold-switching trigger mechanism. Sci. China Life Sci. 66, 152–164 (2023).

25. Bichet, M. et al. The toxoplasma-host cell junction is anchored to the cell cortex to sustain parasite invasive force. BMC Biol. 12, 773 (2014).

26. Blank, M. L. et al. Toxoplasma gondii association with host mitochondria requires key mitochondrial protein import machinery. Proc. Natl. Acad. Sci. U. S. A. 118, e2013336118 (2021).

27. Gold, D. A. et al. The Toxoplasma dense granule proteins GRA17 and GRA23 mediate the movement of small molecules between the host and the parasitophorous vacuole. Cell Host Microbe 17, 642–652 (2015).

28. Wei, X. et al. Mutations in TOMM70 lead to multi-OXPHOS deficiencies and cause severe anemia, lactic acidosis, and developmental delay. J. Hum. Genet. 65, 231–240 (2020).

29. Dutta, D. et al. De novo mutations in TOMM70, a receptor of the mitochondrial import translocase, cause neurological impairment. Hum. Mol. Genet. 29, 1568–1579 (2020).

30. Dekker, P. J. T. et al. Preprotein translocase of the outer mitochondrial membrane: Molecular dissection and assembly of the general import pore complex. Mol. Cell. Biol. 18, 6515–6524 (1998).

31. Wiedemann, N. et al. Machinery for protein sorting and assembly in the mitochondrial outer membrane. Nature 424, 565–571 (2003).

32. Young, J. C., Hoogenraad, N. J. & Hartl, F. U. Molecular chaperones Hsp90 and Hsp70 deliver preproteins to the mitochondrial import receptor Tom70. Cell 112, 41–50 (2003).

33. Hill, K. et al. Tom40 forms the hydrophilic channel of the mitochondrial import pore for preproteins [see comment]. Nature 395, 516–521 (1998).

34. Backes, S. et al. The chaperone-binding activity of the mitochondrial surface receptor Tom70 protects the cytosol against mitoprotein-induced stress. Cell Rep. 35, 108936 (2021).

35. Hassdenteufel, S. et al. A multiplexed approach for genetic screening of human cells by electron microscopy uncovers a critical effector of mitochondrial cristae shape. Cell Biology (2025).

36. Chicherin, I. V. et al. Initiation factor 3 is dispensable for mitochondrial translation in cultured human cells. Sci. Rep. 10, 7110 (2020).

37. Langfelder, P. & Horvath, S. WGCNA: an R package for weighted correlation network analysis. BMC Bioinformatics 9, 559 (2008).

38. Saunders, N. et al. Dynamic label-free analysis of SARS-CoV-2 infection reveals virus-induced subcellular remodeling. Nat. Commun. 15, 4996 (2024).

39. Ng, M. Y. W., Wai, T. & Simonsen, A. Quality control of the mitochondrion. Dev. Cell 56, 881–905 (2021).

40. Munson, M. J. et al. GAK and PRKCD are positive regulators of PRKN-independent mitophagy. Nat. Commun. 12, 6101 (2021).

41. Allen, G. F. G., Toth, R., James, J. & Ganley, I. G. Loss of iron triggers PINK1/Parkin-independent mitophagy. EMBO Rep. 14, 1127–1135 (2013).

42. Han, L. et al. SARS-CoV-2 ORF9b antagonizes type I and III interferons by targeting multiple components of the RIG-I/MDA-5-MAVS, TLR3-TRIF, and cGAS-STING signaling pathways. J. Med. Virol. 93, 5376–5389 (2021).

43. Walsh, D. & Mohr, I. Viral subversion of the host protein synthesis machinery. Nat. Rev. Microbiol. 9, 860–875 (2011).

44. Labbé, K. et al. Specific activation of the integrated stress response uncovers regulation of central carbon metabolism and lipid droplet biogenesis. Nat. Commun. 15, 8301 (2024).

45. Medeiros, T. C. et al. Mitochondria protect against an intracellular pathogen by restricting access to folate. Science 389, eadr6326 (2025).

46. Shen, B. et al. Proteomic and metabolomic characterization of COVID-19 patient Sera. Cell 182, 59–72.e15 (2020).

47. Laudenbach, B. T. et al. NUDT2 initiates viral RNA degradation by removal of 5’-phosphates. Nat. Commun. 12, 6918 (2021).

48. Sia, S. F. et al. Pathogenesis and transmission of SARS-CoV-2 in golden hamsters. Nature 583, 834–838 (2020).

49. de Melo, G. D., et al. Attenuation of clinical and immunological outcomes during SARS-CoV-2 infection by ivermectin. EMBO Mol. Med. 13, e14122 (2021).

50. Yuan, L. et al. Gender associates with both susceptibility to infection and pathogenesis of SARS-CoV-2 in Syrian hamster. Signal Transduct. Target. Ther. 6, 136 (2021).

51. Ackermann, M. et al. Pulmonary vascular endothelialitis, thrombosis, and angiogenesis in Covid-19. N. Engl. J. Med. 383, 120–128 (2020).

52. Norooznezhad, A. H. & Mansouri, K. Endothelial cell dysfunction, coagulation, and angiogenesis in coronavirus disease 2019 (COVID-19). Microvasc. Res. 137, 104188 (2021).

53. Pinto, S. M. et al. Multi-OMICs landscape of SARS-CoV-2-induced host responses in human lung epithelial cells. iScience 26, 105895 (2023).

54. Masud, S. N. et al. Chemical genomics with pyrvinium identifies C1orf115 as a regulator of drug efflux. Nat. Chem. Biol. 18, 1370–1379 (2022).

55. Lau, M.-T. et al. Systematic functional identification of cancer multi-drug resistance genes. Genome Biol. 21, 27 (2020).

56. Lee, C. Q. E. et al. Coding and non-coding roles of MOCCI (C15ORF48) coordinate to regulate host inflammation and immunity. Nat. Commun. 12, 2130 (2021).

57. Clayton, S. A. et al. Inflammation causes remodeling of mitochondrial cytochrome c oxidase mediated by the bifunctional gene C15orf48. Sci. Adv. 7, eabl5182 (2021).

58. Takakura, Y. et al. Mitochondrial protein C15ORF48 is a stress-independent inducer of autophagy that regulates oxidative stress and autoimmunity. Nat. Commun. 15, 953 (2024).

59. Clark, M. L. et al. Mitochondrial complex IV remodeling in tumor-associated macrophages amplifies interferon signaling and promotes anti-tumor immunity. Immunity 58, 1670–1687.e12 (2025).

60. Pierron, D. et al. Cytochrome c oxidase: evolution of control via nuclear subunit addition. Biochim. Biophys. Acta 1817, 590–597 (2012).

61. Mell, O. C., Seibel, P. & Kadenbach, B. Structural organisation of the rat genes encoding liver-and heart-type of cytochrome c oxidase subunit VIa and a pseudogene related to the COXVIa-L cDNA. Gene 140, 179–186 (1994).

62. Sinkler, C. A. et al. Tissue- and condition-specific isoforms of mammalian cytochrome c oxidase subunits: From function to human disease. Oxid. Med. Cell. Longev. 2017, 1534056 (2017).

63. Fukuda, R. et al. HIF-1 regulates cytochrome oxidase subunits to optimize efficiency of respiration in hypoxic cells. Cell 129, 111–122 (2007).

64. Bouhaddou, M. et al. The Global Phosphorylation Landscape of SARS-CoV-2 Infection. Cell 182, 685–712.e19 (2020).

65. Ravindra, N. G. et al. Single-cell longitudinal analysis of SARS-CoV-2 infection in human airway epithelium identifies target cells, alterations in gene expression, and cell state changes. PLoS Biol. 19, e3001143 (2021).

66. Wiedemann, N. & Pfanner, N. Mitochondrial Machineries for Protein Import and Assembly. Annu. Rev. Biochem. 86, 685–714 (2017).

67. Soto Albrecht, Y. E., et al. Mitochondrial OXPHOS restricts SARS-CoV-2 replication. Sci. Adv. 12, eadz3081 (2026).

68. Richman, T. R. et al. Loss of the RNA-binding protein TACO1 causes late-onset mitochondrial dysfunction in mice. Nat. Commun. 7, 11884 (2016).

69. Weraarpachai, W. et al. Mutation in TACO1, encoding a translational activator of COX I, results in cytochrome c oxidase deficiency and late-onset Leigh syndrome. Nat. Genet. 41, 833–837 (2009).

70. Sofou, K. et al. A multicenter study on Leigh syndrome: disease course and predictors of survival. Orphanet J. Rare Dis. 9, 52 (2014).

71. Buchrieser, J. et al. Syncytia formation by SARS-CoV-2-infected cells. EMBO J. 39, e106267 (2020).

72. Saunders, N. et al. TMPRSS2 is a functional receptor for human coronavirus HKU1. Nature 624, 207–214 (2023).

73. Livak, K. J. & Schmittgen, T. D. Analysis of relative gene expression data using real-time quantitative PCR and the 2(−Delta Delta C(T)) Method. Methods 25, 402–408 (2001).

74. Cokelaer, T., Desvillechabrol, D., Legendre, R. & Cardon, M. “Sequana”: a Set of Snakemake NGS pipelines. J. Open Source Softw. 2, 352 (2017).

75. Köster, J. & Rahmann, S. Snakemake--a scalable bioinformatics workflow engine. Bioinformatics 28, 2520–2522 (2012).

76. Chen, S., Zhou, Y., Chen, Y. & Gu, J. fastp: an ultra-fast all-in-one FASTQ preprocessor. Bioinformatics 34, i884–i890 (2018).

77. Dobin, A. et al. STAR: ultrafast universal RNA-seq aligner. Bioinformatics 29, 15–21 (2013).

78. Liao, Y., Smyth, G. K. & Shi, W. featureCounts: an efficient general purpose program for assigning sequence reads to genomic features. Bioinformatics 30, 923–930 (2014).

79. Ewels, P., Magnusson, M., Lundin, S. & Käller, M. MultiQC: summarize analysis results for multiple tools and samples in a single report. Bioinformatics 32, 3047–3048 (2016).

80. Law, C. W., Chen, Y., Shi, W. & Smyth, G. K. voom: Precision weights unlock linear model analysis tools for RNA-seq read counts. Genome Biol. 15, R29 (2014).

81. Ritchie, M. E. et al. limma powers differential expression analyses for RNA-sequencing and microarray studies. Nucleic Acids Res. 43, e47 (2015).

82. Reimand, J. et al. Pathway enrichment analysis and visualization of omics data using g:Profiler, GSEA, Cytoscape and EnrichmentMap. Nat. Protoc. 14, 482–517 (2019).

83. Chen, E. Y. et al. Enrichr: interactive and collaborative HTML5 gene list enrichment analysis tool. BMC Bioinformatics 14, 128 (2013).

84. Subramanian, A. et al. Gene set enrichment analysis: a knowledge-based approach for interpreting genome-wide expression profiles. Proc. Natl. Acad. Sci. U. S. A. 102, 15545–15550 (2005).

85. de Melo, G. D. et al. Neuroinvasion and anosmia are independent phenomena upon infection with SARS-CoV-2 and its variants. Nat. Commun. 14, 4485 (2023).

86. Galaxy Community. The Galaxy platform for accessible, reproducible, and collaborative data analyses: 2024 update. Nucleic Acids Res. 52, W83–W94 (2024).

87. Lindenbach, B. D. Measuring HCV infectivity produced in cell culture and in vivo. Methods Mol. Biol. 510, 329–336 (2009).

88. Bellini, V. et al. Target identification of an antimalarial oxaborole identifies AN13762 as an alternative chemotype for targeting CPSF3 in Apicomplexan parasites. iScience 23, 101871 (2020).

89. Pavlou, G. et al. Coupling polar adhesion with traction, spring, and torque forces allows high-speed helical migration of the protozoan parasite Toxoplasma. ACS Nano 14, 7121–7139 (2020).

90. Hughes, C. S., Sorensen, P. H. & Morin, G. B. A standardized and reproducible proteomics protocol for bottom-up quantitative analysis of protein samples using SP3 and mass spectrometry. Methods Mol. Biol. 1959, 65–87 (2019).

91. Schwanhäusser, B. et al. Global quantification of mammalian gene expression control. Nature 473, 337–342 (2011).

92. Giai Gianetto, Q. Statistical analysis of post-translational modifications quantified by label-free proteomics across multiple biological conditions with R: Illustration from SARS-CoV-2 infected cells. Methods Mol. Biol. 2426, 267–302 (2023).

93. Gianetto, Q. G., Wieczorek, S., Couté, Y. & Burger, T. A peptide-level multiple imputation strategy accounting for the different natures of missing values in proteomics data. bioRxiv (2020) doi:10.1101/2020.05.29.122770.

94. Giai Gianetto, Q., et al. Calibration plot for proteomics: A graphical tool to visually check the assumptions underlying FDR control in quantitative experiments. Proteomics 16, 29–32 (2016).

95. Pounds, S. & Cheng, C. Robust estimation of the false discovery rate. Bioinformatics 22, 1979–1987 (2006).

96. Gu, Z., Eils, R. & Schlesner, M. Complex heatmaps reveal patterns and correlations in multidimensional genomic data. Bioinformatics 32, 2847–2849 (2016).

97. Bankhead, P. et al. QuPath: Open source software for digital pathology image analysis. Sci. Rep. 7, 16878 (2017).

