## Supplemental Information for "SARS-CoV-2 ORF9b exploits mitochondrial recruitment to TOMM70 for proteasomal protection and tunes inflammatory remodeling during lung infection"

**Supplemental Figures 1-7**

**Supplemental Datasets 1-10**
Supplemental Dataset 1 – Proteomics of HeLa FITR cells
Supplemental Dataset 2 – Nanostring of HeLa cels infected with Sendai Virus
Supplemental Dataset 3 – Nanostring of A549 cells infected with SARS-CoV-2
Supplemental Dataset 4 – RNAseq of A549 cells infected with SARS-CoV-2
Supplemental Dataset 5 – Pathway enrichment of RNAseq of A549 cells
Supplemental Dataset 6 – Proteomics of A549 cells infected with SARS-CoV-2
Supplemental Dataset 7 - Pathway enrichment of Proteomics of A549 cells
Supplemental Dataset 8 – RNAseq of Hamsters infected with SARS-CoV-2
Supplemental Dataset 9 – Pathway enrichment of RNAseq of Hamsters
Supplemental Dataset 10 - Proteomics of A549 cells expressing SARS-CoV-2 ORF9b

**Supplemental Tables 1-5**
Supplemental Table 1 – Comparisons of mitochondrial ORF9b accumulation in Figure 2c
Supplemental Table 2 – Comparisons of mitochondrial ORF9b reduction in Figure 2e
Supplemental Table 3 – NGS characterization of TOMM70-deficient HeLa FITR cell lines
Supplemental Table 4 – NGS determination of SARS-CoV-2 viral sequences in A549 cells
Supplemental Table 5 – NGS determination of SARS-CoV-2 viral sequences in Hamster lungs

**Supplemental Movie 1**

**Reagents and Tools Table**

**Supplemental Figure 1**

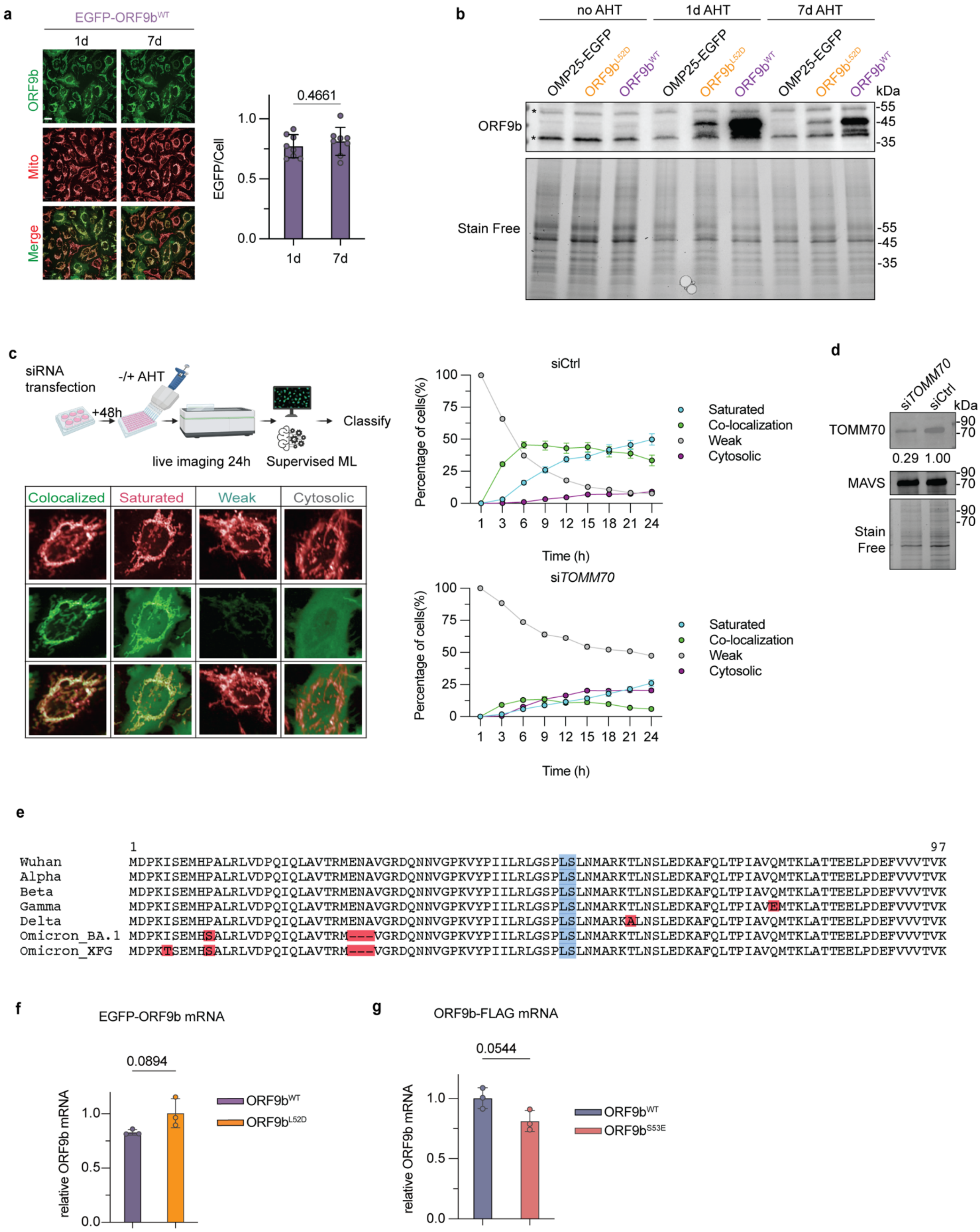


**Supplemental Figure 1: ORF9b recruitment limits its proteasomal degradation**

a) (Left) Representative images of live imaging of HeLa FITR induced with AHT to express EGFP-ORF9b$^{WT}$ (ORF9b$^{WT}$ + AHT) for 1 or 7 days with AHT (10ng/mL). Mitochondria were labeled with Mitotracker DeepRed (MTDR). Scale bar=20 µm. (Right) Single-cell quantification of EGFP intensity (EGFP/Cell) determined with Harmony 5.1. Data are means ± SD of 8 replicates with 285-505 cells measured per replicate analyzed by 2-tailed unpaired Student's t test. P-values are indicated.

b) Western blots of lysates from HeLa FITR induced with AHT to express EGFP-ORF9b$^{WT}$ (ORF9b$^{WT}$, purple), EGFP- ORF9b$^{L52D}$ (ORF9b$^{L52D}$, orange) for 1 or 7 days with AHT (10ng/mL) using anti-ORF9b antibody. HeLa FITR cells stably expressing OMP25-EGFP was used as a negative control. * indicates non-specific immunoreactive bands.
c) Representative images of supervised machine learning classification and quantification EGFP-ORF9b recruitment pattern in cells using Harmony 5.1 (left). Mitochondria were labeled with Mitotracker DeepRed (MTDR). Representative images of the four classes: *colocalized* (green) *saturated* (orange), *weak* (blue) and *cytosolic* (purple) trained on > 200 cells are indicated below. Scale bar=20 μm. Benchmarking of classification system in HeLa FITR EGFP-ORF9b$^{WT}$ downregulated for TOMM70 (si*TOMM70*) or negative control (siCTRL) for 48 hours prior to the induction of EGFP- ORF9b$^{WT}$ with AHT. Data are means ± SEM of 6 replicates with 256-1823 cells measured per replicate.
d) Western blot of lysates from siCTRL and si*TOMM70* cells in Figure S1c probed with the indicated antibodies. Densitometric quantification of TOMM70 is relative to stain-free.
e) Protein sequence alignment of ORF9b from SARS-CoV-2 Wuhan and subsequent variants that emerged from 2020 to 2025 (Alpha, Beta, Gamma, Delta, Omicron_BA.1, and Omicron_XFG), revealing conservation of the L52 and S53 amino acid residues highlighted in blue. Sequence variations are highlighted in red.
f) Measurement of ORF9b mRNA expression by RT-qPCR after 24h induction of EGFP-ORF9b$^{WT}$ (ORF9b$^{WT}$, purple, n=3), or mutant EGFP-ORF9b$^{L52D}$ (ORF9b$^{L52D}$, orange, n=3), expression with anhydrotetracycline (AHT, 10ng/mL). Normalization according to *GAPDH*. Data are means ± SD, 2-tailed unpaired Student's t test. P-value is indicated.
g) Measurement of ORF9b mRNA expression by RT-qPCR after 24h induction of ORF9b$^{WT}$-FLAG (ORF9b$^{WT}$, purple, n=3), or mutant ORF9b$^{S53E}$-FLAG (ORF9b$^{S53E}$, sky blue, n=3), expression with anhydrotetracycline (AHT, 10ng/mL). Normalization according to *GAPDH*. Data are means ± SD, 2-tailed unpaired Student's t test. P-value is indicated.

**Suplemental Figure 2**

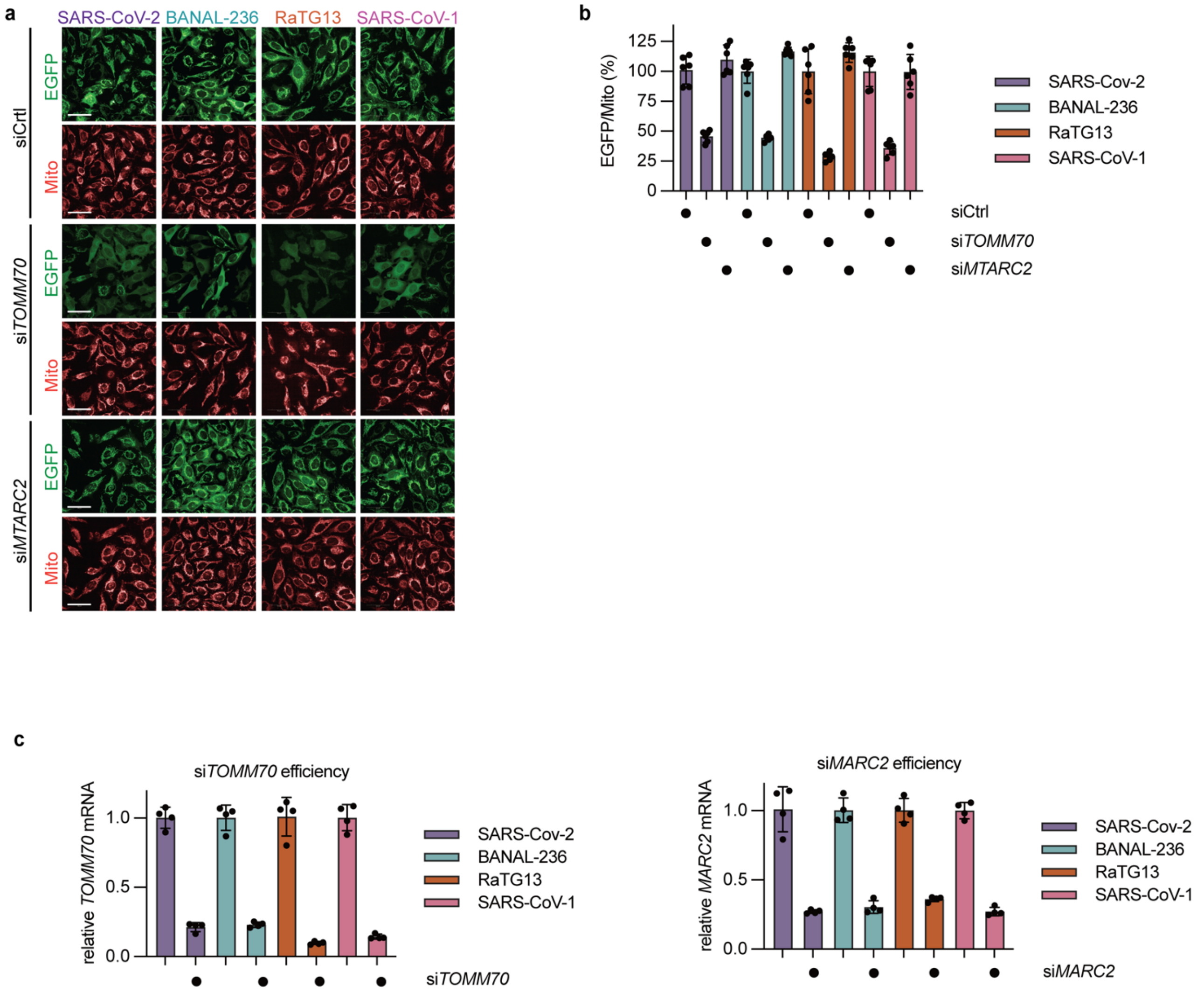


**Supplemental Figure 2: Contribution of TOMM70 and MARC2 on ORF9b recruitment**

a) Representative images of live imaging of HeLa Flp-In™ T-REx™ (HeLa FITR) cells expressing EGFP-ORFb9 variants from SARS-CoV-2 (purple, n=7), BANAL-236 (pink, n=7), RaTG13 (orange, n=7), and SARS-CoV-1 (teal, n=7) with anhydrotetracycline (AHT, 10ng/mL) for indicated time points. Knockdown of *TOMM70* (si*TOMM70,* n=6) , *MARC2* (si*MARC2*, n=6) or negative control (siCTRL, n=6) was performed 48 hours prior to the induction of EGFP- ORFb9$^{WT}$ with AHT. Mitochondria (Mito) were labeled with Mitotracker DeepRed. Scale bar=20 μm.

b) EGFP intensity in Supplemental Figure 2a was quantified using Harmony 5.1 as integrated EGFP fluorescence normalized to mitochondrial area. For each cell line, transfected with the indicated siRNAs, EGFP values were normalized to the siCtrl condition at 21 h after AHT addition, which was set to 100%. Data are shown as relative EGFP recruitment (% of control at 21 h) and represent means ± SEM of four replicates with 245–1061 cells measured per replicate.

c) Measurement of *TOMM70* (n=4) and *MARC2* (n=4) mRNA expression by RT-qPCR after 48h siRNA-mediated knockdown in Supplemental Figure 2a. Normalization according to β-*actin*. Data are means ± SD.

**Supplemental Figure 3**

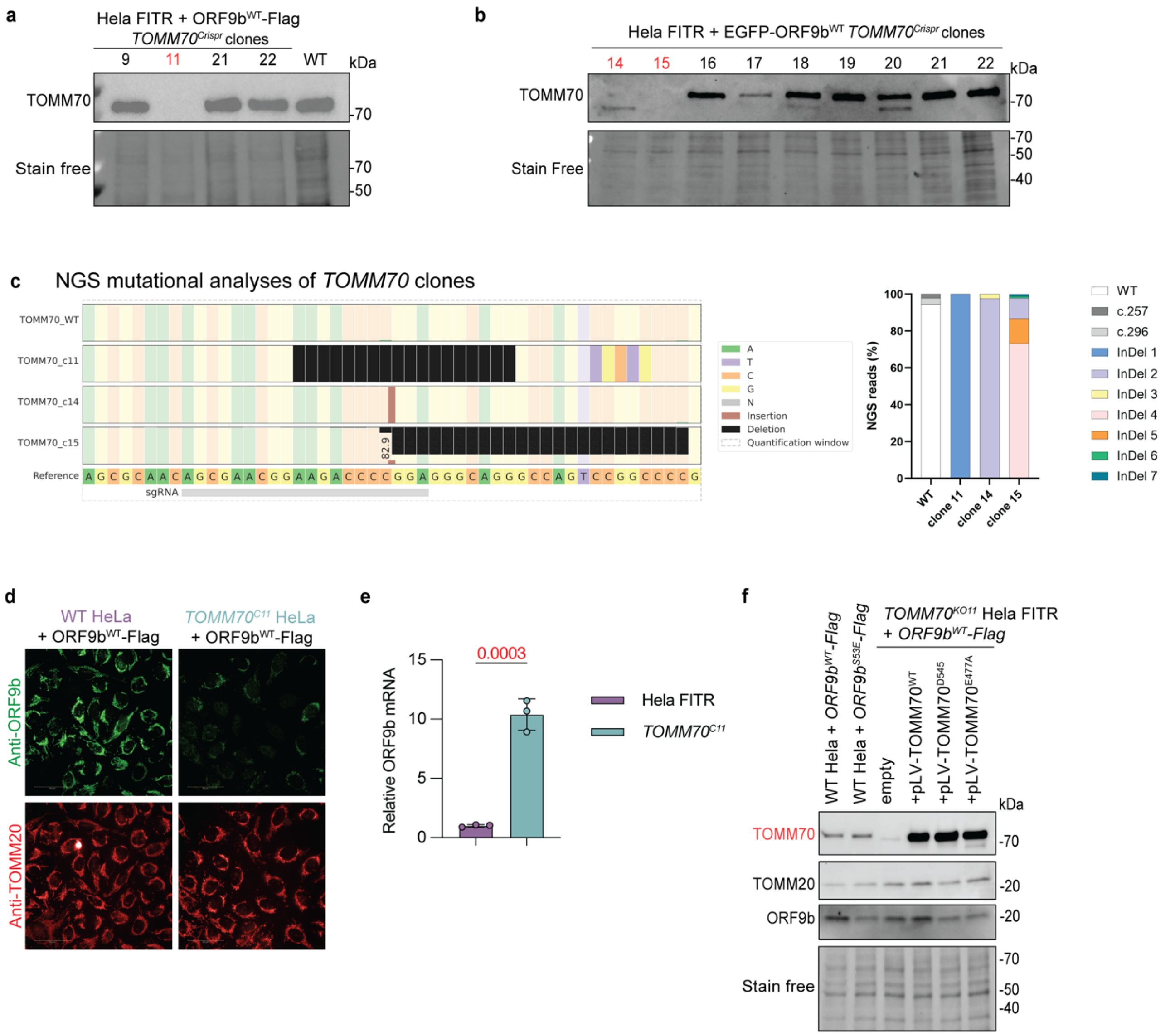

**Supplemental Figure 3: ORF9b recruitment requires the E477 residue on TOMM70**

a) Identification of *TOMM70* knockout clones generated by CRISPR/Cas9 genome editing of HeLa FITR ORFb9$^{WT}$-FLAG cells. Immunoblotting was performed with the indicated antibodies. Stain-free used as a loading control. Wild type (WT) HeLa FITR cells were used as a positive control.

b) Identification of *TOMM70* knockout clones generated by CRISPR/Cas9 genome editing of HeLa FITR EGFP-ORFb9$^{WT}$cells. Immunoblotting was performed with the indicated antibodies. Stain-free was used as a loading control.

c) (Left) NGS deep sequencing was performed on PCR products amplified from TOMM70-deficient clones *TOMM70*$^{C15}$, *TOMM70*$^{C14}$ capable of expressing EGFP-ORF9bWT and *TOMM70*$^{C11}$ capable of expressing ORF9bWT-FLAG, and parental controls (TOMM70_WT). Allelic distributions were analyzed using Crispresso2. sgRNA used to disrupt *TOMM70* is indicated in grey. (Right) percent of NGS reads for each cell line and is described in Supplemental Table 3.

d) Representative images of wild type HeLa FITR cells and *TOMM70*$^{C11}$ HeLa FITR cells induced to express ORFb9$^{WT}$-FLAG with anhydrotetracycline (AHT, 10ng/mL) for 24 hours. Immunofluorescence using anti-TOMM20 and anti-ORF9b antibodies. Scale bar=20 $\mu$m.

e) Measurement of ORF9b mRNA expression by RT-qPCR after 24h induction of EGFP-ORFb9$^{WT}$ in *TOMM70*$^{C15}$ HeLa FITR cells treated with AHT (10ng/mL) for 24 hours. Data are means ± SEM, 2-tailed unpaired Student's t test. Statistically significant exact P-value highlighted in red.

f) Western blots of lysates from *TOMM70*$^{C11}$ HeLa FITR cells in which wild type (pLV-TOMM70$^{WT}$) or mutant (pLV-TOMM70$^{E477A}$ or pLV-TOMM70$^{D545A}$) was stably expressed, and ORF9b$^{WT}$-FLAG expression was induced by anhydrotetracycline (AHT, 10ng/mL). Antibodies for ORF9b, TOMM20, TOMM70, and were used sequentially. Stain-free was used as a loading control.

Supplemental Figure 4

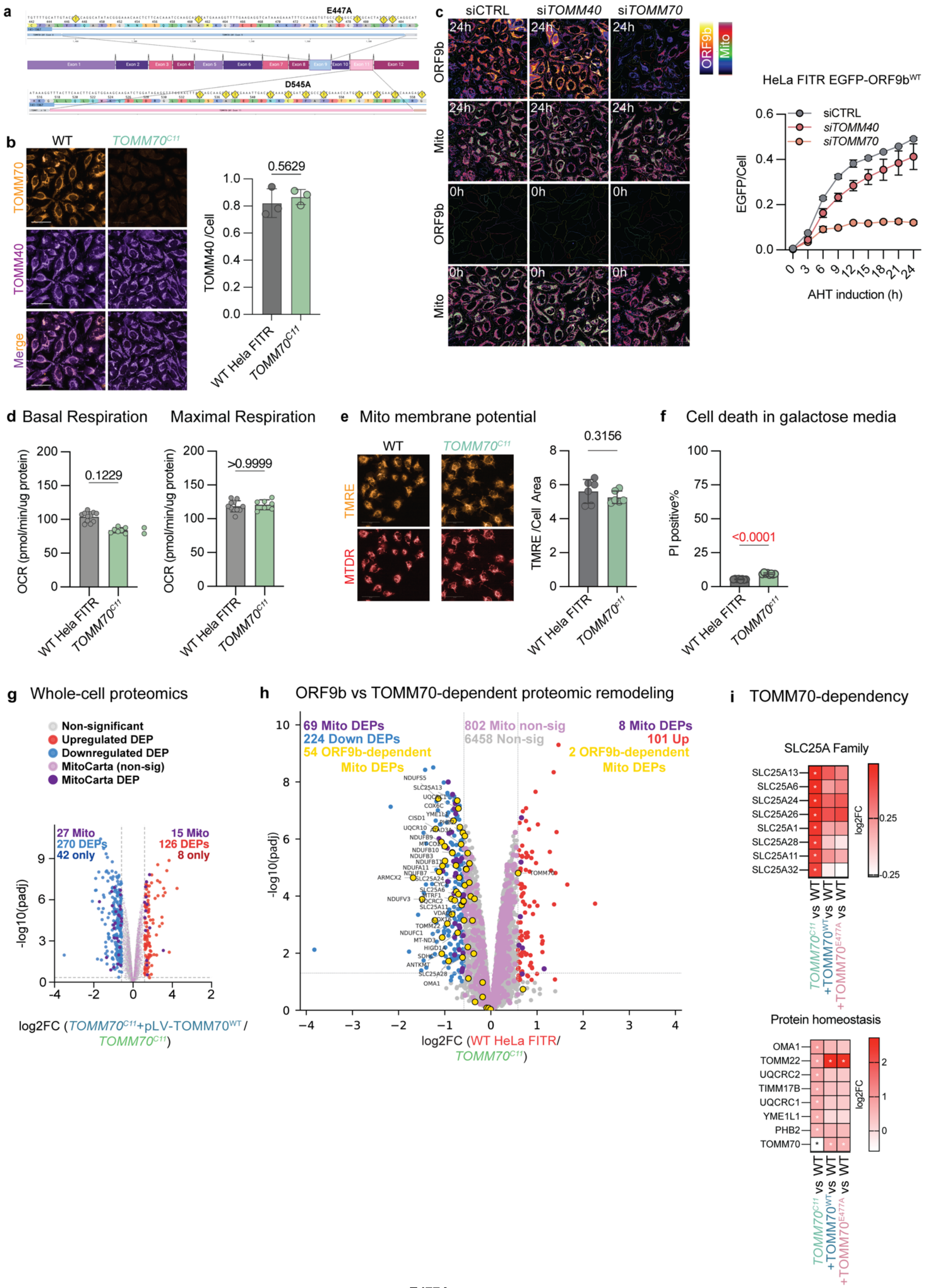


**Supplemental Figure 4: Mutant *TOMM70*$^{E477A}$ preserves canonical functions of TOMM70**

a) Gene structure of human *TOMM70*. Minor allele frequencies predicted to be deleterious according to Polyphen (>0.9) and SIFT (<0.1) are indicated by yellow hazard sign (!). E477A and D545A mutations in Exons 9 and 11, respectively.

b) Immunocytochemistry of WT and *TOMM70*$^{C11}$ HeLa FITR cells with TOMM40 and TOMM70 antibodies. Quantification of single-cell TOMM40 fluorescence signal intensity per cell (TOMM40/Cell). Data are means ± SEM, 2-tailed unpaired Welch's t test of 3 replicates with 430-677 cells were imaged for each replicate. Exact P-value indicated.
c) Representative images of live imaging of EGFP-ORF9b$^{WT}$ expressing cells subjected siRNA-mediated downregulation of *TOMM40* (si*TOMM40*) or *TOMM70* (si*TOMM70*) (left). Mitochondria were labeled with Mitotracker DeepRed (Mito) and Tetramethylrhodamine, Ethyl Ester (TMRE). Pseudo-colour indicates signal intensities for Mito and ORF9b. (Right) (Right) Quantification of single-cell fluorescence signal intensity of EGFP-ORF9bWT over time (EGFP/Cell). Data are means ± SEM.
d) Seahorse FluxAnalyzer analysis of mitochondrial respiration in wild type (WT) HeLa FITR cells, *TOMM70*$^{C11}$ HeLa FITR (*TOMM70*$^{C11}$ ) cultured in galactose-containing media for 5 days. Basal and maximal respiration rates were normalized by protein content. Data represent the mean ± SEM. Statistical significance was assessed using a two-tailed unpaired Student's t-test.
e) (Left) Representative images of live imaging of wild type (WT) HeLa FITR cells, *TOMM70*$^{C11}$ HeLa FITR (*TOMM70*$^{C11}$ ). Mitochondria were labeled with Mitotracker DeepRed (Mito) and Tetramethylrhodamine, Ethyl Ester (TMRE). (Right) Quantification of single-cell fluorescence signal intensity of mitochondrial membrane potential (TMRE/Cell). Data are means ± SEM, 2-tailed unpaired Welch's t test of 6 replicates with 143-623 cells were imaged for each replicate. Exact P-value indicated.
f) Cell death quantified after 2 days of culture in galactose-containing media by measuring. Cells were labeled with propidium iodide (PI) to mark dead cells NucBlue to mark all nuclei. Cell death is represented as a percentage of PI+ cells.
g) Volcano plot of whole-cell proteome of *TOMM70*$^{C11}$ HeLa FITR expressing TOMM70$^{WT}$ (*TOMM70*$^{C11}$ +pLV-TOMM70$^{WT}$, n=4) vs *TOMM70*$^{C11}$ HeLa FITR (*TOMM70*$^{C11}$, n=4) analyzed by mass spectrometry. (Purple) Mitochondrial proteins (MitoCarta 3.0), (light blue) differentially expressed proteins (DEPs) more abundant in *TOMM70*$^{C11}$ HeLa FITR, (dark blue), proteins exclusively measured in *TOMM70*$^{C11}$ HeLa FITR, (light red) DEPs more abundant in *TOMM70*$^{C11}$ +pLV-TOMM70$^{WT}$, (dark red), proteins exclusively measured in *TOMM70*$^{C11}$ +pLV-TOMM70$^{WT}$, (grey) non-significant proteins, (light purple) Non-significant MitoCarta 3.0 proteins. Differential analysis using LIMMA t-test is and corresponding p-values are adjusted using an adaptive Benjamini-Hochberg correction. Differentially expressed proteins (DEPs) are defined by an adjusted p-value is below 0.01 and a log2 fold-change greater than 1, delimited by dotted grey lines..
h) Volcano plot of whole-cell proteome of *TOMM70*$^{C11}$ HeLa FITR (*TOMM70*$^{C11}$, n=4) vs wild type (WT) HeLa FITR cells (WT, n=4) analyzed by mass spectrometry. (Purple) Mitochondrial proteins (MitoCarta 3.0), (light blue) differentially expressed proteins (DEPs) more abundant in *TOMM70*$^{C11}$ HeLa FITR, (light red) DEPs more abundant in WT HeLa FITR, (grey) non-significant proteins, (light purple) Non-significant MitoCarta 3.0 proteins, (yellow) ORF9b-dependent mitochondrial DEPs that are upregulated (54) or downregulated (2) in WT HeLa FITR cells expressing ORF9b$^{WT}$-FLAG (relative to WT HeLa FITR). Differential analysis using LIMMA t-test is and corresponding p-values are adjusted using an adaptive Benjamini-Hochberg correction. Differentially expressed proteins (DEPs) are defined by an adjusted p-value is below 0.01 and a log2 fold-change greater than 1, delimited by dotted grey lines.
i) Heatmaps of carrier proteins (SLC25A family), and protein homeostasis proteins that were quantified by whole-cell proteomics (see Supplemental Dataset 1) and significantly upregulated (red) in *TOMM70*$^{C11}$ HeLa FITR, *TOMM70*$^{C11}$ HeLa FITR expressing TOMM70$^{WT}$ or *TOMM70*$^{C11}$ HeLa FITR expressing TOMM70$^{E477A}$ relative to WT HeLa FITR. Significant differences are indicated with * defined by differential analysis using LIMMA t-test is and adaptive Benjamini-Hochberg correction.

**Supplemental Figure 5**

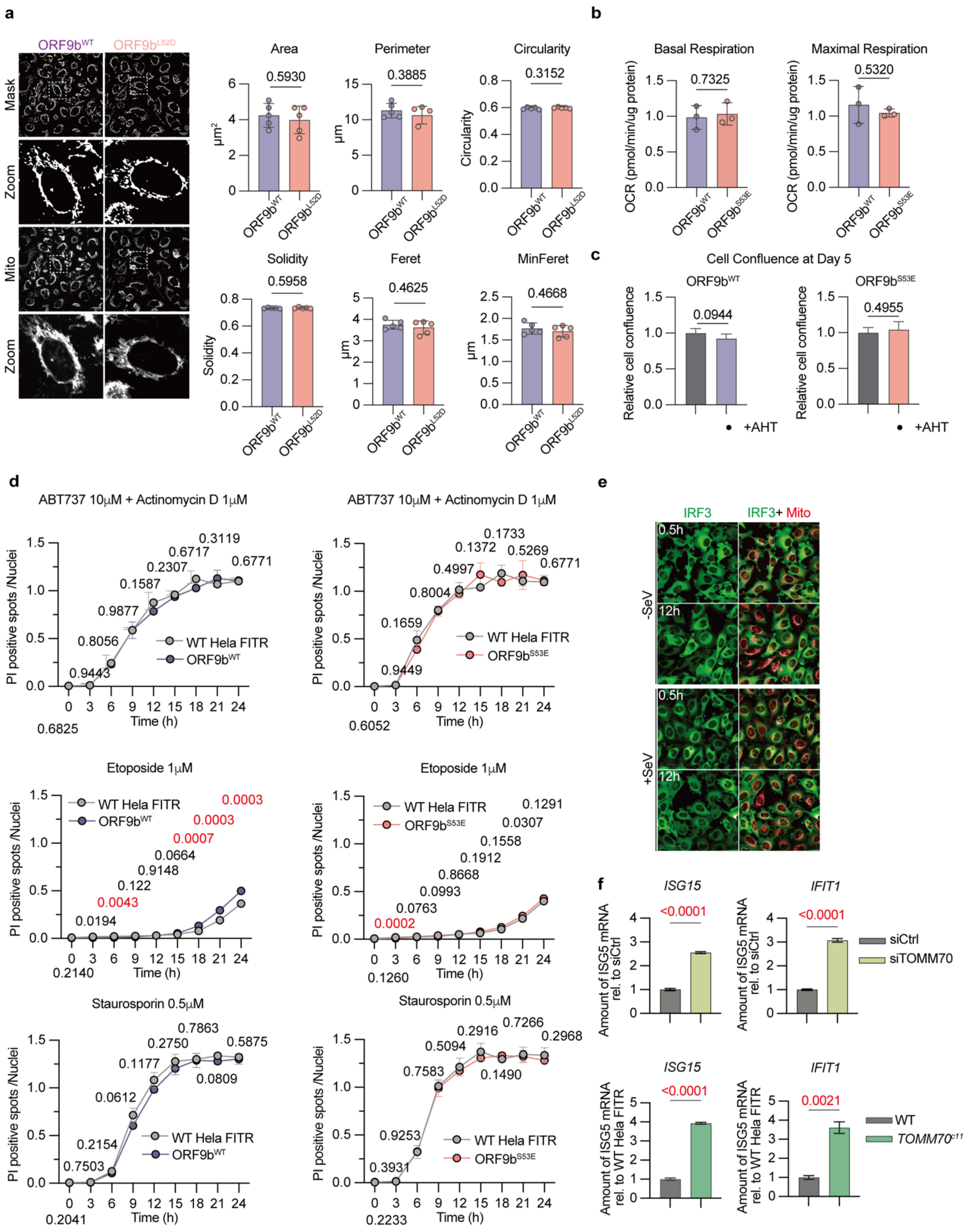

**Supplemental Figure 5: Characterization of ORF9b recruitment on signaling quality control and bioenergetics**

a) Representative images of control HeLa FITR cells induced with AHT to express EGFP-ORFb9$^{WT}$ (ORFb9$^{WT}$) or EGFP-ORFb9$^{L52D}$ (ORFb9$^{L52D}$). Time of induction 15 hours. Mitochondria were labeled with Mitotracker DeepRed (MTDR). Image analysis was performed with the open-source software ImageJ/Fiji using a custom macro for batch processing; masks were generated by automated thresholding, binarization, and morphological refinement. Quantified morphological parameters include area, perimeter, circularity, solidity, Feret diameter, and minimum Feret diameter. Data are means ± SEM, 2-tailed unpaired Welch's t test of 6 replicates with 240-483 cells were imaged for each replicate (3 replicates per group).

b) Seahorse FluxAnalyzer analysis of mitochondrial respiration of cells in grown in glucose-containing media. Basal and maximal respiration rates were measured in HeLa FITR cells treated with AHT to express ORF9b$^{WT}$ (blue, n=3) or ORF9b$^{S53E}$ (pink, n=3), normalized to cell content measured by Incucyte. Data are means ± SEM, 2-tailed unpaired Student's t test.

c) Cell proliferation in galactose-containing media of wild type (WT) HeLa FITR cells (grey, n=5), WT HeLa FITR cells expressing ORF9b$^{WT}$-FLAG (blue, n=5) and mutant ORF9b$^{S53E}$-FLAG (pink, n=5) were stably expressed. Bar graph represents cell confluence after 5 days of proliferation normalized to initial confluence. Data are means ± SEM, 2-tailed unpaired Student's t test. P-values are indicated.

d) Representative images of live imaging of cell death progression in HeLa FITR cells expressing ORF9b$^{WT}$-FLAG (left,) or ORF9b$^{S53E}$-FLAG (right). Induction of cell death with Actinomycin (ActD), ABT-737, Etoposide, Staurosporin for the indicated times and concentrations. Nuclei were labeled with NucBlue (NB) and dying cells were labeled with propidium iodide (PI). Cell death progression represented as the ratio of PI+ cells (PI/NB). Data are means ± SEM, 2-tailed unpaired Student's t test. P-values are indicated and significantly different P-values are highlighted in red.

e) Representative images of HeLa FITR cells stably expressing EGFP-IRF3 (IRF3, green) with mitochondria labeling with MitoTracker DeepRed (Mito, red) demonstrating the absence of mitochondrial localization upon infection with Sendai Virus (SeV) for the indicated times.

f) (Top) measurement *ISG15* and *IFIT1* by qRT-PCR in wild type HeLa FITR cells following 72h transfection with siCTRL (n=4) and siTOMM70 (n=4) siRNAs and measurement of *ISG15* and *IFIT1* by qRT-PCR in wild type (WT, n=4) and TOMM70-deficient ($TOMM70^{C11}$, n=4) HeLa FITR cells. Data are means ± SEM, 2-tailed unpaired Student's t test. P-values are indicated and significantly different P-values are highlighted in red.

### Supplemental Figure 6

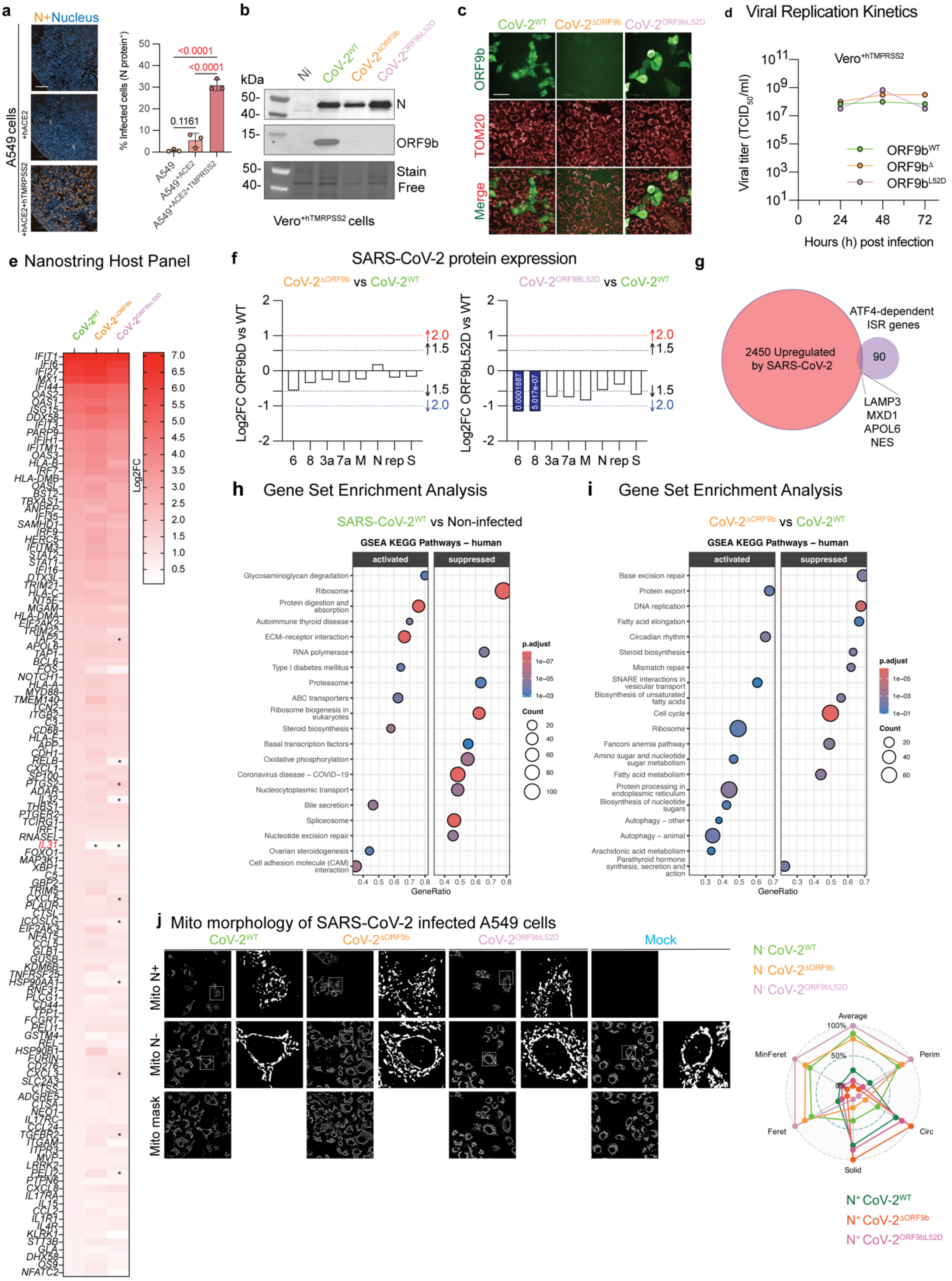


**Supplemental Figure 6: Characterization of ORF9-deficient SARS-CoV-2 in A549 cells.**

a) Immunofluorescence analysis of infection of A549 lung epithelial cells, and A549 cells stably expressing human ACE2, and A549 cells stably expressing hACE2 ($A549^{+hACE2}$) and human TMPRSS2 ($A549^{+hACE2+hTMPRSS2}$) 24h post-infection with wild-type Wuhan SARS-CoV-2 (CoV-$2^{WT}$at MOI 0.5. Anti-Nucleocapsid antibody (N) was used to label infected cells and DAPI was used to label all nuclei and imaged by high-content spinning disc microscopy. Single-cell quantification of N-positive cells (% infected cells) was determined with Harmony 5.1. Data are means ± SEM, of 24710-30796 cells measured per replicate analyzed by one-way ANOVA Tukey's multiple comparisons test. P-values are indicated and statistically significance highlighted in red.

b) Immunoblot analysis of whole-cell lysates from Vero cells stably expressing human TMPRSS2 ($Vero^{+hTMPRSS2}$) 24h post-infection with wild-type Wuhan SARS-CoV-2 ($CoV\text{-}2^{WT}$), SARS- $CoV\text{-}2^{\Delta ORF9b}$ ($CoV\text{-}2^{\Delta ORF9b}$), or $SARS\text{-}CoV\text{-}2^{ORF9bL52D}$ ($CoV\text{-}2^{ORF9bL52D}$) at MOI 0.5 and probed using the indicated antibodies.

c) Representative images of $A549^{+hACE2+hTMPRSS2}$ cells infected with Wuhan SARS-CoV-2 (SARS-$CoV\text{-}2^{WT}$), SARS- $CoV\text{-}2^{\Delta ORF9b}$, or $SARS\text{-}CoV\text{-}2^{Orf9bL52D}$ viruses at MOI 0.5 and fixed for indirect immunocytochemistry Indirect immunocytochemistry was performed with anti-TOMM20 to label mitochondria and anti-ORF9b to label ORF9b. Scale bar=50 µm.

d) Viral replication measured by TCID50 assay of $Vero^{+hTMPRSS2}$ cells infected with wild-type Wuhan SARS-CoV-2 ($CoV\text{-}2^{WT}$), SARS- $CoV\text{-}2^{\Delta ORF9b}$ ($CoV\text{-}2^{\Delta ORF9b}$), or $SARS\text{-}CoV\text{-}2^{ORF9bL52D}$ ($CoV\text{-}2^{ORF9bL52D}$) at MOI 0.01.

e) Host immune cell response measured by Nanostring mRNA analyses of A549 cells stably expressing human ACE2 and human TMPRSS2 ($A549^{+hACE2+hTMPRSS2}$) 24h post-infection with wild-type Wuhan SARS-CoV-2 ($CoV\text{-}2^{WT}$, n=3), SARS- $CoV\text{-}2^{\Delta ORF9b}$ ($CoV\text{-}2^{\Delta ORF9b}$, n=3), or $SARS\text{-}CoV\text{-}2^{ORF9bL52D}$ ($CoV\text{-}2^{ORF9bL52D}$, n=3). Heap map of significantly upregulated genes in $CoV\text{-}2^{WT}$ compared to non-infected mock control (n=3). Differentially expressed genes between $CoV\text{-}2^{ORF9bD}$ versus $CoV\text{-}2^{WT}$ and SARS-CoV- versus $CoV\text{-}2^{WT}$ highlighted in red and denoted with *.

f) Differential expression of SARS-CoV-2 viral proteins determined by proteomics. Pairwise comparisons between (left) Wuhan SARS-CoV-2 ($CoV\text{-}2^{WT}$) versus SARS- $CoV\text{-}2^{\Delta ORF9b}$ ($CoV\text{-}2^{\Delta ORF9b}$) and (right) $CoV\text{-}2^{WT}$ versus $SARS\text{-}CoV\text{-}2^{Orf9bL52D}$ ($CoV\text{-}2^{Orf9bL52D}$)-infected $A549^{+hACE2+hTMPRSS2}$ cells. Pvalues are indicated where significantly downregulated (blue) proteins were identified. Fold-change thresholds of 1.5 (black) and 2.0 (red or blue) are indicated by dotted lines. Data are available in Supplemental Dataset 6.

g) Venn Diagram of 2450 upregulated DEGs (Wuhan SARS-CoV-2 versus non-infected $A549^{+hACE2+hTMPRSS2}$ cells, red) and the 94 (purple) ATF4-dependent ISR genes (according to Labbé et al, 2024[43]) reveal an overlap of 4 genes.

h) Gene Set Enrichment Analyses (GSEA) representation of activated and suppressed KEGG pathways comparing bulk RNAseq transcriptomes of Wuhan SARS-CoV-2 ($SARS\text{-}CoV\text{-}2^{WT}$) versus non-infected mock control $A549^{+hACE2+hTMPRSS2}$ cells. Data are available in Supplemental Dataset 5.

i) Gene Set Enrichment Analyses (GSEA) representation of activated and suppressed KEGG pathways comparing bulk RNAseq transcriptomes of SARS- $CoV\text{-}2^{\Delta ORF9b}$ ($CoV\text{-}2^{\Delta ORF9b}$) versus Wuhan SARS-CoV-2 ($CoV\text{-}2^{WT}$) infected $A549^{+hACE2+hTMPRSS2}$ cells. Data are available in Supplemental Dataset 5.

j) Representative images of A549 cells infected with Wuhan SARS-CoV-2 ($CoV\text{-}2^{WT}$), SARS-$CoV\text{-}2^{\Delta ORF9b}$ ($CoV\text{-}2^{\Delta ORF9b}$), and $SARS\text{-}CoV\text{-}2^{Orf9bL52D}$ ($CoV\text{-}2^{Orf9bL52D}$). Mitochondria were labeled with an anti-TOMM40 antibody, and the viral nucleocapsid was detected using a SARS/SARS-CoV-2 nucleocapsid monoclonal antibody. Image analysis was performed with a custom ImageJ/Fiji macro for batch processing; masks were generated by automated thresholding, binarization, and morphological refinement. For mitochondrial morphology analysis, subtraction of viral nucleocapsid masks from mitochondrial masks was used to classify mitochondria as infected (nucleocapsid-positive/overlapping) or non-infected (nucleocapsid-negative/non-overlapping). Quantified morphological parameters include area, perimeter, circularity, solidity, Feret diameter, and minimum Feret diameter. Data are means ± SEM of 6 replicates with 63-512 infected cells and 399-1631 non-infected cells were imaged for each replicate.

**Supplemental Figure 7**

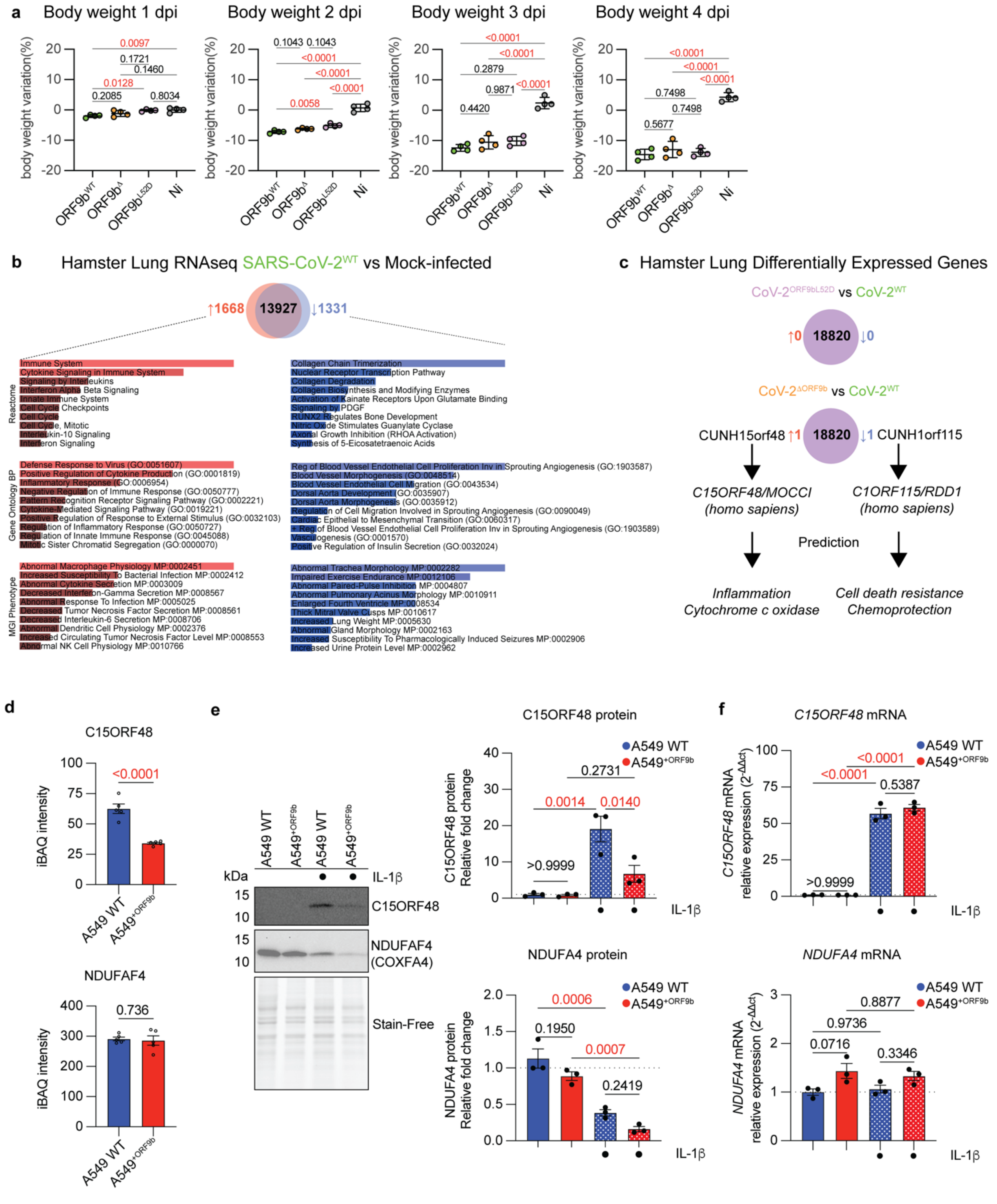


**Supplemental Figure 7: Characterization of ORF9-deficient SARS-CoV-2 in male Syrian hamsters.**

a) Daily body weight of 6-week old male Syrian hamsters at 4 days post-infection (dpi) with Wuhan SARS-CoV-2 (CoV-2$^{WT}$, n=4), SARS- CoV-2$^{\Delta ORF9b}$ (CoV-2$^{\Delta ORF9b}$, n=4) SARS-CoV-2$^{Orf9bL52D}$ (CoV-2$^{Orf9bL52D}$, n=4) viruses or non-infected Mock controls (n=4). Data are means ± SEM, one-way ANOVA Tukey's multiple comparisons test. P-values are indicated and statistically significance is highlighted in red.

b) RNAseq identification of differentially expressed genes (DEGs) in hamster lungs comparing infection with Wuhan SARS-CoV-2 (SARS-CoV-2$^{WT}$, n=4) versus non-infected mock control hamsters (n=4) 4 days post-infection (dpi). Venn diagram shows 13927 overlapping genes (purple), 1168 upregulated DEGs (red) and 1331 downregulated DEGs (blue) caused by infection with SARS-CoV-$2^{WT}$. Pathway analysis using Gene Ontology (GO), Reactome, and MGI Phenotype databases (see Supplemental Dataset 7).

c) Differential expressed genes (DEGs) defined by Log2FC>1 or <-1 between Wuhan SARS-CoV-2 (CoV-$2^{WT}$, n=4, green), SARS- CoV-$2^{\Delta ORF9b}$ (CoV-$2^{\Delta ORF9b}$, n=4, orange) SARS-CoV-$2^{Orf9bL52D}$ (CoV-$2^{Orf9bL52D}$, n=4, purple)

d) Differentially expression (iBAQ) of C15ORF48 in $A549^{+ORF9b}$ cells infected (n=5, red) versus parental wild type (WT, n=5, blue) control cells determined by whole-cell proteomics (see Supplemental Dataset 10). Differential analysis using LIMMA t-test is and corresponding p-values are adjusted using an adaptive Benjamini-Hochberg correction. P-values are indicated and statistically significance is highlighted in red.

e) Immunoblot analyses of wild type (WT, n=3) and ORF9b-expressing A549 cells ($A549^{+ORF9b}$, n=3) subjected to IL-1b (10ng/ml) stimulation for 24 hours using indicated antibodies. Relative intensity was normalized to stain-free. Data are means ± SEM, analyzed by one-way ANOVA Tukey's multiple comparisons test. P-values are indicated and statistically significance highlighted in red.

f) Quantification of C15ORF48 and NDUFA4 mRNA levels by wild type (WT, n=3) and ORF9b-expressing A549 cells ($A549^{+ORF9b}$, n=3) subjected to IL-1b (10ng/ml) stimulation for 24 hours and normalized to beta-acting. Data are means ± SEM, analyzed by one-way ANOVA Tukey's multiple comparisons test. P-values are indicated and statistically significance highlighted in red.

*h – Tukey's multiple comparisons test (two-way ANOVA).*
*Number of families: 17 | Number of comparisons per family: 6 | Alpha: 0.05*

| Time (h) | Comparison | Mean diff. | 95% CI of diff. | Sig. | Adj. P |
|---|---|---|---|---|---|
| 0 | SARS-CoV-2 vs. BANAL-236 | 0.00 | -21.24 to 21.24 | ns | >0.9999 |
| 0 | SARS-CoV-2 vs. RaTG13 | 0.00 | -21.24 to 21.24 | ns | >0.9999 |
| 0 | SARS-CoV-2 vs. SARS-CoV-1 | 0.00 | -21.24 to 21.24 | ns | >0.9999 |
| 0 | BANAL-236 vs. RaTG13 | 0.00 | -21.24 to 21.24 | ns | >0.9999 |
| 0 | BANAL-236 vs. SARS-CoV-1 | 0.00 | -21.24 to 21.24 | ns | >0.9999 |
| 0 | RaTG13 vs. SARS-CoV-1 | 0.00 | -21.24 to 21.24 | ns | >0.9999 |
| 1.5 | SARS-CoV-2 vs. BANAL-236 | 2.24 | -19.00 to 23.48 | ns | 0.9929 |
| 1.5 | SARS-CoV-2 vs. RaTG13 | -0.88 | -22.12 to 20.36 | ns | 0.9996 |
| 1.5 | SARS-CoV-2 vs. SARS-CoV-1 | 11.95 | -9.291 to 33.18 | ns | 0.4682 |
| 1.5 | BANAL-236 vs. RaTG13 | -3.12 | -24.36 to 18.12 | ns | 0.9814 |
| 1.5 | BANAL-236 vs. SARS-CoV-1 | 9.71 | -11.53 to 30.94 | ns | 0.6405 |
| 1.5 | RaTG13 vs. SARS-CoV-1 | 12.83 | -8.412 to 34.06 | ns | 0.4040 |
| 3 | SARS-CoV-2 vs. BANAL-236 | 2.90 | -18.34 to 24.13 | ns | 0.9851 |
| 3 | SARS-CoV-2 vs. RaTG13 | -4.96 | -26.20 to 16.28 | ns | 0.9313 |
| 3 | SARS-CoV-2 vs. SARS-CoV-1 | 18.95 | -2.287 to 40.19 | ns | 0.0993 |
| 3 | BANAL-236 vs. RaTG13 | -7.85 | -29.09 to 13.38 | ns | 0.7755 |
| 3 | BANAL-236 vs. SARS-CoV-1 | 16.05 | -5.182 to 37.29 | ns | 0.2089 |
| 3 | RaTG13 vs. SARS-CoV-1 | 23.91 | 2.672 to 45.15 | * | 0.0202 |
| 4.5 | SARS-CoV-2 vs. BANAL-236 | -5.81 | -27.04 to 15.43 | ns | 0.8949 |
| 4.5 | SARS-CoV-2 vs. RaTG13 | -7.90 | -29.13 to 13.34 | ns | 0.7727 |
| 4.5 | SARS-CoV-2 vs. SARS-CoV-1 | 19.66 | -1.576 to 40.90 | ns | 0.0809 |
| 4.5 | BANAL-236 vs. RaTG13 | -2.09 | -23.33 to 19.15 | ns | 0.9943 |
| 4.5 | BANAL-236 vs. SARS-CoV-1 | 25.47 | 4.231 to 46.70 | * | 0.0113 |
| 4.5 | RaTG13 vs. SARS-CoV-1 | 27.56 | 6.320 to 48.79 | ** | 0.0049 |
| 6 | SARS-CoV-2 vs. BANAL-236 | -6.40 | -27.64 to 14.84 | ns | 0.8647 |
| 6 | SARS-CoV-2 vs. RaTG13 | -17.96 | -39.20 to 3.278 | ns | 0.1301 |
| 6 | SARS-CoV-2 vs. SARS-CoV-1 | 9.68 | -11.56 to 30.92 | ns | 0.6426 |
| 6 | BANAL-236 vs. RaTG13 | -11.56 | -32.80 to 9.678 | ns | 0.4975 |
| 6 | BANAL-236 vs. SARS-CoV-1 | 16.08 | -5.159 to 37.32 | ns | 0.2078 |
| 6 | RaTG13 vs. SARS-CoV-1 | 27.64 | 6.400 to 48.87 | ** | 0.0048 |
| 7.5 | SARS-CoV-2 vs. BANAL-236 | -10.41 | -31.65 to 10.83 | ns | 0.5861 |
| 7.5 | SARS-CoV-2 vs. RaTG13 | -7.23 | -28.46 to 14.01 | ns | 0.8164 |
| 7.5 | SARS-CoV-2 vs. SARS-CoV-1 | 11.23 | -10.01 to 32.47 | ns | 0.5227 |
| 7.5 | BANAL-236 vs. RaTG13 | 3.18 | -18.05 to 24.42 | ns | 0.9803 |
| 7.5 | BANAL-236 vs. SARS-CoV-1 | 21.64 | 0.4017 to 42.88 | * | 0.0439 |
| 7.5 | RaTG13 vs. SARS-CoV-1 | 18.46 | -2.782 to 39.69 | ns | 0.1139 |
| 9 | SARS-CoV-2 vs. BANAL-236 | -14.80 | -36.03 to 6.440 | ns | 0.2761 |
| 9 | SARS-CoV-2 vs. RaTG13 | -12.43 | -33.66 to 8.809 | ns | 0.4326 |
| 9 | SARS-CoV-2 vs. SARS-CoV-1 | 3.30 | -17.94 to 24.54 | ns | 0.9782 |
| 9 | BANAL-236 vs. RaTG13 | 2.37 | -18.87 to 23.61 | ns | 0.9917 |
| 9 | BANAL-236 vs. SARS-CoV-1 | 18.10 | -3.142 to 39.33 | ns | 0.1255 |
| 9 | RaTG13 vs. SARS-CoV-1 | 15.73 | -5.511 to 36.96 | ns | 0.2253 |
| 10.5 | SARS-CoV-2 vs. BANAL-236 | -16.67 | -37.90 to 4.571 | ns | 0.1807 |
| 10.5 | SARS-CoV-2 vs. RaTG13 | -16.09 | -37.32 to 5.149 | ns | 0.2073 |
| 10.5 | SARS-CoV-2 vs. SARS-CoV-1 | 3.29 | -17.94 to 24.53 | ns | 0.9783 |
| 10.5 | BANAL-236 vs. RaTG13 | 0.58 | -20.66 to 21.81 | ns | 0.9999 |
| 10.5 | BANAL-236 vs. SARS-CoV-1 | 19.96 | -1.276 to 41.20 | ns | 0.0741 |
| 10.5 | RaTG13 vs. SARS-CoV-1 | 19.38 | -1.854 to 40.62 | ns | 0.0878 |
| 12 | SARS-CoV-2 vs. BANAL-236 | -11.78 | -33.02 to 9.455 | ns | 0.4806 |
| 12 | SARS-CoV-2 vs. RaTG13 | -3.52 | -24.76 to 17.72 | ns | 0.9738 |
| 12 | SARS-CoV-2 vs. SARS-CoV-1 | 2.23 | -19.00 to 23.47 | ns | 0.9930 |
| 12 | BANAL-236 vs. RaTG13 | 8.26 | -12.97 to 29.50 | ns | 0.7473 |
| 12 | BANAL-236 vs. SARS-CoV-1 | 14.01 | -7.222 to 35.25 | ns | 0.3237 |
| 12 | RaTG13 vs. SARS-CoV-1 | 5.75 | -15.48 to 26.99 | ns | 0.8976 |
| 13.5 | SARS-CoV-2 vs. BANAL-236 | -14.44 | -35.67 to 6.800 | ns | 0.2974 |
| 13.5 | SARS-CoV-2 vs. RaTG13 | -9.12 | -30.36 to 12.12 | ns | 0.6849 |
| 13.5 | SARS-CoV-2 vs. SARS-CoV-1 | 7.36 | -13.88 to 28.59 | ns | 0.8081 |
| 13.5 | BANAL-236 vs. RaTG13 | 5.32 | -15.92 to 26.55 | ns | 0.9170 |
| 13.5 | BANAL-236 vs. SARS-CoV-1 | 21.79 | 0.5582 to 43.03 | * | 0.0418 |
| 13.5 | RaTG13 vs. SARS-CoV-1 | 16.48 | -4.758 to 37.72 | ns | 0.1890 |
| 15 | SARS-CoV-2 vs. BANAL-236 | -9.44 | -30.68 to 11.80 | ns | 0.6607 |
| 15 | SARS-CoV-2 vs. RaTG13 | -1.35 | -22.58 to 19.89 | ns | 0.9984 |
| 15 | SARS-CoV-2 vs. SARS-CoV-1 | 8.31 | -12.93 to 29.55 | ns | 0.7440 |
| 15 | BANAL-236 vs. RaTG13 | 8.10 | -13.14 to 29.33 | ns | 0.7591 |
| 15 | BANAL-236 vs. SARS-CoV-1 | 17.75 | -3.485 to 38.99 | ns | 0.1374 |
| 15 | RaTG13 vs. SARS-CoV-1 | 9.66 | -11.58 to 30.89 | ns | 0.6442 |
| 16.5 | SARS-CoV-2 vs. BANAL-236 | -9.78 | -31.02 to 11.46 | ns | 0.6349 |
| 16.5 | SARS-CoV-2 vs. RaTG13 | -12.44 | -33.68 to 8.792 | ns | 0.4314 |
| 16.5 | SARS-CoV-2 vs. SARS-CoV-1 | 1.25 | -19.99 to 22.48 | ns | 0.9988 |
| 16.5 | BANAL-236 vs. RaTG13 | -2.67 | -23.90 to 18.57 | ns | 0.9883 |
| 16.5 | BANAL-236 vs. SARS-CoV-1 | 11.02 | -10.21 to 32.26 | ns | 0.5385 |
| 16.5 | RaTG13 vs. SARS-CoV-1 | 13.69 | -7.547 to 34.93 | ns | 0.3448 |
| 18 | SARS-CoV-2 vs. BANAL-236 | -5.62 | -26.85 to 15.62 | ns | 0.9038 |
| 18 | SARS-CoV-2 vs. RaTG13 | 0.23 | -21.01 to 21.46 | ns | >0.9999 |
| 18 | SARS-CoV-2 vs. SARS-CoV-1 | 2.00 | -19.23 to 23.24 | ns | 0.9949 |
| 18 | BANAL-236 vs. RaTG13 | 5.84 | -15.39 to 27.08 | ns | 0.8932 |
| 18 | BANAL-236 vs. SARS-CoV-1 | 7.62 | -13.62 to 28.86 | ns | 0.7911 |
| 18 | RaTG13 vs. SARS-CoV-1 | 1.78 | -19.46 to 23.01 | ns | 0.9964 |
| 19.5 | SARS-CoV-2 vs. BANAL-236 | -9.92 | -31.16 to 11.32 | ns | 0.6240 |
| 19.5 | SARS-CoV-2 vs. RaTG13 | -4.59 | -25.83 to 16.65 | ns | 0.9445 |
| 19.5 | SARS-CoV-2 vs. SARS-CoV-1 | 2.36 | -18.88 to 23.60 | ns | 0.9918 |
| 19.5 | BANAL-236 vs. RaTG13 | 5.33 | -15.91 to 26.57 | ns | 0.9164 |
| 19.5 | BANAL-236 vs. SARS-CoV-1 | 12.28 | -8.956 to 33.52 | ns | 0.4434 |
| 19.5 | RaTG13 vs. SARS-CoV-1 | 6.95 | -14.29 to 28.19 | ns | 0.8332 |
| 21 | SARS-CoV-2 vs. BANAL-236 | 1.72 | -19.51 to 22.96 | ns | 0.9968 |
| 21 | SARS-CoV-2 vs. RaTG13 | -0.56 | -21.79 to 20.68 | ns | 0.9999 |
| 21 | SARS-CoV-2 vs. SARS-CoV-1 | -1.13 | -22.36 to 20.11 | ns | 0.9991 |
| 21 | BANAL-236 vs. RaTG13 | -2.28 | -23.51 to 18.96 | ns | 0.9926 |
| 21 | BANAL-236 vs. SARS-CoV-1 | -2.85 | -24.08 to 18.39 | ns | 0.9858 |
| 21 | RaTG13 vs. SARS-CoV-1 | -0.57 | -21.81 to 20.67 | ns | 0.9999 |
| 22.5 | SARS-CoV-2 vs. BANAL-236 | -6.52 | -27.76 to 14.72 | ns | 0.8580 |
| 22.5 | SARS-CoV-2 vs. RaTG13 | 6.61 | -14.63 to 27.84 | ns | 0.8532 |
| 22.5 | SARS-CoV-2 vs. SARS-CoV-1 | 0.05 | -21.19 to 21.29 | ns | >0.9999 |
| 22.5 | BANAL-236 vs. RaTG13 | 13.13 | -8.109 to 34.36 | ns | 0.3828 |
| 22.5 | BANAL-236 vs. SARS-CoV-1 | 6.57 | -14.67 to 27.81 | ns | 0.8553 |
| 22.5 | RaTG13 vs. SARS-CoV-1 | -6.56 | -27.79 to 14.68 | ns | 0.8560 |
| 24 | SARS-CoV-2 vs. BANAL-236 | 0.00 | -21.24 to 21.24 | ns | >0.9999 |
| 24 | SARS-CoV-2 vs. RaTG13 | 0.00 | -21.24 to 21.24 | ns | >0.9999 |
| 24 | SARS-CoV-2 vs. SARS-CoV-1 | 0.00 | -21.24 to 21.24 | ns | >0.9999 |
| 24 | BANAL-236 vs. RaTG13 | 0.00 | -21.24 to 21.24 | ns | >0.9999 |
| 24 | BANAL-236 vs. SARS-CoV-1 | 0.00 | -21.24 to 21.24 | ns | >0.9999 |
| 24 | RaTG13 vs. SARS-CoV-1 | 0.00 | -21.24 to 21.24 | ns | >0.9999 |

**Supplemental Table 2.** *Washout AHT 24h – Tukey's multiple comparisons test (two-way ANOVA).*
*Within each row, compare columns (simple effects within rows). Number of families: 9 | Number of comparisons per family: 6 | Alpha: 0.05*

| Time (h) | Comparison | Mean diff. | 95% CI of diff. | Sig. | Adj. P |
|---|---|---|---|---|---|
| 0 | SARS-CoV-2 vs. BANAL-236 | 0.00 | -49.94 to 49.94 | ns | >0.9999 |
| 0 | SARS-CoV-2 vs. RaTG13 | -9.94 | -59.79 to 39.91 | ns | 0.9326 |
| 0 | SARS-CoV-2 vs. SARS-CoV-1 | 0.00 | -44.24 to 44.24 | ns | >0.9999 |
| 0 | BANAL-236 vs. RaTG13 | -9.94 | -56.74 to 36.87 | ns | 0.9198 |
| 0 | BANAL-236 vs. SARS-CoV-1 | 0.00 | -40.12 to 40.12 | ns | >0.9999 |
| 0 | RaTG13 vs. SARS-CoV-1 | 9.94 | -30.60 to 50.48 | ns | 0.8689 |
| 3 | SARS-CoV-2 vs. BANAL-236 | 29.30 | -3.616 to 62.21 | ns | 0.0833 |
| 3 | SARS-CoV-2 vs. RaTG13 | 19.07 | -14.63 to 52.76 | ns | 0.3522 |
| 3 | SARS-CoV-2 vs. SARS-CoV-1 | 2.26 | -43.41 to 47.94 | ns | 0.9988 |
| 3 | BANAL-236 vs. RaTG13 | -10.23 | -31.63 to 11.17 | ns | 0.4997 |
| 3 | BANAL-236 vs. SARS-CoV-1 | -27.04 | -68.04 to 13.97 | ns | 0.2361 |
| 3 | RaTG13 vs. SARS-CoV-1 | -16.80 | -58.29 to 24.69 | ns | 0.6122 |
| 6 | SARS-CoV-2 vs. BANAL-236 | 0.47 | -43.93 to 44.87 | ns | >0.9999 |
| 6 | SARS-CoV-2 vs. RaTG13 | -4.89 | -61.02 to 51.24 | ns | 0.9938 |
| 6 | SARS-CoV-2 vs. SARS-CoV-1 | -20.69 | -65.99 to 24.60 | ns | 0.5339 |
| 6 | BANAL-236 vs. RaTG13 | -5.36 | -52.84 to 42.12 | ns | 0.9823 |
| 6 | BANAL-236 vs. SARS-CoV-1 | -21.16 | -46.68 to 4.365 | ns | 0.1196 |
| 6 | RaTG13 vs. SARS-CoV-1 | -15.80 | -63.87 to 32.27 | ns | 0.7382 |
| 9 | SARS-CoV-2 vs. BANAL-236 | 9.96 | -32.66 to 52.58 | ns | 0.8879 |
| 9 | SARS-CoV-2 vs. RaTG13 | 11.44 | -30.42 to 53.29 | ns | 0.8173 |
| 9 | SARS-CoV-2 vs. SARS-CoV-1 | -7.13 | -52.05 to 37.78 | ns | 0.9628 |
| 9 | BANAL-236 vs. RaTG13 | 1.48 | -18.50 to 21.46 | ns | 0.9959 |
| 9 | BANAL-236 vs. SARS-CoV-1 | -17.09 | -47.63 to 13.44 | ns | 0.3630 |
| 9 | RaTG13 vs. SARS-CoV-1 | -18.57 | -47.78 to 10.63 | ns | 0.2359 |
| 12 | SARS-CoV-2 vs. BANAL-236 | -0.44 | -24.89 to 24.00 | ns | >0.9999 |
| 12 | SARS-CoV-2 vs. RaTG13 | 13.22 | -11.18 to 37.62 | ns | 0.4206 |
| 12 | SARS-CoV-2 vs. SARS-CoV-1 | -17.52 | -46.42 to 11.37 | ns | 0.3135 |
| 12 | BANAL-236 vs. RaTG13 | 13.67 | -8.546 to 35.88 | ns | 0.3189 |
| 12 | BANAL-236 vs. SARS-CoV-1 | -17.08 | -44.59 to 10.43 | ns | 0.2876 |
| 12 | RaTG13 vs. SARS-CoV-1 | -30.75 | -58.23 to -3.269 | * | 0.0280 |
| 15 | SARS-CoV-2 vs. BANAL-236 | -18.40 | -38.27 to 1.468 | ns | 0.0723 |
| 15 | SARS-CoV-2 vs. RaTG13 | 1.22 | -17.32 to 19.76 | ns | 0.9973 |
| 15 | SARS-CoV-2 vs. SARS-CoV-1 | -24.92 | -62.11 to 12.26 | ns | 0.2141 |
| 15 | BANAL-236 vs. RaTG13 | 19.62 | -1.594 to 40.84 | ns | 0.0736 |
| 15 | BANAL-236 vs. SARS-CoV-1 | -6.52 | -44.19 to 31.15 | ns | 0.9459 |
| 15 | RaTG13 vs. SARS-CoV-1 | -26.15 | -63.57 to 11.28 | ns | 0.1962 |
| 18 | SARS-CoV-2 vs. BANAL-236 | -9.07 | -30.27 to 12.12 | ns | 0.5886 |
| 18 | SARS-CoV-2 vs. RaTG13 | 12.58 | -9.594 to 34.76 | ns | 0.3734 |
| 18 | SARS-CoV-2 vs. SARS-CoV-1 | -10.75 | -38.80 to 17.30 | ns | 0.6787 |
| 18 | BANAL-236 vs. RaTG13 | 21.66 | 3.467 to 39.85 | * | 0.0179 |
| 18 | BANAL-236 vs. SARS-CoV-1 | -1.68 | -27.59 to 24.24 | ns | 0.9972 |
| 18 | RaTG13 vs. SARS-CoV-1 | -23.33 | -49.91 to 3.248 | ns | 0.0924 |
| 21 | SARS-CoV-2 vs. BANAL-236 | -1.35 | -17.25 to 14.55 | ns | 0.9945 |
| 21 | SARS-CoV-2 vs. RaTG13 | 22.38 | 5.932 to 38.83 | ** | 0.0070 |
| 21 | SARS-CoV-2 vs. SARS-CoV-1 | -5.71 | -28.52 to 17.11 | ns | 0.8708 |
| 21 | BANAL-236 vs. RaTG13 | 23.73 | 8.147 to 39.31 | ** | 0.0029 |
| 21 | BANAL-236 vs. SARS-CoV-1 | -4.36 | -26.79 to 18.07 | ns | 0.9305 |
| 21 | RaTG13 vs. SARS-CoV-1 | -28.09 | -50.76 to -5.414 | * | 0.0155 |
| 24 | SARS-CoV-2 vs. BANAL-236 | -6.26 | -24.34 to 11.81 | ns | 0.7456 |
| 24 | SARS-CoV-2 vs. RaTG13 | 18.41 | 4.242 to 32.59 | * | 0.0104 |
| 24 | SARS-CoV-2 vs. SARS-CoV-1 | -11.05 | -36.27 to 14.16 | ns | 0.5678 |
| 24 | BANAL-236 vs. RaTG13 | 24.68 | 8.182 to 41.17 | ** | 0.0042 |
| 24 | BANAL-236 vs. SARS-CoV-1 | -4.79 | -30.80 to 21.22 | ns | 0.9456 |
| 24 | RaTG13 vs. SARS-CoV-1 | -29.47 | -54.01 to -4.930 | * | 0.0197 |

**Supplemental Table 3** – NGS characterization of TOMM70-deficient HeLa FITR cell lines

| Name | cDNA sequence change | Amino Acid change | Reference (Transcript ID) | Clones |
|---|---|---|---|---|
| c.257 | c.257C>A | p.P86Q | ENST00000284320 | WT |
| c.296 | c.296C>A | p.P94Q | ENST00000284320 | WT |
| InDel 1 | c.A250_G267del, c.274C>T, c.275C>G, c.276G>C, c.277G>T, c.278C>G | p.K84_R89del p.P92C p.A93C | ENST00000284320 | 11 |
| InDel 2 | c.257_258insC | p.P86fsX14 | ENST00000284320 | 14, 15 |
| InDel 3 | c.257_258insC, c.281C>A | p.P86fsX14 | ENST00000284320 | 14 |
| InDel 4 | c.G258_C281del | p.E87_P94del | ENST00000284320 | 15 |
| InDel 5 | c.C257del | p.P86RfsX22 | ENST00000284320 | 15 |
| InDel 6 | c.257C>A, c.G258_C281del | p.P86HfsX6 | ENST00000284320 | 15 |
| InDel 7 | c.A178_G302del | p.Q62_G101del p.P102CfsX8 | ENST00000284320 | 15 |

**Supplemental Table 4** NGS determination of SARS-CoV-2 viral sequences

SARS-CoV-2

| | Protein | Nucleotide | Amino Acid | Frequency in NGS reads (%) |
|---|---|---|---|---|
| A | ORF1 a-b (NSP3) | C6666A | P2134H | 14.1 |
| | ORF1 a-b (NSP3) | C8043T | A2593V | 10.4 |
| | N | C29518T | D415D | 12.1 |
| B | ORF1 a-b (NSP3) | C6666A | P2134H | 12.5 |
| C | ORF1 a-b (NSP3) | C6666A | P2134H | 11.6 |
| | ORF1 a-b (NSP3) | C8043T | A2593V | 11.6 |
| | N | C29518T | D415D | 14.7 |
| D | ORF1 a-b (NSP3) | C6666A | P2134H | 15.7 |
| | ORF1 a-b (NSP3) | C8043T | A2593V | 11.1 |
| | S | C24768A | P1069H | 10.1 |
| | N | C28892A | P207T | 10.2 |
| | N | C29518T | D415D | 12.7 |

SARS-CoV-2$^{\Delta ORF9b}$

| | Protein | Nucleotide | Amino Acid | Frequency in NGS reads (%) |
|---|---|---|---|---|
| A | ORF1 a-b (NSP3) | C6666A | P2134H | 11.5 |
| | N | C29518T | D415D | 15.7 |
| B | ORF1 a-b (NSP3) | C6666A | P2134H | 14.0 |
| | N | C29518T | D415D | 12.8 |
| C | ORF1 a-b (NSP3) | C6666A | P2134H | 15.3 |
| | N | C29518T | D415D | 10.0 |
| D | ORF1 a-b (NSP3) | C6666A | P2134H | 13.7 |
| | N | C29518T | D415D | 11.5 |

SARS-CoV-2$^{ORF9bL52D}$

| | Protein | Nucleotide | Amino Acid | Frequency in NGS reads (%) |
|---|---|---|---|---|
| A | ORF1 a-b (NSP3) | C6666A | P2134H | 12.5 |
| | S | C24768A | P1069H | 10.0 |
| | N | C29518T | D415D | 16.0 |
| B | ORF1 a-b (NSP3) | C6666A | P2134H | 19.1 |
| | N | C29518T | D415D | 14.8 |
| C | ORF1 a-b (NSP3) | C6666A | P2134H | 14.3 |
| | S | C24768A | P1069H | 10.2 |
| | N | C29518T | D415D | 12.8 |
| D | ORF1 a-b (NSP3) | C6666A | P2134H | 15.3 |
| | S | C24768A | P1069H | 10.8 |
| | N | C29518T | D415D | 10.8 |

**Supplemental Table 5** NGS determination of SARS-CoV-2 viral sequences in Hamster lungs

Hamsters

SARS-CoV-2

| | Protein | Nucleotide | Amino Acid | Frequency in NGS reads (%) |
|---|---|---|---|---|
| M1 | ORF1 a-b (NSP3) | C8354T | R3697C | 12.3 |
| | N | C29518T | D415D | 17.5 |
| M2 | ORF1 a-b (NSP3) | C8354T | R3697C | 20.2 |
| | ORF1 a-b (NSP10) | T13123C | S4286S | 13.6 |
| | N | C29518T | D415D | 14.8 |
| M3 | ORF1 a-b (NSP3) | C8354T | R3697C | 12.4 |
| | N | C29518T | D415D | 19.1 |
| M4 | N | C29518T | D415D | 18.9 |

SARS-CoV-2$^{\Delta ORF9b}$

| | Protein | Nucleotide | Amino Acid | Frequency in NGS reads (%) |
|---|---|---|---|---|
| M1 | N | C29518T | D415D | 16.6 |
| M2 | N | C29518T | D415D | 19.9 |
| M3 | N | C29518T | D415D | 23.7 |
| M4 | ORF1 a-b (NSP10) | T13123C | S4286S | 10.0 |
| | N | C29518T | D415D | 22.4 |

SARS-CoV-2$^{ORF9bL52D}$

| | Protein | Nucleotide | Amino Acid | Frequency in NGS reads (%) |
|---|---|---|---|---|
| M1 | ORF1 a-b (NSP10) | T13123C | S4286S | 10.0 |
| | N | C29518T | D415D | 19.3 |
| M2 | ORF1 a-b (NSP2) | T1614A | L450H | 10.3 |
| | N | C29518T | D415D | 22.8 |
| M3 | ORF1 a-b (NSP3) | C8354T | R3697C | 12.8 |
| M4 | ORF1 a-b (NSP2) | T1614A | L450H | 11.2 |
| | N | C29518T | D415D | 26.5 |

**Reagents and Tools Table**

| Reagent/Resource | Reference or Source | Identifier or Catalog Number |
|---|---|---|
| **Experimental Models** | | |
| HEK293T (293T) cells | ATCC | CRL-3216 |
| Vero-E6 cells | ATCC | CRL-1586 |
| HeLa Flp-In™ T-REx™ (HeLa FITR) cells | Pujol et al. 2022 | N/A |
| HeLa FITR EGFP-ORF9b$^{WT}$ (AHT-inducible) | This study | N/A |
| HeLa FITR EGFP-ORF9b$^{L52D}$ (AHT-inducible) | This study | N/A |
| HeLa FITR ORF9b$^{WT}$-FLAG and ORF9b$^{s53E}$-FLAG (single-copy, AHT-inducible) | This study | N/A |
| HeLa FITR EGFP-ORF9b (SARS-CoV-1, BANAL-236, RaTG13 variants) | This study | N/A |
| HeLa FITR TOMM70-deficient clones (*TOMM70$^{C11}$*, *TOMM70$^{C14}$*, *TOMM70$^{C15}$*) | This study | N/A |
| HeLa FITR *TOMM70$^{C11}$*: pLV-TOMM70$^{WT}$ | This study | N/A |
| HeLa FITR *TOMM70$^{C11}$*: pLV-TOMM70$^{E477A}$ | This study | N/A |
| HeLa FITR *TOMM70$^{C11}$*: pLV-TOMM70$^{D545A}$ | | |
| HeLa FITR cells stably expressing iMLS-QC mitophagy reporter | This study | N/A |
| HeLa FITR cells stably expressing IRF3-EGFP | This study | N/A |
| HeLa FITR ORF9b$^{WT}$-FLAG cells stably expressing C15ORF48 (pCDH hC15orf48-ORF) | This study | N/A |
| A549$^{+hACE2+hTMPSS2}$ cells (stably expressing human ACE2 and TMPRSS2) | This study | N/A |
| A549 cells stably expressing ORF9b$^{WT}$ (pLV-ORF9b$^{WT}$) | This study | N/A |
| Toxoplasma gondii type I RH Δku80, cytosolic NanoLuc-GFP | Gift from A. Bougdour, Univ. Grenoble-Alpes | N/A |
| Toxoplasma gondii type I RH Δku80, GRA17-HA | Gift from A. Bougdour, Univ. Grenoble-Alpes | N/A |
| Sendai virus (SeV) | American Type Culture Collection | VR-105 |

| | | |
|---|---|---|
| SARS-CoV-2 (Wuhan/wild-type strain: BetaCoV/France/IDF00372/2020) | de Melo et al. 2021 | EVAg collection, Ref-SKU: 014V-03890) |
| SARS-CoV-2$^{\Delta ORF9b}$ (ORF9b-deleted, recombinant) | This study | N/A |
| SARS-CoV-2$^{ORF9bL52D}$ (ORF9b L52D, recombinant) | This study | N/A |
| Male Syrian hamsters (Mesocricetus auratus), 5–6 weeks old | N/A | N/A |
| **Antibodies** | | |
| Anti-TOMM70 (rabbit) | Proteintech | 66593-1-IG |
| Anti-TOMM20 | Proteintech | 66777-1-Ig |
| Anti-TOMM40 | Proteintech | 18409-1-AP |
| Anti-NDUFB7 (Complex I) | Proteintech | 14912-1-AP |
| Anti-ORF9b (SARS-CoV-2) | Abcam | ab308403 (clone HL1918) |
| Anti-C15ORF48 (NMES1) | Proteintech | 25102-1-AP |
| Anti-NDUFA4 | Clinisciences | BS3883 |
| Anti-FLAG | Sigma-Aldrich | F3165 (M2) |
| Anti-Nucleocapsid (SARS-CoV-2) | Fisher Scientific | 16363694 |
| Alexa Fluor 488/568/647-conjugated secondary antibodies | Thermo Fisher Scientific (Invitrogen) | host/fluorophore-specific (e.g., A-21140) |
| HRP-conjugated secondary antibodies (anti-rabbit/anti-mouse) | Euromedex (Bethyl) | A120-101P (goat anti-rabbit-HRP) |
| **Recombinant DNA** | | |
| pcDNA5/FRT/TO EGFP-ORF9b$^{WT}$ (SARS-CoV-2, Wuhan) | Addgene | #165122 |
| pcDNA5/FRT/TO EGFP-ORF9b$^{L52D}$ | Addgene | #165129 |
| pDONR223-SARS-CoV-2_ORF9b | Addgene | #141280 |
| pDONR221 (Gateway entry vector) | Invitrogen | N/A |
| pcDNA5/FRT/TO (Gateway-compatible, tetracycline-inducible destination vector) | Pujol et al. 2022 | N/A |
| pOG44 (Flp recombinase expression plasmid) | Invitrogen | N/A |
| EGFP-ORF9b entry/destination clones (SARS-CoV-1, BANAL-236, RaTG13) | This study | N/A |
| pDONR223-ORF9b$^{WT}$-FLAG entry clone | This study | N/A |
| pDONR223-ORF9b$^{S53E}$-FLAG entry clone | This study | N/A |

| | | |
|---|---|---|
| Codon-optimized human TOMM70 cDNA | Eurofins | N/A |
| pDONR221-TOMM70 (wild-type entry clone) | This study | N/A |
| pDONR221-TOMM70$^{E477A}$ entry clone | This study | N/A |
| pDONR221-TOMM70$^{D545A}$ entry clone | This study | N/A |
| pLV_Destination (Gateway-compatible lentiviral destination vector, CMV promoter) | N/A | N/A |
| pLV-TOMM70$^{WT}$ | This study | N/A |
| pLV-TOMM70$^{E477A}$ | This study | N/A |
| pLV-TOMM70$^{D545A}$ | This study | N/A |
| lentiCRISPR-sgTOMM70 (sgRNA targeting human TOMM70) | Gift from Prof. Lena Pernas | N/A |
| pCMV-delta R8.2 (lentiviral packaging plasmid) | Addgene | #12263 |
| pCMV-VSV-G (lentiviral envelope plasmid) | Addgene | #8454 |
| pLenti-III-PGK-Zeo-iMLS (mCherry-GFP iMLS mitophagy reporter) | Gift from Anne Simonsen | N/A |
| pCDH hC15orf48-ORF construct | Gift from Lena Ho | N/A |
| **Oligonucleotides and other sequence-based reagents** | | |
| **Gene (Primer ID)** | **Sequence (5′→3′)** | **Purpose** |
| SARS-CoV-2 ORF9b N-terminal FLAG, forward | GATGACGACAAGTAATACCCAACTTTCTTGTACAAAG | Site-directed mutagenesis: insert N-terminal FLAG tag into ORF9b$^{WT}$ (with reverse primer) |
| SARS-CoV-2 ORF9b N-terminal FLAG, reverse | GTCTTTGTAGTCCTTCACGGTCACCACCAC | |
| SARS-CoV-2 ORF9b S53E, forward | CAGCCCTCTGGAACTGAACATGG | Site-directed mutagenesis: ORF9b$^{WT}$-FLAG → ORF9b$^{S53E}$-FLAG (with reverse primer) |
| SARS-CoV-2 ORF9b S53E, reverse | CCCAGTCTCAGGATGATAGG | |
| *TOMM70* E477A, forward | GGGATATGGATATGCCCTGTATGCC | Site-directed mutagenesis: TOMM70 E477A (with reverse primer) |
| *TOMM70* E477A, reverse | GCGGCACACGGCACATCGGGGAAATT | |
| *TOMM70* D545A, forward | TGCCTTCTTCGCCTATGAGACAATG | Site-directed mutagenesis: TOMM70 D545A (with reverse primer) |
| *TOMM70* D545A, reverse | CACTTATTCACACTTATTGTCGATTTC | |

| | | |
|---|---|---|
| *TOMM70* Illumina adapted genotyping, forward | TCGTCGGCAGCGTCAGATGTGTATAAGAGACAGCGGGTGCCATATACCTGTG | Genotyping: deep NGS of CRISPR-targeted TOMM70 locus, flanking sgRNA cut site |
| *TOMM70* Illumina adapted genotyping, reverse | GTCTCGTGGGCTCGGAGATGTGTATAAGAGACAGCAAATGCCCCACTCCCATCT | |
| Beta Actin, forward | CATGTACGTTGCTATCCAGGC | Human RT-qPCR normalization |
| Beta Actin, forward | CTCCTTAATGTCACGCACGAT | |
| *MTARC2* (Human), forward | GGAGTCATAGACAGGAAACAGCC | RT-qPCR quantification of MTARC2 expression |
| *MTARC2* (Human), reverse | GTCACCAACTCTCAGGCTTCCA | |
| *TOMM70* (Human), forward | TGTTTTGCATTGTACCGCCAG | RT-qPCR quantification of TOMM70 expression |
| *TOMM70* (Human), reverse | TAGTGCATAGCCTTCGGCAC | |
| *MAVS* (Human), forward | ATGGTGCTCACCAAGGTGTCTG | RT-qPCR quantification of MAVS expression |
| *MAVS* (Human), reverse | TCTCAGAGCTGCTGTCTAGCCA | |
| *C15ORF48* (Human), forward | AGGAAGGAACTCATTCCCTTGG | RT-qPCR quantification of C15orf48 (COXFA4L3 / NMES1 / MOCCI) expression |
| *C15ORF48* (Human), reverse | TTTTGAGGTACAGTAGGGTCCA | |
| CUNH15orf48 (Hamster), forward | ACTGGAGCCTCATCTTTTGCT | RT-qPCR quantification of C15orf48 (COXFA4L3 / NMES1 / MOCCI) expression |
| CUNH15orf48 (Hamster), reverse | CTTCAACGGGCTTCCATTGC | |
| b-actin (Hamster), forward | GGCCAGGTCATCACCATT | RT-qPCR normalization (hamster b-actin) |
| b-actin (Hamster), reverse | GAGTTGAATGTAGTTTCGTGGATG | |
| **Chemicals, Peptides and Recombinant Proteins** | | |
| **Cell culture and selection** | | |
| DMEM 1X + GlutaMAX™ (4.5 g/L D-glucose, sodium pyruvate) | Thermo Fisher Scientific | 10569010 |
| Ham's F-12K (Kaighn's) Medium | Thermo Fisher Scientific | 21127030 |
| Glucose-free DMEM | Thermo Fisher Scientific | 11966025 |
| Fetal Bovine Serum (FBS) | Thermo Fisher Scientific | 10270106 |

| | | |
|---|---|---|
| Penicillin–Streptomycin | Thermo Fisher Scientific | P4333 |
| D-(+)-Galactose | Euromedex | 1042-C |
| Anhydrotetracycline (AHT) | IBA Lifesciences | 2-0401-002 |
| Hygromycin B | Invivogen | ant-hg-1 |
| Blasticidin | Invivogen | ant-bl-1 |
| Puromycin | Invivogen | ant-pr-1 |
| Zeocin | Invivogen | ant-zn-1 |
| Lipofectamine 2000 | Invitrogen | 11668019 |
| Lipofectamine RNAiMAX | Invitrogen | 13778150 |
| Q5 Site-Directed Mutagenesis Kit | New England Biolabs | E0554S |
| HEPES buffer (pH 7.2) | Thermo Fisher Scientific | 15630080 |
| GlutaMAX | Thermo Fisher Scientific | 41090028 |
| Poly-L-lysine | Sigma-Aldrich | P8920 |
| **siRNA** | | |
| Non-targeting control siRNA | Dharmacon (Horizon Discovery) | D-001220-01-05 |
| siRNA targeting human TOMM70 | Dharmacon (Horizon Discovery) | L-054815-01-0005 |
| siRNA targeting human TOMM40 | Dharmacon (Horizon Discovery) | M-012732-00-0005 |
| siRNA targeting human MAVS | Dharmacon (Horizon Discovery) | M-024237-02-0005 |
| siRNA targeting human MARC2 | Dharmacon (Horizon Discovery) | M-018689-00-0005 |
| **Proteasome inhibitors and cell death/mitophagy reagents** | | |
| MG132 (1 μM) | Sigma-Aldrich | M7449 |
| Bortezomib (0.5 μM) | Sigma-Aldrich | 5043140001 |
| DMSO (vehicle control) | Euromedex | UD8050-05-A |
| ABT-737 (10 μM) | Clinisciences | A8193 |
| Actinomycin D (1 μM) | Sigma-Aldrich | SBR00013 |
| Staurosporine (STS, 0.5 μM) | Sigma-Aldrich | S6942 |
| Etoposide (100 μM) | Sigma-Aldrich | E1383 |
| 3-Hydroxy-1,2-dimethyl-4(1*H*)-pyridone (DFP) | Sigma-Aldrich | 379409 |
| **Imaging reagents** | | |
| NucBlue™ Live ReadyProbes™ Reagent | Thermo Fisher Scientific | R37605 |
| MitoTracker™ Deep Red FM (MTDR) | Thermo Fisher Scientific | M22426 |
| Tetramethylrhodamine Ethyl Ester Perchlorate (TMRE) | Sigma-Aldrich | 87917 |
| Propidium Iodide (PI) | Sigma-Aldrich | P4170 |

| | | |
|---|---|---|
| Hoechst 33342 | N/A | H3570 |
| CellCarrier Ultra imaging plates (96-well) | PerkinElmer | 6055300 |
| Acryloyl-X, SE | N/A | A20770 |
| Proteinase K | New England Biolabs | P8007S |
| **Fixation, permeabilization and general buffers** | | |
| Paraformaldehyde (PFA), 4% | N/A | 15714 |
| Triton X-100 | Euromedex | 2000-C |
| Tween-20 | Euromedex | 2001-C |
| PBS | Thermo Fisher Scientific | N/A |
| RIPA buffer components (Tris-HCl, NaCl, Triton X-100, SDS, sodium deoxycholate, EDTA) | N/A | N/A |
| cOmplete EDTA-free Protease Inhibitor Cocktail | Roche | 4693159001 |
| Buffer 17 (HEPES, potassium acetate, glycerol, 6-aminohexanoic acid, EDTA) | N/A | N/A |
| Trehalose / freezing buffer components | N/A | N/A |
| Bradford Assay Reagent | Sigma-Aldrich | 10495315 |
| **SDS-PAGE, BN-PAGE and immunoblotting** | | |
| 4–20% Mini-PROTEAN® TGX Stain-Free™ precast gels | Bio-Rad | 4568096 |
| 6x Laemmli Sample Buffer with 2-mercaptoethanol | N/A | N/A |
| Non-fat dry milk | N/A | 711160 |
| Clarity™ Western ECL Substrate | Bio-Rad | 1705061 |
| Digitonin (ultra-pure) | N/A | D141-500MG |
| NativePAGE™ 5% G-250 Sample Additive | Thermo Fisher Scientific | BN2004 |
| NativePAGE™ 3–12% Bis-Tris gels | Thermo Fisher Scientific | BN1001BOX |
| PVDF membranes | N/A | 1704272 |
| Acetic acid / methanol (BN-PAGE membrane fixation/destaining) | N/A | A6283-1L (acetic acid); 179337-1L (methanol) |
| **RNA extraction, RT-qPCR and proteomics sample prep** | | |
| TRIzol™ Reagent | Thermo Fisher Scientific | 15596026 |
| Chloroform | N/A | C2432-1L |
| TCEP (tris(2-carboxyethyl)phosphine) | N/A | T2556 |
| Iodoacetamide | N/A | GERPN6302 |

| | | |
|---|---|---|
| Sera-Mag Carboxylate-Modified SpeedBeads, hydrophobic | Cytiva | 65152105050250 |
| Sera-Mag Carboxylate-Modified SpeedBeads, hydrophilic | Cytiva | 45152105050250 |
| Sequencing-grade modified Trypsin | N/A | V5111 |
| Ammonium bicarbonate | N/A | N/A |
| Formic acid (FA) | N/A | N/A |
| Acetonitrile (ACN, LC-MS grade) | N/A | N/A |
| Trifluoroacetic acid (TFA) | N/A | N/A |
| **Plaque assay and viral culture reagents** | | |
| MEM 10X | Gibco™ | 21430020 |
| Distilled water (cell-culture grade) | Gibco™ | 15230162 |
| L-Glutamine | Gibco™ | 25030123 |
| Gentamicin (10 mg/mL) | Gibco™ | 15710064 |
| Sodium Phosphate, 0.25% | Gibco™ | 25080094 |
| AVICEL (microcrystalline cellulose, overlay medium) | DuPont | RC-581 |
| Crystal Violet | Merck | V5265 |
| Formaldehyde, 4% (tissue fixation) | Pasteur Thermo Fisher Scientific | 10532955 |
| **Animal procedures** | | |
| Ketamine (anesthesia) | N/A | N/A |
| Xylazine (anesthesia) | N/A | N/A |
| Physiological saline | N/A | N/A |
| **Critical Commercial Assays** | | |
| NucleoSpin RNA Kit | Macherey-Nagel | 740955 |
| RNeasy Kit | Qiagen | 74104 |
| Direct-zol RNA MiniPrep w/ Zymo-Spin IIC Columns (hamster RNA) | Zymo Research / OZYME | R2050 |
| iScript Reverse Transcription Supermix | Bio-Rad | 1708891 |
| SYBR® Green Master Mix | Bio-Rad | 1725125 |
| TruSeq Stranded mRNA Library Preparation Kit | Illumina | 20020594 |
| nCounter XT Human Host Response CodeSet (Panel XT Hs Host Response CSO) | NanoString Technologies | 115000449 |
| nCounter XT Master Kit (12 reactions) | NanoString Technologies | 100052 |
| AssayMAP C18 cartridges (5 µL bed volume) | Agilent Technologies | 5190-6532 |
| **Deposited Data** | | |

| | | |
|---|---|---|
| HeLa FITR cell proteomics dataset | This study | PRIDE: PXD074755 |
| A549 cell proteomics dataset | This study | PRIDE: PXD074770 |
| A549 cell bulk RNAseq dataset | This study | ENA: E-MTAB-16841 |
| Hamster lung bulk RNAseq dataset | This study | ENA: E-MTAB-16842 |
| SARS-CoV-2 Wuhan reference genome (for RNAseq mapping) | NCBI | GCF_009858895.2 |
| Human TOM70–SARS-CoV-2 ORF9b complex structure | Gao et al. 2021 | PDB: 7DHG |
| **Software and Algorithms** | | |
| PyMOL | Schrödinger, LLC | pymol.org |
| AlphaFold 3.0 | DeepMind / EMBL-EBI | alphafold.ebi.ac.uk |
| Fiji / ImageJ | NIH | fiji.sc |
| DeconvolutionLab2 (Fiji plugin) | EPFL Biomedical Imaging Group | github.com/Biomedical-Imaging-Group/DeconvolutionLab2 |
| 3D Viewer (ImageJ plugin) | N/A | imagej.net/plugins/3d-viewer |
| Custom mitochondrial morphology macro (MitoShape) | This study | github.com/jdhercam10/MitoShape |
| MetaMorph software | Molecular Devices | RRID:SCR_002368 |
| Harmony 5.1 (high-content imaging analysis) | PerkinElmer | RRID:SCR_018809 |
| PhenoLOGIC™ supervised machine-learning module | PerkinElmer | N/A |
| GraphPad Prism v10 | GraphPad Software | graphpad.com |
| Sequana v0.18.1 / RNA-seq pipeline v0.20.0 | Cokelaer et al. 2017 | github.com/sequana/sequana_rnaseq |
| Snakemake v7.32.4 | Köster & Rahmann 2012 | snakemake.github.io |
| Fastp v0.23.2 | Chen et al. 2018 | github.com/OpenGene/fastp |
| STAR v2.7.10a | Dobin et al. 2013 | github.com/alexdobin/STAR |
| FeatureCounts v2.0.1 | Liao et al. 2014 | subread.sourceforge.net |
| MultiQC v1.16.0 | Ewels et al. 2016 | multiqc.info |
| Bowtie 2 (v2.4.5; SARS-CoV-2 read mapping) | N/A | RRID:SCR_016368 |

| | | |
|---|---|---|
| DESeq2 v1.38.3 (R/Bioconductor) | Love et al. 2014 | bioconductor.org/packages/DESeq2 |
| limma v3.66.0 (R/Bioconductor; voom, moderated t-test) | Ritchie et al. 2015 | bioconductor.org/packages/limma |
| imp4p (R package; missing value imputation) | Gianetto et al. 2020 | cran.r-project.org/package=imp4p |
| cp4p (R package; adaptive Benjamini-Hochberg FDR) | Giai Gianetto et al. 2016 | cran.r-project.org/package=cp4p |
| WGCNA (R package; co-expression module detection) | Langfelder & Horvath | 10.1186/1471-2105-9-559 |
| ComplexHeatmap (R package) | Gu et al. 2016 | bioconductor.org/packages/ComplexHeatmap |
| g:Profiler | Reimand et al. 2019 | biit.cs.ut.ee/gprofiler |
| Enrichr | Chen et al. 2013 | maayanlab.cloud/Enrichr |
| GSEA (Gene Set Enrichment Analysis) | Subramanian et al. 2005 | gsea-msigdb.org |
| WebGestalt (over-representation analysis) | N/A | webgestalt.org |
| Spectronaut v20.2.250922.92449 (directDIA / Pulsar Search) | Biognosys | biognosys.com/software/spectronaut |
| IDPicker algorithm (protein inference) | N/A | proteowizard.sourceforge.io/idpicker |
| nSolver Analysis Software | NanoString Technologies | RRID:SCR_003420 |
| QuPath (digital pathology / fibrosis quantification) | Bankhead et al. 2017 | qupath.github.io |
| Image Lab Software (densitometry) | Bio-Rad | RRID:SCR_014210 |
| IncuCyte confluence analysis software | Sartorius | RRID:SCR_023147 |
| BioRender (figure preparation) | BioRender | biorender.com |
| Benchling (figure/sequence preparation) | Benchling | benchling.com |
| Adobe Illustrator (figure preparation) | Adobe Inc. | adobe.com |
| R / prcomp (PCA, statistical computing) | R Core Team | r-project.org |
| **Other (Equipment)** | | |

| | | |
|---|---|---|
| Operetta CLS High-Content Imaging System (20x Water/1.0 NA) | PerkinElmer | HH16000020 |
| Opera Phenix Plus High-Content Screening System (63x Water/1.15 NA) | PerkinElmer | HH14001000 |
| IncuCyte SX5 Live-Cell Analysis System | Sartorius | N/A |
| Seahorse XFe96 Analyzer | Agilent (Seahorse Biosciences) | S7800A |
| Infinite M2000 microplate reader | TECAN | N/A |
| NanoQuant Plate™ (Infinite M200) | TECAN | N/A |
| ChemiDoc® Imaging System | Bio-Rad | 12003153 |
| XCell SureLock™ Mini-Cell electrophoresis system | Thermo Fisher Scientific | EI0001 |
| Trans-Blot® Turbo™ Transfer System | Bio-Rad | 1704150 |
| CFX384 Touch Real-Time PCR Detection System | Bio-Rad | 1855485 |
| nCounter Analysis System | NanoString Technologies | N/A |
| Eclipse Ti inverted confocal microscope (CSU-X1 spinning disk) | Nikon / Yokogawa | N/A |
| sCMOS Prime camera | Photometrics | N/A |
| Chamlide live-cell imaging chambers | LCI Corp. | N/A |
| AxioScan.Z1 Automated Slide Scanner | Zeiss | 430038-9001-000 |
| Covaris E220 focused-ultrasonicator | Covaris | 500239 |
| microTUBE-15 AFA Beads Screw-Cap tubes | Covaris | 520145 |
| Bravo automated liquid-handling platform | Agilent Technologies | G5409A |
| NanoElute 2 UHPLC system | Bruker Daltonics | N/A |
| timsTOF Ultra 2 mass spectrometer (DIA-PASEF) | Bruker Daltonics | N/A |
| PepSep Ultra C18 analytical column (25 cm × 75 μm, 1.5 μm) | Bruker Daltonics | 1893484 |
| Agilent BioAnalyzer | Agilent Technologies | G2939BA |
| Illumina NextSeq 2000 | Illumina | 20038897 |
| Glass Potter-Elvehjem homogenizer | N/A | 10299651 |
| Biopsy punch (4 mm, expansion microscopy) | N/A | N/A |
